# Microbial-associated acylated putrescines as immunomodulatory molecules in inflammatory bowel diseases

**DOI:** 10.64898/2026.08.24.746719

**Authors:** Sena Bae, Julian Avila-Pacheco, Slater L. Clay, Sunghee Bang, Milcah C. Scott, Natalia Andreeva, Monia Michaud, Diogo Fonseca-Pereira, Eunyoung Chun, Y. Grace Cao, Yancong Zhang, Amrisha Bhosle, Rosa M. Perez, Gleb Pishchany, Hera Vlamakis, Geniver El Tekle, Xochitl C. Morgan, Sway P. Chen, Jonathan N. Glickman, Ramnik J. Xavier, Daniel B. Graham, Clary B. Clish, Jon Clardy, Eric A. Franzosa, Curtis Huttenhower, Wendy S. Garrett

## Abstract

The gut microbiota profoundly shapes intestinal immunity through the production of small-molecule metabolites; yet in inflammatory bowel diseases (IBD), where the microbial metabolome is substantially altered, the identities and immunological functions of most disease-associated metabolites remain unknown. Here, we integrate untargeted fecal metabolomics from two independent IBD patient cohorts with gnotobiotic mouse metabolomic data to identify N-acyl putrescines as a class of microbiome-associated metabolites consistently enriched in both Crohn’s disease (CD) and ulcerative colitis (UC). Among these, N-oleoylputrescine (NOP) induces potent and selective transcriptional responses in bone marrow-derived dendritic cells (BMDCs) and colonic organoids, establishing it as the primary immunomodulatory candidate in this metabolite class. *Enterocloster* species harboring nonribosomal peptide synthetase (NRPS) biosynthetic gene clusters produce NOP via conjugation of oleic acid with putrescine, confirmed by isotope-tracing in vitro and germ-free mouse mono-colonization in vivo. NOP suppresses five core IBD-associated inflammatory pathways in mouse dendritic cells and human monocytes, reduces gene signatures of histologic inflammation and IBD therapy non-response, and ameliorates colitis in four murine models. NOP dampens NF-κB activation and iNOS expression in myeloid cells and suppresses type 1 immune responses through a T cell-intrinsic mechanism. The enrichment of NOP in IBD despite its anti-inflammatory activity supports a holobiont defense hypothesis: that the gut microbiota mounts a compensatory metabolic response to intestinal inflammation that may contribute to the restoration of organismal homeostasis.

**In Brief:** Bae et al. identify N-acyl putrescines as gut microbiota-associated metabolites enriched in IBD and demonstrate that N-oleoylputrescine (NOP), produced by *Enterocloster* species via NRPS biosynthetic machinery, broadly suppresses innate and adaptive inflammatory programs and ameliorates colitis in mice, supporting a holobiont defense hypothesis for microbiota-mediated immunomodulation.

**Highlights:**

- N-acyl putrescines are microbiome-associated metabolites enriched in IBD feces across two independent human cohorts
- *Enterocloster* species harboring NRPS biosynthetic gene clusters produce NOP via oleic acid– putrescine conjugation
- NOP suppresses NF-κB, iNOS, and IBD therapy non-response gene signatures in mouse and human myeloid cells
- NOP ameliorates colitis in four distinct murine models and suppresses type 1 immunity via a T cell-intrinsic mechanism

## INTRODUCTION

The gut microbiota exerts profound influence over host immunity through the production of small-molecule metabolites that serve as chemical mediators of host-microbiome communication.^1–3^ These microbially derived molecules traverse the intestinal epithelial barrier, accumulate in mucosal tissue, and in some cases enter systemic circulation, enabling them to modulate immune cell function at local and distal sites.^4,5^ The catalog of microbiota-derived immunomodulatory metabolites has expanded considerably over the past decade, encompassing short-chain fatty acids, secondary bile acids, tryptophan catabolites, and N-acyl amino acids, among others.^1,3,6^ Nevertheless, untargeted metabolomics studies consistently reveal that potentially biologically active features in the gut metabolome remain structurally and functionally uncharacterized, indicating that important immunomodulatory molecules await to be uncovered.^2,7^

Inflammatory bowel diseases (IBD), encompassing Crohn’s disease (CD) and ulcerative colitis (UC), are chronic immune-mediated disorder characterized by dysregulated intestinal inflammation that affects millions of individuals worldwide and remain incompletely controlled by available therapies.^6^ IBD is associated with dysbiosis, disruption of normal gut microbial community structure and function, accompanied by sweeping alterations in the intestinal metabolome.^8,9^ Multi-omics studies have catalogued hundreds of metabolic features differentially abundant in IBD patients compared to healthy controls, yet for the vast majority the molecular identities and immunological activities remain unknown.^9,10^ This gap is consequential: understanding which disease-enriched metabolites are produced by the microbiota and what effects they exert on host immunity could illuminate new mechanisms of IBD pathogenesis and identify candidate molecules for therapeutic development.

Polyamines represent a particularly compelling class of metabolites in this context. Putrescine, spermidine and spermine, the most abundant dietary and endogenous polyamines, have established roles in macrophage polarization and efferocytosis, T cell differentiation and lineage fidelity, and effector function, all of which are central to IBD pathogenesis.^11–18^ We found that putrescine and N-acetylputrescine are among the most significantly enriched metabolites in IBD feces across independent patient cohorts, raising the question of whether acylated derivatives of putrescine, products of microbiota-mediated conjugation of putrescine with fatty acids, might represent a broader class of immunomodulatory molecules in the inflamed gut. Notably, the gut microbiota encodes a rich repertoire of nonribosomal peptide synthetase (NRPS) biosynthetic gene clusters (BGCs) capable of mediating fatty acid amide conjugation reactions, and gut-inhabiting Clostridia have been shown to produce N-acyl amide GPCR ligands through analogous enzymatic chemistry, ^19,20^ suggesting that the chemical space of microbiota-derived N-acyl conjugates is likely far broader than currently appreciated.

The identification and functional characterization of disease-enriched, microbiota-derived metabolites is technically challenging given the large number of differentially abundant metabolic features detected in untargeted LC-MS datasets and the scarcity of structural annotations. Computational prioritization tools, such as MACARRoN (Metabolome Analysis and Combined Annotation Ranks to pRioritize Novel bioactives),^7^ which prioritizes novel bioactive metabolites by leveraging co-abundance with known metabolites and differential abundance across phenotypes, provide a systematic framework for narrowing the discovery space. We cross-referenced disease-associated metabolic features in human datasets with matched features in gnotobiotic mouse metabolomes, where host genetics, diet, and microbial community composition are precisely controlled, as a strategy to identify features that are both IBD-enriched and of microbial origin. Applying this integrated approach to the Human Microbiome Project 2 (HMP2/iHMP) fecal metabolomic,^9^ with validation in the independent PRISM cohort^8^ and confirmation of microbial origin using the FARMM human antibiotic intervention cohort,^21^ identified a polyamine-anchored metabolic module as a top-priority candidate and revealed an acylated putrescine enriched in IBD that is anti-inflammatory.

Here, we report the discovery, chemical characterization, microbial biosynthesis, and immunological activities of N-acyl putrescines as a family of microbiome-associated metabolites enriched in the IBD gut. We demonstrate that the leading member of this class, N-oleoylputrescine (NOP), is produced by *Enterocloster* species via NRPS-mediated conjugation of oleic acid with putrescine, potently suppresses five core IBD-associated inflammatory pathways in mouse and human myeloid cells, reduces transcriptional signatures of histologic inflammation and IBD therapy non-response, and ameliorates colitis in four distinct mouse models by dampening innate immune activation and suppressing type 1 T cell responses through a cell-intrinsic mechanism. The paradoxical enrichment of this anti-inflammatory metabolite in patients with IBD supports a holobiont defense hypothesis, that the gut microbiota mounts a compensatory metabolic response to intestinal inflammation that may be insufficient to resolve established disease but may nonetheless limit injury and contribute to homeostatic restoration. These findings expand our understanding of host-microbiome chemical signaling in intestinal inflammation and identify NOP as a candidate for therapeutic development in IBD.

## RESULTS

### Identification of acylated putrescines as IBD-enriched, microbiome-associated metabolites

Analysis of 546 fecal metabolomes from the Integrative Human Microbiome Project (HMP2/iHMP)^9^ revealed thousands of metabolic features that are differentially abundant in IBD and of likely microbial origin. To identify which of these were both disease-relevant and microbiota-associated, we integrated human IBD-associated metabolic features prioritized by MACARRoN^7^ with microbial association data derived from gnotobiotic mice harboring defined microbial communities, germ-free (GF), minimal microbiota (altered Schaedler flora, ASF), and complex microbiota (specific pathogen-free, SPF), in which host genetics, diet and environment are controlled, aligning 10,668 matched metabolic features across datasets (Figure 1A; STAR Methods). Among 13,787 total differentially abundant features in IBD, this approach identified 956 microbe-associated features differentially abundant in CD and 2,019 in UC, with approximately 79% of all IBD-associated features confirmed as microbe-associated (Figure 1B–C; Figure S1A; Tables S1-2). Notably, a parallel analysis using the FARMM human antibiotic-depletion cohort identified a comparable number of microbe-associated metabolic features (n=2,793) to the gnotobiotic filtering approach (n=2,891), supporting the cross-species validity of the microbiota-association strategy employed here^21^ (Table S3).

**Figure 1.**
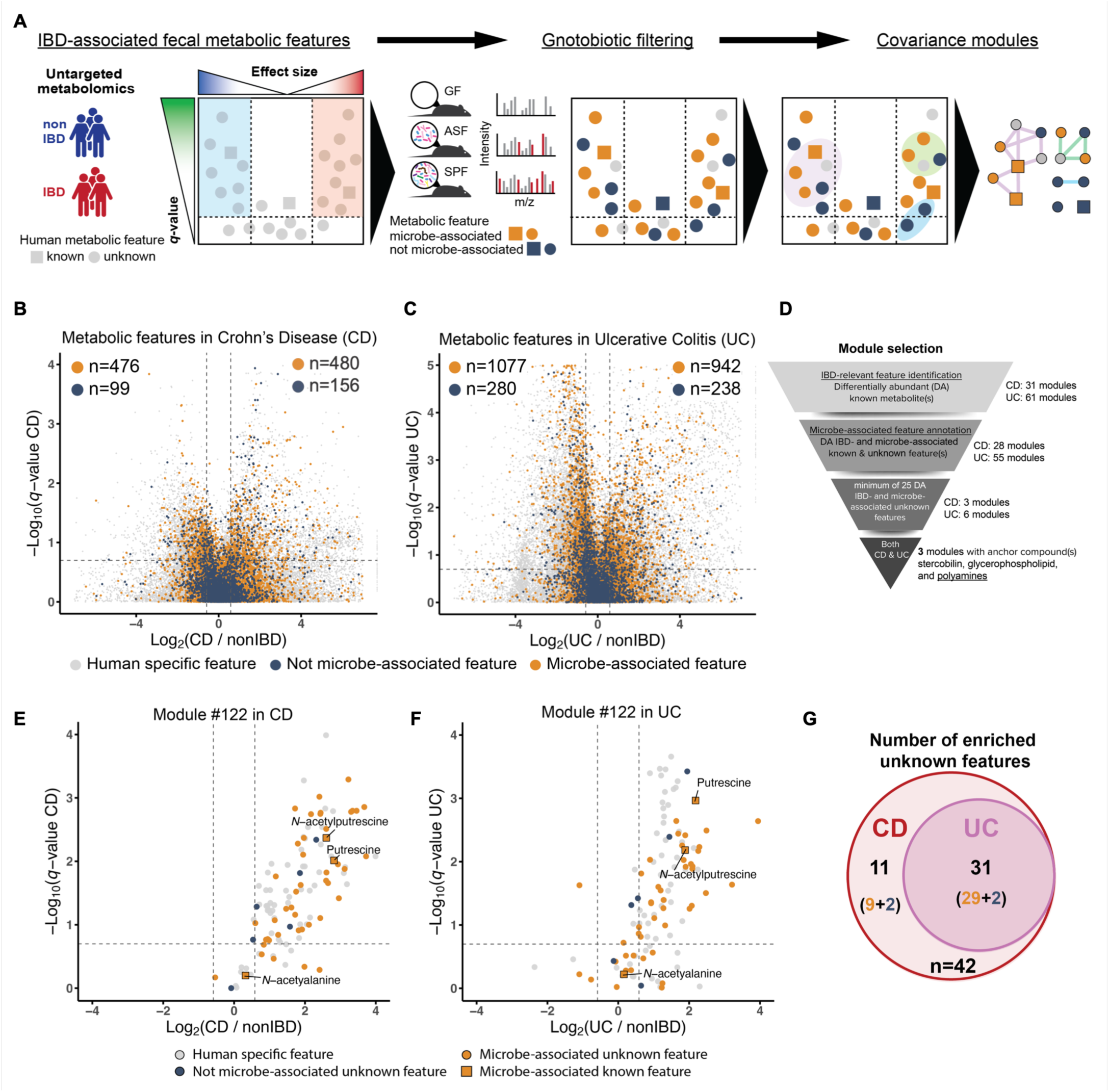
Prioritization of IBD- and microbe-associated metabolic features. (A) Prioritization schematic. (B–C) Volcano plots depicting q-value (FDR-adjusted P value, y-axis) from a linear mixed model analysis and log₂ fold change (x-axis) between CD and non-IBD samples (B), and between UC and non-IBD samples (C) from 546 HMP2 fecal untargeted metabolomes; differentially abundant (DA) features in IBD were defined as |log₂ fold change| > log₂(1.5) and FDR-adjusted P < 0.2 (linear mixed model; STAR Methods). Gray dots represent metabolic features only detected in HMP2. Orange and navy blue dots represent features detected in both HMP2 and gnotobiotic mouse fecal metabolomic datasets, as microbe-associated (orange) and non-microbe-associated (navy blue), respectively. (D) Disease-relevant and microbe-associated module selection funnel analysis. Prioritized modules contained at least one DA known metabolite and 25 IBD- and microbe-associated unknown metabolic features meeting these criteria across both IBD phenotypes. (E–F) Volcano plots depicting metabolic features in Module #122 (n=115) selected for investigation in CD (E) and UC (F) samples. (G) Venn diagram of Module #122 depicting features selected using the prioritization scheme (A). Thirty-one unknown features were enriched in both CD and UC; 29 were microbe-associated (orange) and two were non-microbe-associated (navy blue). Eleven were enriched only in CD.

Among the disease-relevant, microbiota-associated metabolic features, those with correlated abundance patterns, suggestive of shared biosynthetic origins, were grouped into modules anchored by known biologically active metabolites (Figure 1D; STAR Methods). Three modules met stringent criteria for biological relevance in both IBD phenotypes, anchored by stercobilin (module #11, depleted in IBD), glycerophospholipids (module #44, enriched in IBD), and polyamines (module #122, enriched in IBD), respectively. Module #122, anchored by putrescine and N-acetylputrescine, both significantly enriched in IBD (Figure S1B), stood out as the highest-priority candidate. Polyamines, and putrescine in particular, regulate macrophage polarization, T cell differentiation, lineage fidelity, and effector function,^11–18^ all of which are central to IBD pathogenesis. In addition to its known polyamine anchors, module #122 contained 29 uncharacterized metabolic features that are microbe-associated and enriched in both CD and UC, and eleven further features enriched only in CD (Figure 1E–G; Table S4). Twelve of these candidate features were independently replicated in an external IBD cohort (PRISM; Figure S1C; Table S5),^8^ supporting their robust association with IBD across patient populations.

### Chemical identification and IBD enrichment of acylated putrescines

Analysis of module #122 candidate features by tandem mass spectrometry followed by in silico m/z prediction (STAR Methods) led to identification of five previously uncharacterized acylated putrescines: diacetylputrescine (DAP), N-propionoylputrescine (NPP), N-valeroyl/N-isovaleroylputrescine (NVP/N-isoVP), N-oleoylputrescine (NOP), and N-stearoylputrescine (NSP). To definitively establish their chemical identities, we synthesized authentic reference standards for NPP, NOP, NSP and NVP, compounds not commercially available, and confirmed their structures by matching retention time, precursor m/z, and product ion spectra in HMP2, PRISM, and mouse fecal metabolomes (Figure 2A; Figures S2A–G and S5A–B; Table S6). We additionally synthesized two acylated putrescines not included in module #122 to enable direct comparison of a range of acylated putrescine bioactivities: N-butyroylputrescine (NBP), detected exclusively in the gnotobiotic dataset, and N-myristoylputrescine (NMP), a recently reported microbiota-produced acylated putrescine enriched in the IBD gut and implicated in disrupting intestinal barrier integrity.^10^

**Figure 2.**
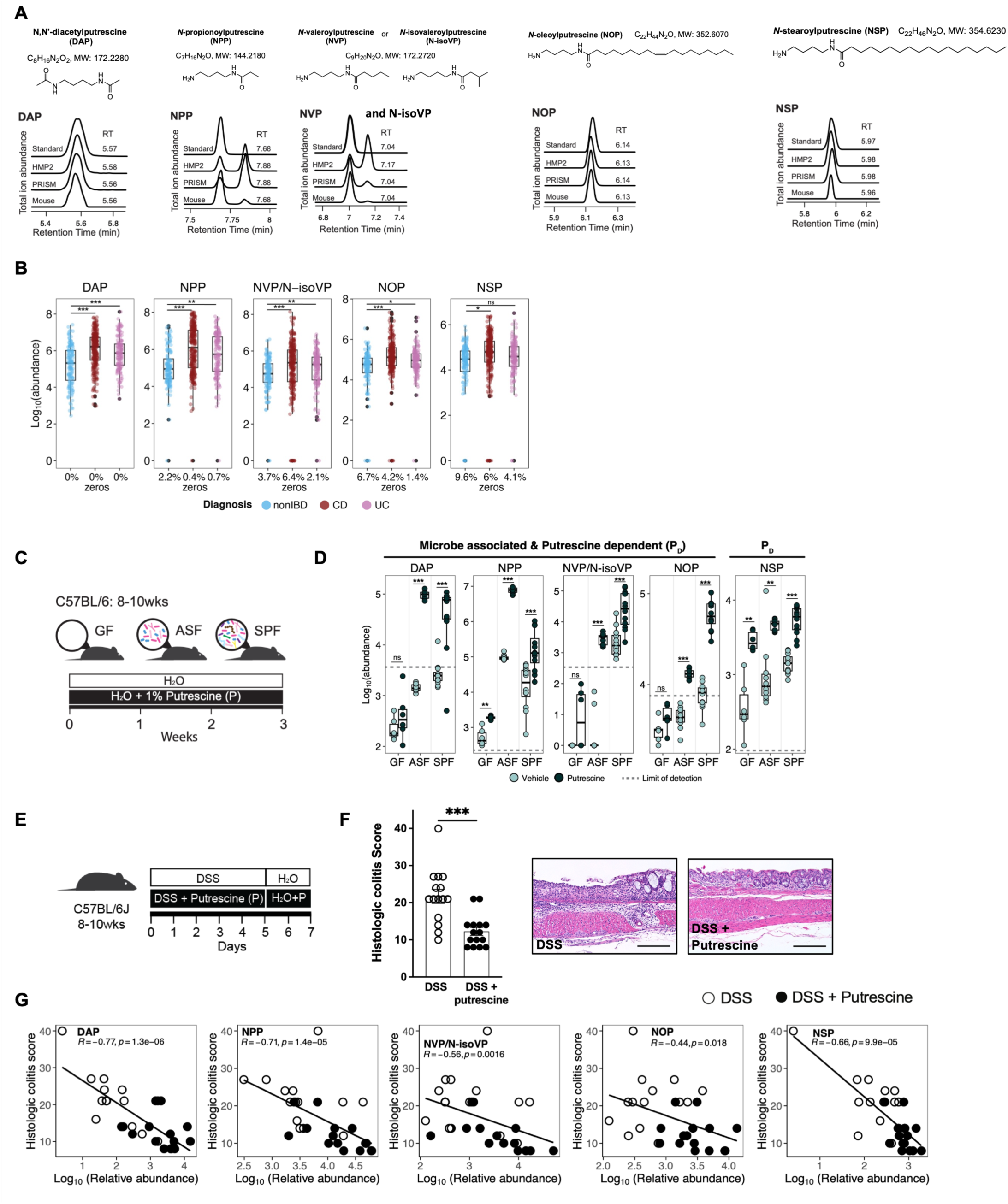
Discovery of microbe-associated putrescine derivatives with a strong IBD and intestinal inflammation association. (A) Acylated putrescines with chemical structures, formulas, molecular weights, and validation in HMP2 and PRISM (human IBD cohorts) and mouse metabolomics data with synthetic standards, total ion abundance, and retention time (RT). (B) Box plots of the relative abundances of the acylated putrescines in HMP2 fecal samples (non-IBD n=135, CD n=265, UC n=146). Percentage of zeros shown on x-axis. Statistical analysis performed using a linear mixed model. *q-value (FDR-adjusted p-value) < 0.2, **q < 0.05, ***q < 0.01. (C) Experimental schematic of gnotobiotic mice treated with or without 1% w/v putrescine in drinking water for stool metabolomics (GF control n=6, putrescine n=6; ASF control n=10, putrescine n=9; SPF control n=12, putrescine n=12). (D) Box plots of relative abundances of DAP, NPP, NVP/N-isoVP, NOP, and NSP from GF (control n=6; putrescine n=6), ASF (control n=10; putrescine n=9), and SPF (control n=12; putrescine n=12) C57BL/6 mice with and without putrescine supplementation. *p < 0.05, **p < 0.01, ***p < 0.001, n.s. not significant, Wilcoxon test. Gray dotted line indicates the limit of detection, determined from estimated fecal acylated putrescine concentrations using an external calibration curve with synthetic standards. (E) Experimental schematic of DSS intestinal injury and inflammation model. (F) Histologic colitis scores of putrescine-supplemented (n=15) and control (n=15) mice with 3% w/v DSS treatment. Data reflect 3 independent experiments. Circles represent individual mice; means ± SEM shown; ***p < 0.001, two-tailed Mann-Whitney U test. Representative photomicrographs of H&E-stained colon sections. Scale bars, 200 µm. (G) Spearman’s correlation coefficient and P values for relative abundances of DAP, NPP, NVP/N-isoVP, NOP, and NSP versus histologic colitis scores. Control mice (n=14, white circles) and putrescine-supplemented mice (n=15, black circles). Boxplot boxes (B, C) indicate 1st, 2nd, and 3rd quartiles. Boxplot whiskers (B) indicate inner fences with outliers plotted individually.

All five newly identified acylated putrescines, together with NMP, were highly prevalent in human fecal samples. All but one were significantly enriched in both CD and UC compared to non-IBD controls in the HMP2 cohort (non-IBD n=135, CD n=265, UC n=146; Figure 2B; Figure S5C). DAP, NPP, NOP, NSP, and NMP were further enriched in CD patients in the independent PRISM cohort compared to non-IBD controls (Figures S4A and S5C), supporting conserved enrichment of these metabolites across IBD patient populations. NMP did not cluster with the other acylated putrescines in the HMP2 abundance covariance analysis, suggesting that its production in the human gut may be governed by distinct factors and that it represents a functionally independent member of this metabolite class.

### The gut microbiota converts putrescine into acylated putrescines that accumulate during intestinal inflammation

Putrescine, the shared diamine backbone of these metabolites, is widely prevalent in human diets^22,23^ but present only in trace amounts in our gnotobiotic mouse chow. To determine if acylation of putrescine is microbiota-associated, we supplemented GF, ASF, and SPF mice with putrescine in the drinking water and profiled fecal acylated putrescine levels (Figure 2C). Putrescine supplementation elevated fecal putrescine and N-acetylputrescine levels without significantly altering microbiome composition (Figures S4B and S4D–E; Tables S7–S8). DAP, NPP, NVP/N-isoVP, NOP, and NBP were significantly elevated in putrescine-supplemented ASF and SPF mice compared to GF mice (Figure 2D; Figure S4C; Table S8), establishing their biosynthesis as microbiota-associated and putrescine-substrate-dependent. In contrast, NSP and NMP were elevated by putrescine supplementation comparably across all three microbial contexts, including GF mice, indicating that both host and microbial biosynthetic activity can contribute to their production (Figure 2D; Figure S5D). Together, these data demonstrate that DAP, NPP, NVP/N-isoVP, NOP, and NBP are microbe-associated and putrescine-dependent, whereas NSP and NMP are putrescine-dependent metabolites accessible through both host and microbial biosynthesis.

To confirm that these acylated putrescines are of microbial origin specifically in humans, we examined fecal metabolomes from the Food and Resulting Microbial Metabolites (FARMM) study,^21^ a longitudinal antibiotic intervention cohort in which gut microbiota depletion was induced by antibiotic and polyethylene glycol treatment in diet-controlled inpatient healthy subjects (Figure S3A). DAP, NOP, and NSP were detected in FARMM fecal samples, whereas NPP and NVP/N-isoVP were not detected, likely reflecting differences in the composition or activity of the resident microbiota in this cohort (Figure S3B). DAP and NOP were significantly depleted in omnivores following microbiota depletion, supporting their gut microbiota association in humans (Figure S3C).

To explore if acylated putrescines accumulate under intestinal inflammatory conditions and if this accumulation is microbiota-associated, we assessed their levels in GF and SPF mice with DSS-induced colitis supplemented with putrescine in the drinking water (Figure 2E; Figure S4F). Putrescine supplementation had no significant effect on histologic colitis scores in GF mice (Figure S4G), confirming that putrescine itself does not directly protect against intestinal injury in the absence of the microbiota. In contrast, putrescine supplementation significantly reduced colitis severity in SPF mice (Figure 2F), implicating microbiota-mediated putrescine metabolism in modulating intestinal inflammation. Consistent with this, fecal levels of NAP, NPP, NVP/N-isoVP, NOP, NSP, and NBP were inversely correlated with histologic colitis scores in SPF mice (Figure 2G; Figures S4H and S5E; Table S9), whereas putrescine itself and NMP showed no significant correlation. These data demonstrate that microbiota-derived acylated putrescines accumulate under pro-inflammatory intestinal conditions and are negatively associated with disease severity, identifying them as compelling candidates for functional investigation.

### NOP is a uniquely bioactive acylated putrescine produced by *Enterocloster* species

To identify which, if any, of the acylated putrescines and their polyamine precursors exert host bioactivity, we performed 3′ Digital Gene Expression (3′-DGE) profiling^24^ of mouse BMDCs and colonic organoids treated with putrescine, spermidine, spermine, and each of the acylated putrescines. Dendritic cells were selected given their central roles in initiating and regulating adaptive immune responses critical to IBD pathogenesis,^25,26^ while colonic organoids were included to capture potential epithelial responses.^27,28^ Among all compounds tested, NOP was uniquely and potently bioactive, inducing 488 differentially expressed genes in BMDCs and 318 in colonic organoids, in striking contrast to the minimal transcriptional effects observed with polyamines and all other acylated putrescines combined (n=56 differentially expressed genes) (Figure 3A–B; Figure S5F; Table S10). These data identified NOP as the primary immunomodulatory candidate within this metabolite class and motivated investigation of its microbial biosynthesis and immunoregulatory mechanisms.

**Figure 3.**
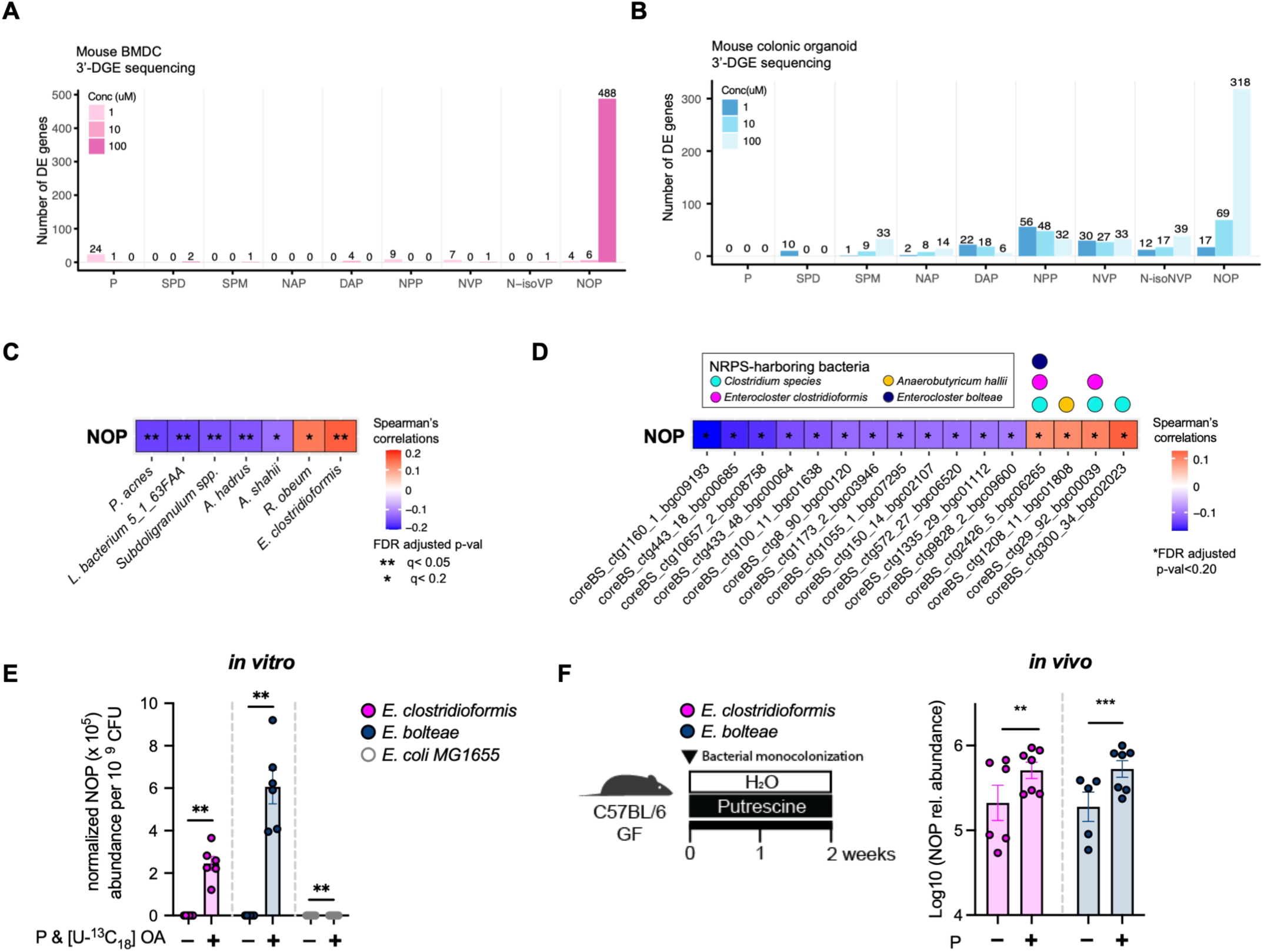
N-oleoylputrescine (NOP) bioactivity and source identification. (A) Number of differentially expressed genes (Wald’s test, FDR-adjusted P < 0.05) in mouse BMDCs co-cultured with polyamines (P: putrescine, SPD: spermidine, SPM: spermine) and acylated putrescines (NAP: N-acetylputrescine, DAP, NPP, NVP, N-isoVP, NOP) at 1, 10, 100 µM, profiled using 3′-DGE sequencing. (B) Number of differentially expressed genes in mouse colonic organoids treated identically. Mouse BMDC and colonic organoid transcriptional profiling assays (A, B) were performed in triplicate for each compound. (C–F) Identification of NOP-producing bacteria. (C) Correlation between microbial species abundances and NOP abundance in HMP2 (Spearman correlation with FDR-adjusted p-value; *q < 0.2, **q < 0.05). (D) Correlation between abundances of core NRPS biosynthetic gene clusters and NOP abundance in HMP2 (Spearman correlation; *q < 0.2). NRPS-harboring bacteria and their BGCs positively correlated with NOP abundance, with taxa indicated by colored circles. Correlation analyses (C, D) based on residual microbial and metabolite abundance after correcting for diagnosis, consent age, and antibiotic use. (E) Bar plot of relative NOP abundance in NRPS-harboring human bacterial isolates cultured with putrescine (P, 100 µM) and labeled oleic acid ([U-13C₁₈] OA, 100 µM), measured in cell pellets (3 biological replicates per isolate; 2 independent experiments). NOP abundance normalized to CFU; *E. coli* MG1655, which lacks NRPS clusters, used as negative control. Data shown as mean ± SEM. *p < 0.05, **p < 0.01, two-tailed Mann-Whitney U test. (F) Bar plot of relative fecal NOP abundance in GF C57BL/6 mice mono-colonized with NRPS-harboring bacteria with or without putrescine supplementation (2 independent experiments). Data shown as mean ± SEM. **p < 0.01, ***p < 0.001, linear model with Dunnett-adjusted p-value.

To identify which gut bacteria produce NOP, we analyzed 461 paired metagenomic and metabolomic samples from the HMP2 cohort by correlating microbial species abundances with NOP abundance (STAR Methods). *Enterocloster clostridioformis* showed the strongest positive association with NOP abundance among all species examined (Figure 3C). To investigate the biosynthetic basis of NOP production, we examined NRPS, enzymatic components of microbial BGCs known to mediate fatty acid amide conjugation. Using a curated, non-redundant core NRPS dataset,^10^ we identified four core NRPS genes that positively correlated with NOP abundance in HMP2, harbored by *Clostridium* species, *E. clostridioformis*, *Anaerobutyricum hallii*, and *Enterocloster bolteae* (Figure 3D), implicating NRPS-dependent biosynthetic machinery in NOP production by specific members of the human gut microbiota.

To directly test if NRPS-harboring bacteria produce NOP, we cultured two human intestinal isolates, *E. clostridioformis* 2_1_49FAA and *E. bolteae* 22-5-S 3 D6FAA, with isotope-labeled substrate ([U-13C₁₈] oleic acid) and putrescine, and quantified NOP production normalized to colony-forming units (CFU). Both isolates produced significantly more NOP under substrate-supplemented conditions compared to unsupplemented controls, and NOP production by both NRPS-harboring strains was significantly greater than that of *E. coli* MG1655, which lacks NRPS biosynthetic gene clusters, included as a negative control (Figure 3E). To confirm microbial NOP biosynthesis in vivo, GF C57BL/6 mice were mono-colonized with either *E. clostridioformis* or *E. bolteae* and provided with putrescine or vehicle control in the drinking water. Intestinal NOP levels were significantly elevated in mice receiving putrescine supplementation compared with vehicle-treated mono-colonized controls (Figure 3F). Notably, low-level NOP production was also detected in vehicle-treated mono-colonized mice, consistent with the ability of both *E. clostridioformis* and *E. bolteae* to catabolize dietary spermine and spermidine to putrescine, providing endogenous substrate even in the absence of supplementation. Together, these in vitro and in vivo data confirm that specific NRPS-harboring gut bacteria are sufficient for NOP biosynthesis, with production dependent on precursor availability.

### NOP suppresses inflammatory pathways central to IBD in mouse and human myeloid cells

Given NOP’s potent transcriptional activity in BMDCs (Figure 3A; Figures S6A–B; Table S11), we characterized its anti-inflammatory transcriptional program using full-length RNA sequencing, leveraging the central role of dendritic and myeloid cells in IBD and available human IBD immune cell transcriptomic datasets for comparison (for NOP-treated colonic organoid full-length RNA-seq, see Figure S10A). Hallmark pathway enrichment analysis revealed significant and dose-dependent downregulation of five core IBD-related immune signaling:^29–31^ TNFα signaling via NF-κB, interferon-γ signaling, interferon-α signaling, inflammatory response, and IL-6/JAK/STAT3 signaling (Figures S6C–G). These same five pathways are significantly upregulated in inflamed CD and UC intestinal biopsies from the HMP2 cohort compared to non-inflamed non-IBD tissues (Figure 4A; Tables S12–S13), demonstrating that NOP suppresses precisely the inflammatory programs pathologically elevated in human IBD.

**Figure 4.**
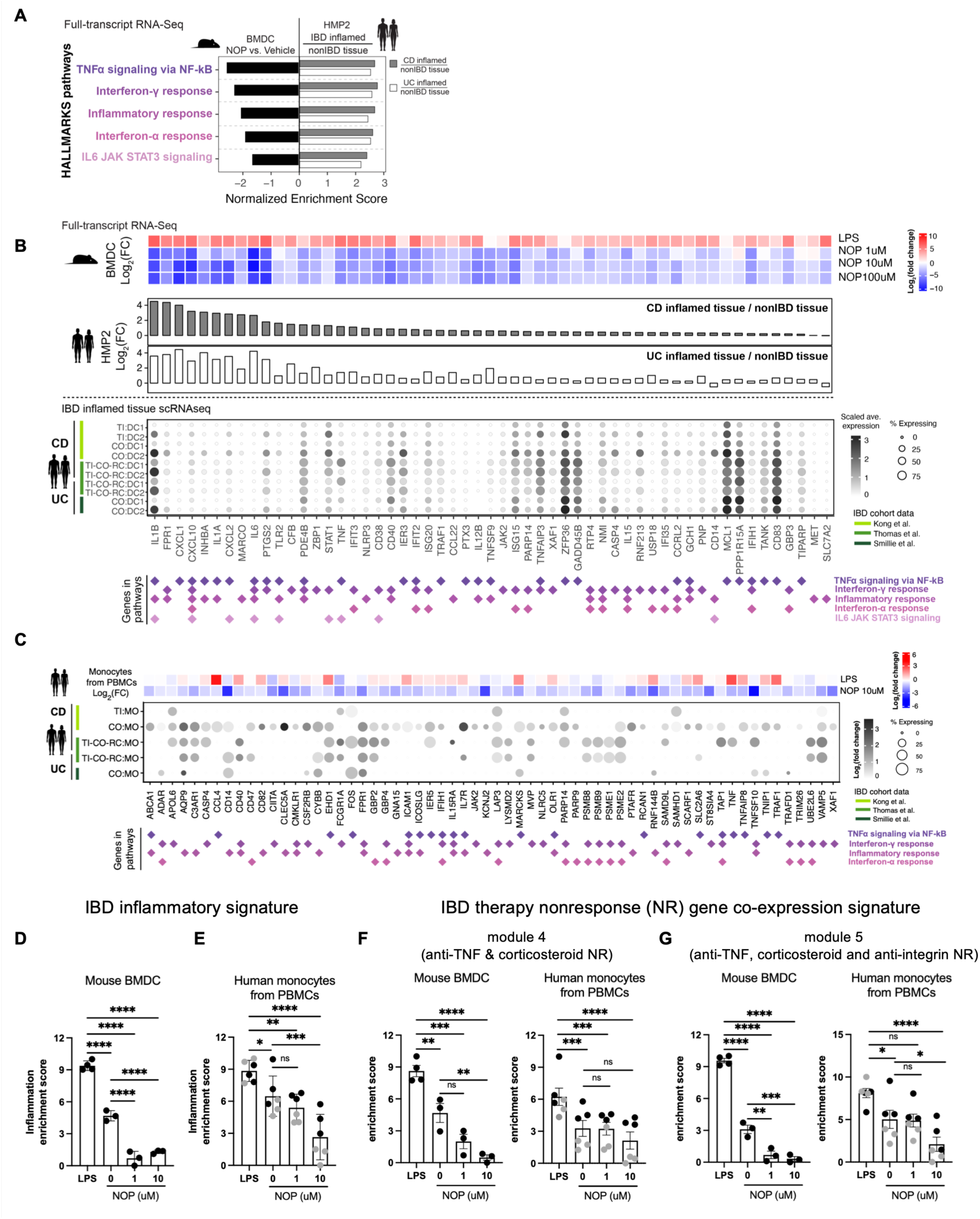
NOP suppresses IBD-associated inflammatory pathways in mouse and human myeloid cells. (A) Hallmark pathway enrichment analysis of HMP2 intestinal biopsy RNA-seq data comparing IBD-inflamed tissue (CD inflamed, UC inflamed) to non-IBD non-inflamed tissue. (B) Heatmap of genes downregulated (Wald’s test, FDR-adjusted P < 0.05) in BMDCs by NOP (1, 10, 100 µM) and upregulated in human inflamed CD or UC tissues (Wald’s test, FDR-adjusted P < 0.05). Top heatmap: log₂ fold change of NOP-treated BMDCs vs. vehicle, with LPS as positive control. Middle bar plot: log₂ fold change in HMP2 CD (gray) and UC (white) inflamed vs. non-IBD uninflamed tissues. Bottom dot plot: average expression (dot color) and fraction of expressing cells (dot size) in human intestinal DC populations (DC1: CLEC9A, XCR1; DC2: CLEC10A, FCER1A) across three public scRNA-seq IBD datasets.^32–34^ Biopsy locations: terminal ileum (TI), colon (CO), rectum (RC). (C) Human monocyte transcriptional profiling. Top heatmap: genes downregulated (Wald’s test, FDR-adjusted P < 0.05) by 10 µM NOP in human PBMC-derived monocytes and upregulated (likelihood ratio test, FDR-adjusted P < 0.2) in IBD inflammatory monocytes in at least one of three scRNA-seq datasets. LPS treatment as positive control. Bottom dot plot: fraction of expressing cells (dot size) and log₂ fold change (dot color) in monocytes from IBD inflamed vs. non-IBD non-inflamed tissues across three scRNA-seq datasets. Genes associated with specific inflammatory pathways indicated by diamond symbols (B, C). Mouse BMDC and human monocyte transcriptional profiling assays (A–C) performed in triplicate. (D–E) Inflammation enrichment scores in NOP- and control-treated mouse BMDC (D) and human monocyte (E) transcriptomes, calculated using inflammatory gene signatures derived from Thomas et al. (Thomas et al., 2024). (F–G) Enrichment scores for IBD therapy non-response (NR) gene co-expression signatures from Friedrich et al.^35^. (F) Module 4 enrichment scores (anti-TNF and corticosteroid non-responders) in BMDC and PBMC-derived monocyte transcriptomes. (G) Module 5 enrichment scores (anti-TNF, corticosteroid, and anti-integrin non-responders). Data are presented as mean ± SEM. *p < 0.05, **p < 0.01, ***p < 0.001, ****p < 0.0001, n.s. not significant. One-way ANOVA with Dunnett-adjusted p-value for (D) and BMDC datasets in (F, G); pairwise comparisons of estimated means with Tukey-adjusted p-values for (E) and monocyte datasets in (F, G) (STAR Methods). Data from two independent PBMC donors indicated with different colors (E, F, G).

To directly evaluate the human relevance of NOP-responsive pathways, we compared genes downregulated by NOP in BMDCs with genes upregulated in human IBD including those identified in the HMP2 biopsy transcriptome and intestinal dendritic cell populations in three independent scRNA-seq.^32–34^ NOP significantly downregulated genes upregulated in HMP2 IBD-inflamed tissues, and were both prevalently and highly expressed in human intestinal DC populations across all three scRNA-seq datasets (Figure 4B). To determine if this anti-inflammatory transcriptional program is conserved in primary human myeloid cells, we profiled the transcriptional response to NOP in human PBMC-derived monocytes. NOP significantly downregulated genes upregulated in inflammatory monocyte populations from inflamed IBD intestinal tissue across three independent IBD scRNA-seq datasets (Figure 4C). Genes with established roles in IBD pathogenesis enriched in inflamed IBD monocytes,^32^ including TNF, FPR1, and JAK2, were significantly suppressed by NOP in human monocytes (Figure S6J), confirming that NOP’s anti-inflammatory transcriptional effects are conserved across mouse and human myeloid cells.

To assess the translational relevance of NOP’s anti-inflammatory effects in the context of active human IBD, we evaluated its ability to suppress published gene signatures of histologic inflammation and IBD therapy non-response. Using an inflammation gene signature derived from differentially expressed genes between histologically inflamed IBD resections and non-inflamed/non-IBD gut tissue,^34^ NOP treatment significantly reduced inflammation enrichment scores in both mouse BMDCs and human monocytes (Figure 4D–E). The magnitude of this reduction was similar to the difference reported between IBD patients in remission versus non-remission in the Thomas et al.^34^ cohort, suggesting NOP suppresses a transcriptional program directly relevant to active human IBD disease activity. To further address potential patient heterogeneity in NOP responsiveness, we evaluated enrichment of gene co-expression modules associated with IBD therapy non-response:^35^ module 4, predictive of non-response to anti-TNF therapy and corticosteroids, and module 5, predictive of non-response to anti-TNF, corticosteroids, and anti-integrin therapies, spanning the major therapeutic classes used in IBD management. NOP treatment significantly reduced enrichment scores for both modules in mouse BMDCs (Figure 4F–G). In human monocytes, NOP significantly reduced module 5 enrichment scores and showed a trend toward reduced module 4 enrichment scores compared to untreated controls (Figure 4F–G). Enrichment scores for both modules were significantly lower in NOP-treated cells compared to LPS-treated cells. We observed donor-dependent differences in baseline enrichment scores and in the magnitude of score changes between module 4 and module 5, providing initial evidence for patient heterogeneity in NOP responsiveness and motivating future evaluation in patient-derived immune cells. Collectively, these data establish that NOP suppresses gene expression programs associated with histologic inflammation, active IBD, and treatment resistance across the major current IBD therapeutic classes.

### NOP suppresses colitis in multiple murine models through innate and adaptive immune mechanisms

To evaluate NOP’s anti-inflammatory activity in vivo, we selected a treatment dose of 24 mg/kg per day based on tolerability and minimal effect on body weight in dose-range testing (Figure S8A). Direct NOP treatment (24mg/kg/day) in mice resulted in fecal NOP concentrations that were 2-20-fold higher than those observed in HMP2 and PRISM cohorts (Figures S8B-C). Given that TNFα signaling via NF-κB was the most downregulated pathway in NOP-treated BMDCs, and that dendritic cells are central drivers of innate immune activation in IBD, we first tested NOP in the anti-CD40 colitis model, an NF-κB activation and dendritic cell-driven model of intestinal inflammation^36,37^ (Figure 5A). NOP treatment significantly reduced histologic colitis scores compared to vehicle control (Figure 5B). Consistent with transcriptional suppression of TNF, IL-1, and IL-6 pathway genes in NOP-treated BMDCs, we observed significant reductions in the proinflammatory cytokines IL-1α, IL-1β, and IL-6, and chemokine CCL2 in proximal colonic tissue homogenates from NOP-treated mice (Figure 5C), confirming that NOP exerts anti-inflammatory effects mediated through innate immune cells in vivo.

**Figure 5.**
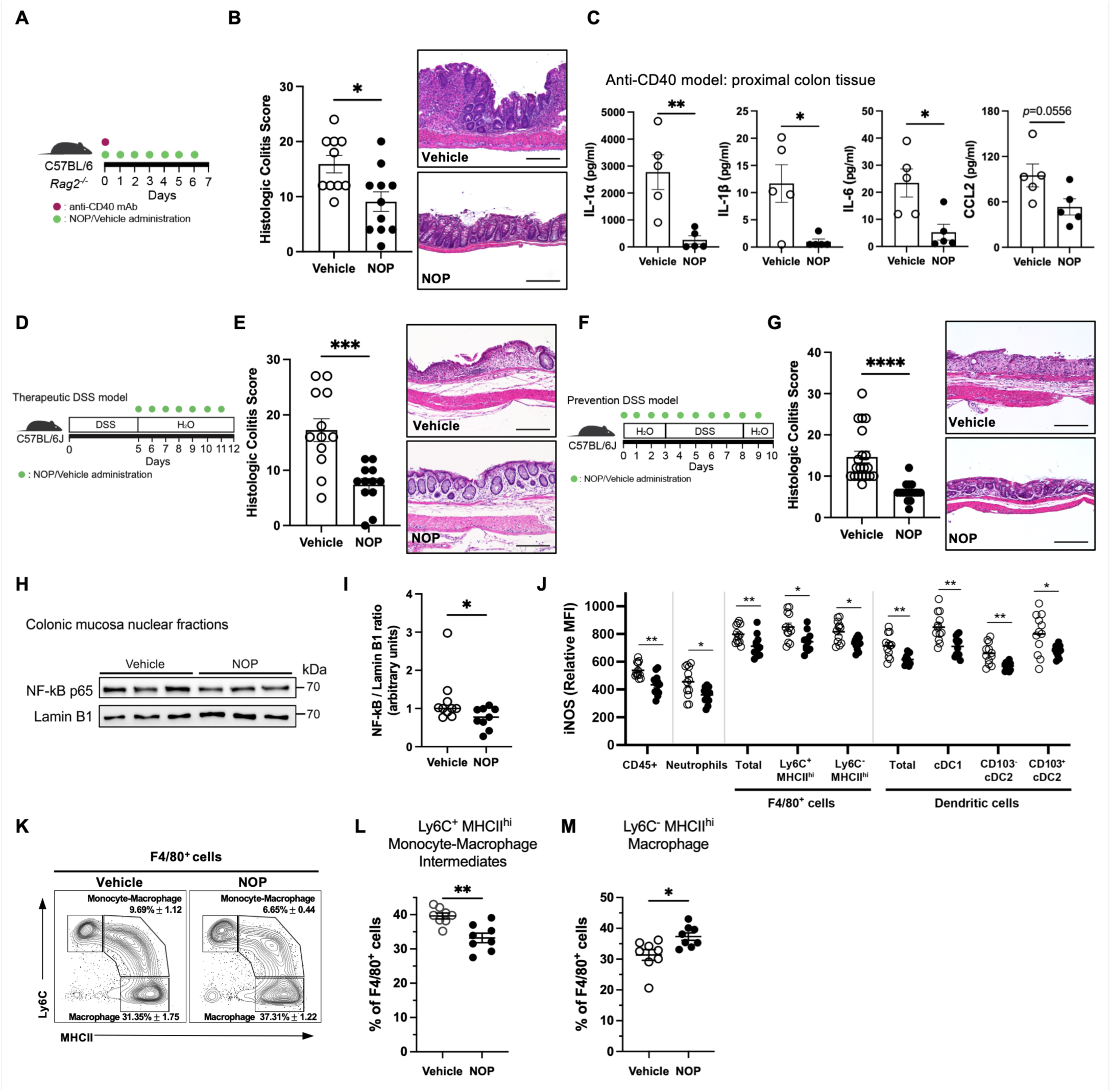
NOP treatment suppresses inflammatory responses in IBD mouse models. (A) Schematic of anti-CD40 colitis model. (B) Histologic colitis scores of vehicle-treated (n=11) or NOP-treated (n=10) mice with representative photomicrographs. Scale bars, 200 µm; 2 independent experiments. (C) IL-1α, IL-1β, IL-6, CCL2 cytokine levels in proximal colon tissue homogenates (vehicle n=5; NOP n=5). (D) Schematic of 3% w/v DSS-induced colitis therapeutic model. (E) Histologic colitis scores of vehicle-treated (n=12) or NOP-treated (n=12) mice with representative photomicrographs. Scale bars, 200 µm; 4 independent experiments. (F–M) 3% w/v DSS-induced colitis prevention model. (F) Experimental schematic. (G) Histologic colitis scores of vehicle-treated (n=20) and NOP-treated (n=16) mice with representative photomicrographs. Scale bars, 200 µm; 4 independent experiments. (H) Representative NF-κB p65 immunoblot of nuclear extracts from colonic mucosa and (I) relative quantification of p65 levels, normalized to Lamin B1 (vehicle n=10; NOP n=9); 3 independent experiments. (J) Intracellular iNOS expression in colonic lamina propria CD45⁺ cells, neutrophils, F4/80⁺ cells, and dendritic cells (vehicle n=12; NOP n=12). (K) Representative FC plots and (L) FC analysis of Ly6C⁺ MHCII^hi^ F4/80⁺ monocyte-macrophage intermediates, (M) Ly6C⁻ MHCII^hi^ F4/80⁺ macrophages. Flow cytometry (FC) analysis (vehicle n=8; NOP n=8); 2 independent experiments. Circles represent individual mice; mean ± SEM shown. *p < 0.05, **p < 0.01, ***p < 0.001, ****p < 0.0001. Two-tailed Mann-Whitney U test (B, C, E, G, I, J, L, M).

To determine if NOP’s anti-inflammatory activity generalizes across models with distinct mechanisms of inflammation induction and different genetic backgrounds, we evaluated NOP in two additional colitis models. NOP significantly attenuated histologic colitis scores in TNBS-treated BALB/c mice (Figures S8D–E), a hapten-driven model of Th1-mediated colitis, and in DSS-treated C57BL/6J mice following established colitis, a therapeutic rather than preventive treatment paradigm (Figure 5D–E). Collectively, the efficacy of NOP across mechanistically distinct colitis models, anti-CD40 (NF-κB/DC-driven), TNBS (hapten/Th1-driven), and DSS (epithelial injury-driven), in two different mouse strains demonstrates that NOP exerts broad, robust anti-inflammatory effects that promote resolution of intestinal inflammation.

Given that a central goal of IBD therapy is disease interception and the prevention of inflammatory flares, we next evaluated NOP in a colitis prevention paradigm, administering NOP prior to and during DSS exposure (Figure 5F). NOP significantly reduced histologic colitis scores in this prevention model (Figure 5G). To probe the mechanism of protection in vivo, we examined NF-κB activation by western blot of nuclear extracts from the colonic mucosa. NOP-treated mice showed significantly reduced nuclear NF-κB p65 compared to vehicle-treated controls (Figure 5H–I), confirming in vivo suppression of a key mediator of intestinal inflammation. Consistent with the significant downregulation of NOS2 in NOP-treated BMDC (Figures S6H-I), NOP treatment significantly suppressed the NF-κB target enzyme iNOS^36,37^ in total colonic lamina propria immune cells, neutrophils, F4/80⁺ macrophages, and dendritic cells (Figure S7, Figure 5J), directly linking the transcriptional anti-inflammatory program observed in vitro to functional suppression of nitric oxide production in vivo.

Profiling of colonic lamina propria immune cell populations by flow cytometry in the DSS prevention model revealed that NOP modestly decreased pro-inflammatory Ly6C⁺MHCII^hi^F4/80⁺ monocyte-macrophage intermediate populations (Figure 5K–L) and slightly increased Ly6C⁻MHCII^hi^ tissue-resident macrophages (Figures 5K and 5M), a shift consistent with reduced inflammatory monocyte recruitment and enhanced tissue-reparative macrophage polarization. No significant changes were observed in neutrophil or dendritic cell populations (Figures S8F–I). Importantly, none of these myeloid population changes were observed under homeostatic conditions in the absence of intestinal inflammation (Figures S8J–Q), establishing that NOP’s immunomodulatory effects on myeloid cells are inflammation-context-dependent. Given that these myeloid population changes were marginal and likely insufficient to fully account for the observed histologic protection, these findings suggested that additional immune cell compartments, particularly lymphocytes, may contribute meaningfully to NOP’s anti-inflammatory activity.

### NOP suppresses type 1 T cell responses through a T cell-intrinsic mechanism

Given that lymphocytes are central mediators of intestinal inflammation and well-established drivers of disease in the DSS model,^38–40^ we examined lymphocyte populations in the DSS prevention model. In the CD4⁺ T cell compartment, NOP significantly reduced the frequency of T-bet⁺ Th1 cells (Figure 6A–B) and IFN-γ⁺T-bet⁺ Th1 cells (Figure 6C), consistent with suppression of type 1 helper T cell responses. While decreases in intracellular IFN-γ expression within T-bet⁺CD4⁺ T cells were modest and did not reach statistical significance (Figure 6D), the directionality was consistent. In the CD8⁺ T cell compartment, NOP significantly reduced frequencies of activated CD44^hi^ CD8⁺ T cells (Figures 6E–F), T-bet⁺ CD8⁺ T cells (Figure 6I), IFN-γ⁺ CD44^hi^ CD8⁺ T cells (Figure 6G), IFN-γ⁺T-bet⁺ CD44^hi^ CD8⁺ T cells (Figure 6J), and significantly reduced intracellular IFN-γ expression in both CD44^hi^ and T-bet⁺ CD8⁺ T cell populations (Figures 6H and 6K). NOP also reduced T-bet⁺ B cell frequencies (Figure S9A), suggesting broader suppression of T-bet-driven type 1 immune responses beyond T cells. Importantly, none of these effects were observed under homeostatic conditions in the absence of intestinal inflammation (Figures S9B–E), demonstrating that NOP’s suppression of type 1 lymphocyte responses is inflammation-context-dependent, mirroring the pattern observed for myeloid cells.

**Figure 6.**
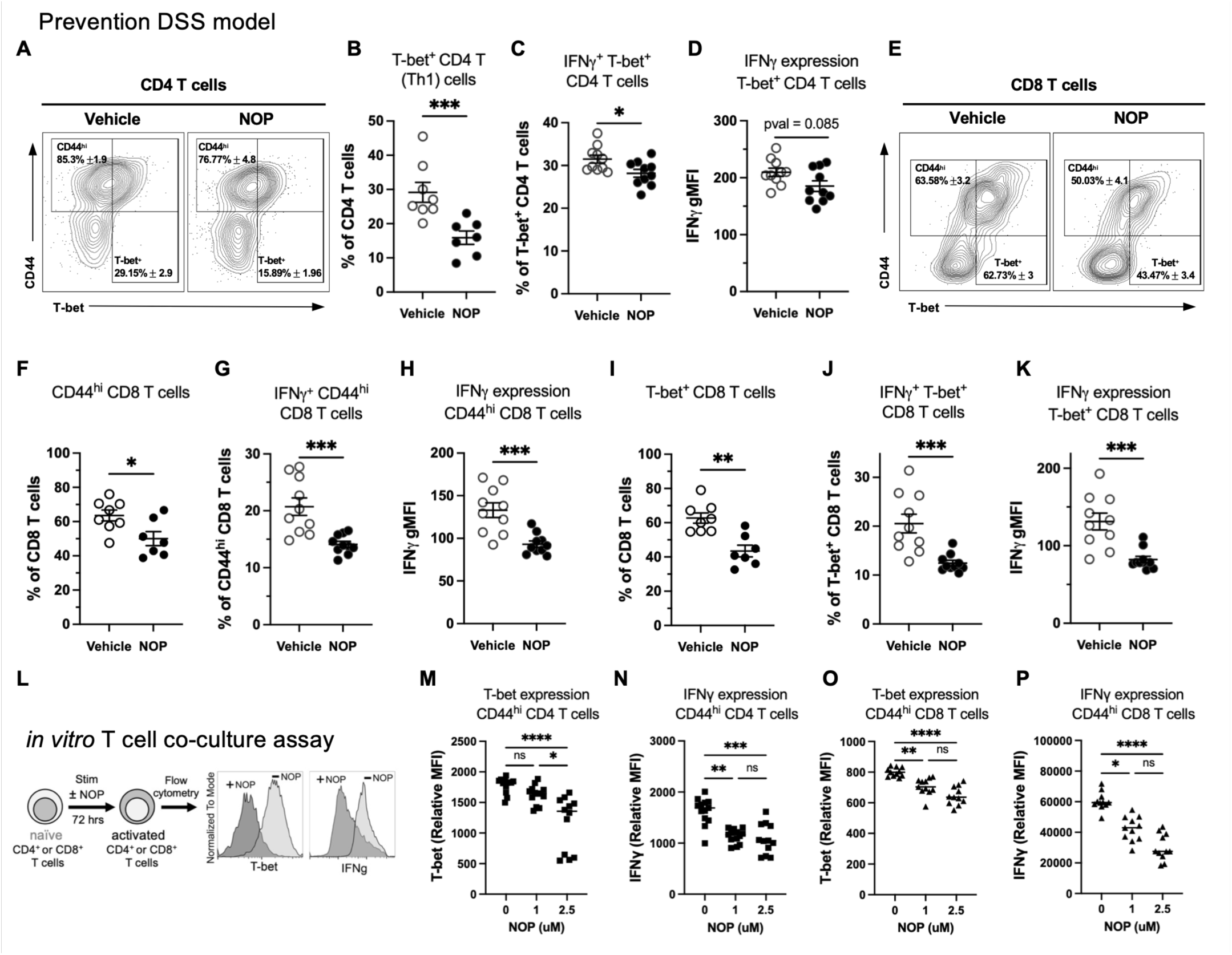
NOP decreases T-bet⁺ IFN-γ-producing CD4⁺ and CD8⁺ T cells in vivo and reduces T-bet and IFN-γ in vitro. (A) Representative FC plots and (B) FC analysis of T-bet⁺ CD4⁺ T (Th1) cells. (C) FC analysis of IFN-γ⁺T-bet⁺ CD4⁺ T cells and (D) intracellular IFN-γ expression in T-bet⁺ CD4⁺ T cells. (E) Representative FC plots and (F) FC analysis of CD44^hi^ CD8⁺ T cells. (G) FC analysis of IFN-γ⁺ CD44^hi^ CD8⁺ T cells and (H) intracellular IFN-γ expression in CD44^hi^ CD8⁺ T cells. (I) FC analysis of T-bet⁺ CD8⁺ T cells. (J) FC analysis of IFN-γ⁺T-bet⁺ CD44^hi^ CD8⁺ T cells and (K) IFN-γ expression in T-bet⁺ CD8⁺ T cells. T cell profiling FC analysis: vehicle n=8, NOP n=8; 2 independent experiments (A–B, E–F, I). IFN-γ profiling FC analysis: vehicle n=10, NOP n=10; 2 independent experiments (C–D, G–H, J–K). Circles represent individual mice; mean ± SEM shown. *p < 0.05, **p < 0.01, ***p < 0.001. Two-tailed Mann-Whitney U test (B, D– K). (L–P) In vitro T cell co-culture assay; 3 independent experiments. (L) Experimental scheme. Naïve CD4⁺ or CD8⁺ T cells isolated from mouse spleens, activated with anti-CD3 and B7-1, and cultured with IL-2 and IL-12 in the presence or absence of NOP on day 0. T cells collected on day 3 and gated for flow cytometry. (M) FC analysis of T-bet and (N) IFN-γ expression in CD44^hi^ CD4⁺ T cells. (O) FC analysis of T-bet and (P) IFN-γ expression in CD44hi CD8⁺ T cells. Mean ± SEM shown. *p < 0.05, **p < 0.01, ***p < 0.001, ****p < 0.0001. Kruskal–Wallis test with Dunn’s post hoc multiple comparison analysis (M–P).

The in vivo data raised the question of whether NOP’s effects on T cells are mediated indirectly through NOP’s documented suppression of the myeloid cell compartment, or if NOP acts directly on T cells to suppress their effector function. To distinguish between these possibilities, we performed in vitro activation and differentiation assays of naïve CD4⁺ and CD8⁺ T cells cultured in the presence or absence of NOP, without myeloid cells present (Figure 6L; Figures S9F–G). Naïve T cells were activated with anti-CD3 and B7-1 and cultured with IL-2 and IL-12 to promote type 1 T cell differentiation. NOP treatment significantly suppressed T-bet and IFN-γ expression in both CD44hi CD4⁺ T cells (Figures 6M–N) and CD44^hi^ CD8⁺ T cells (Figures 6O–P) compared to vehicle-treated controls, demonstrating that NOP can directly suppress type 1 T cell effector programming through a T cell-intrinsic mechanism.

Collectively, the in vivo and in vitro data from Figures 5 and 6 demonstrate that NOP modulates intestinal inflammation through convergent effects on both myeloid and lymphocyte compartments, suppressing innate immune activation, NF-κB signaling, iNOS-mediated nitric oxide production, and type 1 T cell responses via mechanisms that are selectively engaged under inflammatory conditions.

## DISCUSSION

This study identifies N-acyl putrescines as a class of microbiome-associated metabolites enriched in the IBD gut and demonstrates that one member of this class, N-oleoylputrescine (NOP), exerts potent and broad anti-inflammatory activity across multiple experimental systems, including mouse and human myeloid cells, colonic organoids, and four murine colitis models. NOP suppresses core IBD-associated inflammatory pathways, reduces gene signatures of histologic inflammation and therapy non-response, and suppresses type 1 T cell responses through a cell-intrinsic mechanism. Collectively, these findings establish NOP as a biologically active microbiome-derived metabolite with properties relevant to IBD pathophysiology and therapy.

The enrichment of NOP in IBD despite its anti-inflammatory properties may appear paradoxical. We propose that this enrichment reflects a holobiont defense response, a compensatory metabolic program in which the gut microbiota upregulates biosynthesis of immunomodulatory metabolites in response to intestinal inflammatory signals, generating molecules that may help limit injury and contribute to homeostatic restoration. Several lines of evidence support this model. First, NOP accumulation is inflammation context-dependent in both mice and humans: acylated putrescine levels are elevated in DSS-treated SPF mice, inversely correlated with colitis severity, and enriched in IBD patients across two independent cohorts. Second, a structurally related host-derived anti-inflammatory fatty acid amide, palmitoylethanolamide (PEA), exhibits a strikingly parallel pattern, as it is enriched in both CD and UC patients. PEA is now being investigated in active, mild to moderate UC in a clinical trial (NCT07609810), suggesting that enrichment of anti-inflammatory metabolites in IBD may be a recurring feature of host-microbiome interactions under inflammatory stress rather than an isolated observation. We propose that the endogenous levels of NOP in IBD, estimated at 0.6–12 nM in HMP2 and PRISM fecal samples, fall below the threshold required to fully resolve established inflammation, analogous to PEA, but that this does not preclude therapeutic utility through exogenous supplementation to achieve higher concentrations.

The identification of *Enterocloster* species as NOP producers via NRPS biosynthetic gene clusters reveals a previously unappreciated metabolic axis connecting gut bacterial biosynthetic capacity to host immune cell regulation. The NRPS-mediated conjugation of oleic acid with putrescine to form NOP places this metabolite within a growing family of gut microbiota-derived fatty acid amide conjugates, which includes N-acyl amino acid GPCR ligands produced by gut-inhabiting Clostridia via analogous enzymatic chemistry.^19,20^ The dependence of NOP production on NRPS machinery and available putrescine substrate implies that host dietary putrescine intake and microbial community composition jointly regulate NOP biosynthesis, positioning NOP at the intersection of diet, microbiome, and immunity. The detection of low-level NOP in vehicle-treated germ-free mice mono-colonized with NOP-producing strains, attributable to endogenous catabolism of dietary spermine and spermidine, suggests that these bacteria provide their own substrate under limiting conditions, potentially buffering NOP production against fluctuations in dietary putrescine availability.

The divergent immunomodulatory properties of NMP and NOP, two structurally related acylated putrescines sharing similar IBD-associated abundance trajectories yet exerting markedly different effects on immune cell function, underscore a critical principle: chemical specificity in host-microbiome metabolic signaling is extraordinarily precise, and immunological function cannot be predicted from structural class membership alone. The host’s capacity to discriminate between a C14 saturated acyl chain (NMP) and a C18 monounsaturated acyl chain (NOP) with opposing immunological consequences highlights the importance of rigorous structural characterization of individual gut metabolites and suggests that other acylated putrescines may have distinct, and potentially opposing, immunological roles that warrant individual investigation. The pro-barrier-disrupting activity attributed to NMP^10^ versus the anti-inflammatory activity of NOP represents one such functional divergence within a single metabolite class.

The translational potential of NOP is supported by multiple lines of evidence presented here. NOP reduced inflammation enrichment scores in human monocytes at a magnitude comparable to the difference between IBD patients in remission versus non-remission,^34^ providing a clinically calibrated benchmark for its transcriptional potency. NOP also reduced enrichment of gene signatures predictive of non-response to anti-TNF therapy, corticosteroids, and anti-integrin therapies, the three major current IBD therapeutic classes, raising the possibility that NOP’s mechanism of action may be complementary to, or synergistic with, existing treatments and potentially efficacious in patients who fail standard therapy. The donor-dependent differences in NOP responsiveness observed in human monocytes motivate a systematic evaluation of NOP activity across a larger panel of patient-derived immune cells, which could identify biomarkers of NOP responsiveness and inform patient stratification in future clinical development.

Several important questions remain to be addressed. While NOP treatment achieved fecal concentrations approximately 2–20-fold higher than endogenous IBD levels, the therapeutic threshold for intestinal NOP concentration in humans is not established and cannot be extrapolated directly from mouse models. The molecular receptor or receptor-independent mechanism through which NOP suppresses NF-κB activation and type 1 T cell responses remains unknown; its structural similarity to oleoylethanolamide (OEA) and other N-acyl amides that activate PPAR-α and GPCRs including GPR119^20^ provides candidate mechanisms that warrant direct investigation. If NOP accumulates in advance of, or reactively to, inflammatory flares cannot be fully resolved from the cross-sectional and longitudinally sparse sampling in the HMP2 and PRISM cohorts; prospective studies with dense temporal sampling around disease flares will be required to establish these dynamics and to determine whether NOP biosynthesis is itself regulated by inflammatory signals in the gut microenvironment. Finally, NOP’s efficacy and safety in patients with IBD will require dedicated clinical evaluation.

This work demonstrates that the IBD gut microbiota produces a class of immunomodulatory acylated putrescines, and that at least one member, NOP, has the capacity to broadly suppress innate and adaptive immune responses central to IBD across multiple experimental systems and disease models. These findings expand the catalog of microbiome-derived metabolites with therapeutic potential in IBD, provide a mechanistic framework for understanding how microbial NRPS biosynthetic activity contributes to intestinal immune regulation, and offer a conceptual lens, the holobiont defense hypothesis, for interpreting the enrichment of anti-inflammatory metabolites in chronic inflammatory disease states more broadly. As the chemical space of gut microbial metabolites continues to be explored with increasing structural resolution, we anticipate that similar compensatory metabolic strategies will be revealed in other inflammatory contexts, reflecting an evolutionarily conserved principle of host-microbiome co-regulation.

## RESOURCE AVAILABILITY

### Lead contact

- Requests for further information and resources should be directed to and will be fulfilled by the lead contact, W. S. Garrett.

### Materials availability

- This study did not generate new unique reagents.

### Data and code availability

- The 16S rRNA gene amplicon and RNA-seq datasets are available from SRA BioProject PRJNA1259007.
- The raw untargeted metabolomics data have been submitted to Metabolic Workbench and will be made publicly available upon manuscript acceptance. Tables of processed metabolites are available as Supplemental Table S15 and 16.
- Raw HMP2 metabolomics data are available at the Metabolic Workbench, Project ID PR000639, and the table of processed metabolites are available at the IBDMDB website (https://ibdmdb.org). Raw data of PRISM metabolomics are available at the Metabolomic Workbench, Project ID PR000677, and the table of processed metabolites are available from the associated publication’s supporting information.

## ACKNOWLEDGMENTS

We thank the members of the Garrett and Huttenhower laboratories (Harvard T.H. Chan School of Public Health) for discussion. This work was supported by NIH NIDDK grants R24DK110499 (W.S.G., C.H., and R.J.X.) and the Kenneth Rainin Foundation (W.S.G.). We are especially grateful to the participants in the HMP2 and PRISM cohorts who made this study possible. The computations in this paper were run in part on the FASRC Cannon cluster supported by the FAS Division of Science Research Computing Group at Harvard University. *E. clostridioformis* 2_1_49FAA was generously provided by the laboratory of Dennis Kasper and *E. bolteae* 22-5-S 3 D6FAA was generously provided by the laboratory of Emma Allen-Vercoe.

## AUTHOR CONTRIBUTIONS

S.B., C.H., and W.S.G. designed the research (Conceptualization). S.B. carried out computational analysis and experiments (Investigation, Formal Analysis). J.A.-P. and C.B.C. performed mouse stool untargeted metabolomic data acquisitions and acylated putrescines prediction and validation in mouse fecal, HMP2, and PRISM metabolomics (Investigation, Methodology). Sunghee Bang and J.C. performed the synthesis of acylated putrescines (Investigation, Resources). S.L.C. performed flow cytometry experiments and analysis (Investigation, Formal Analysis). M.C.S. conducted in vitro T cell activation assays (Investigation). M.M. performed microscopy to obtain images (Investigation). D.B.G. and G.P. performed transcriptional profiling (Investigation, Methodology). J.N.G. performed histologic assessments of colitis (Investigation, Formal Analysis). N.A., D.F.-P., E.A.F., E.C., Y.G.C., Y.Z., A.B., R.M.P., H.V., G.E.T., S.P.C., X.C.M., and R.J.X. participated in the investigation. S.B. and W.S.G. wrote the manuscript with feedback from all authors (Writing – Original Draft; Writing – Review & Editing). C.H., E.A.F., and W.S.G. supervised the research (Supervision). All authors approved the final manuscript.

## STAR★METHODS

### KEY RESOURCES TABLE

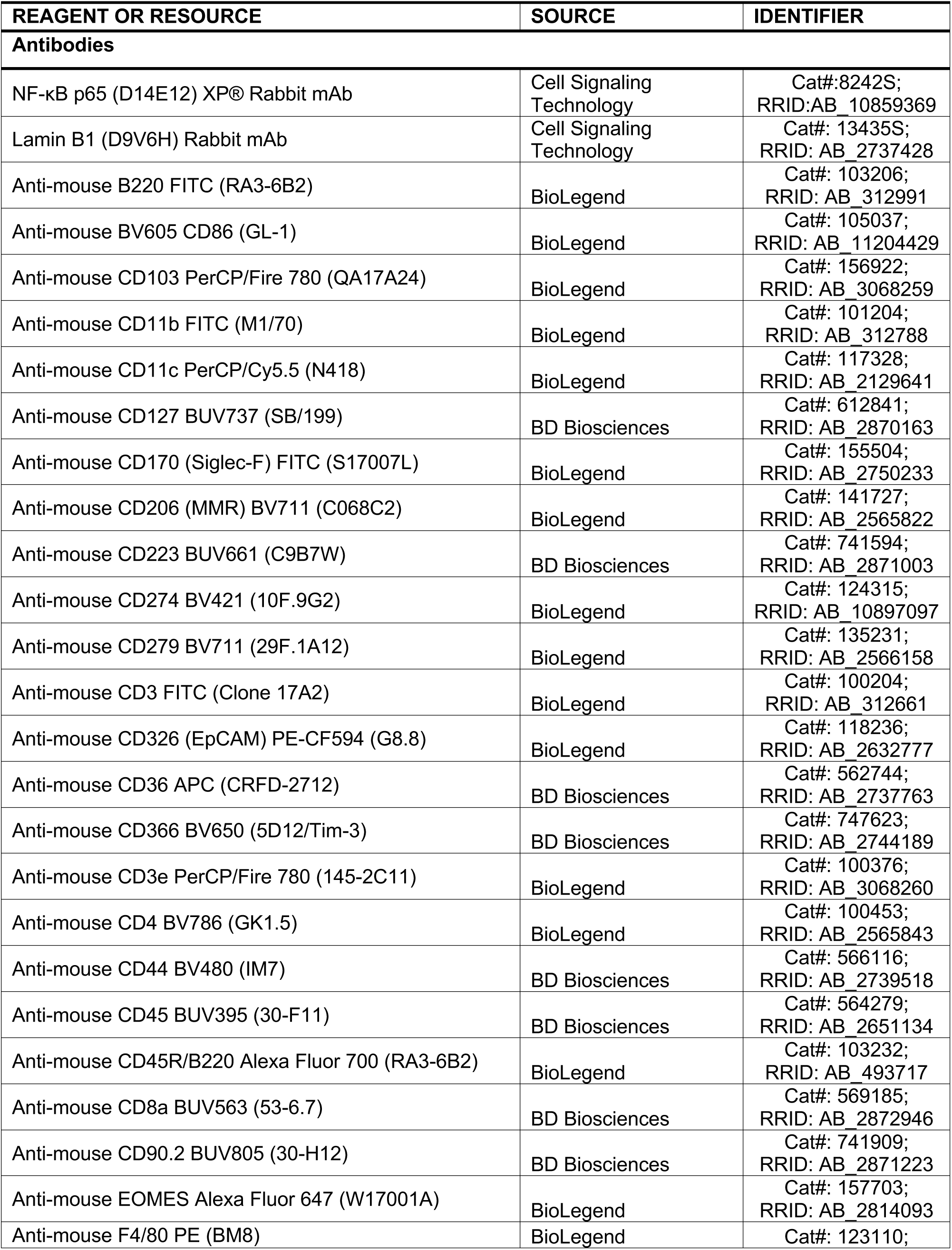
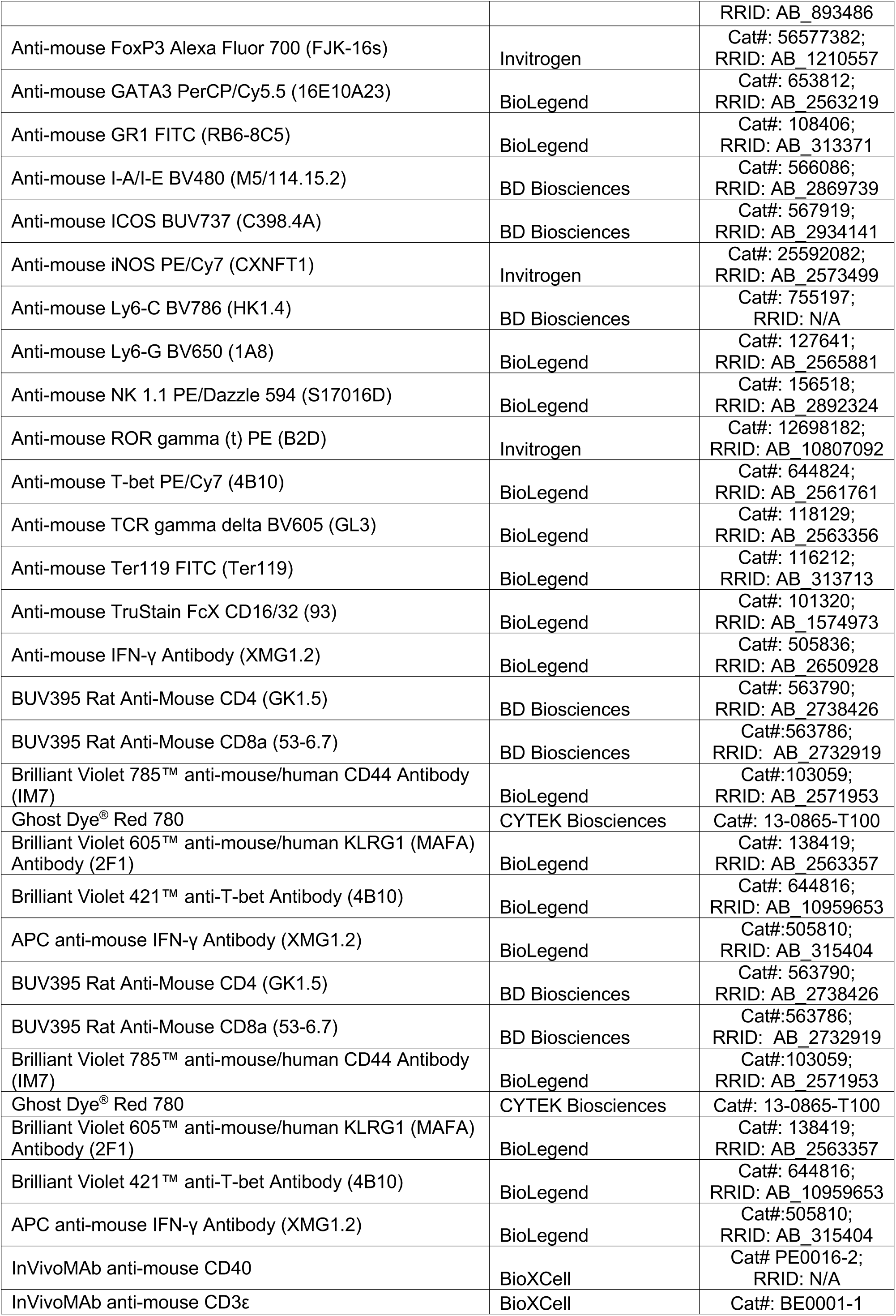
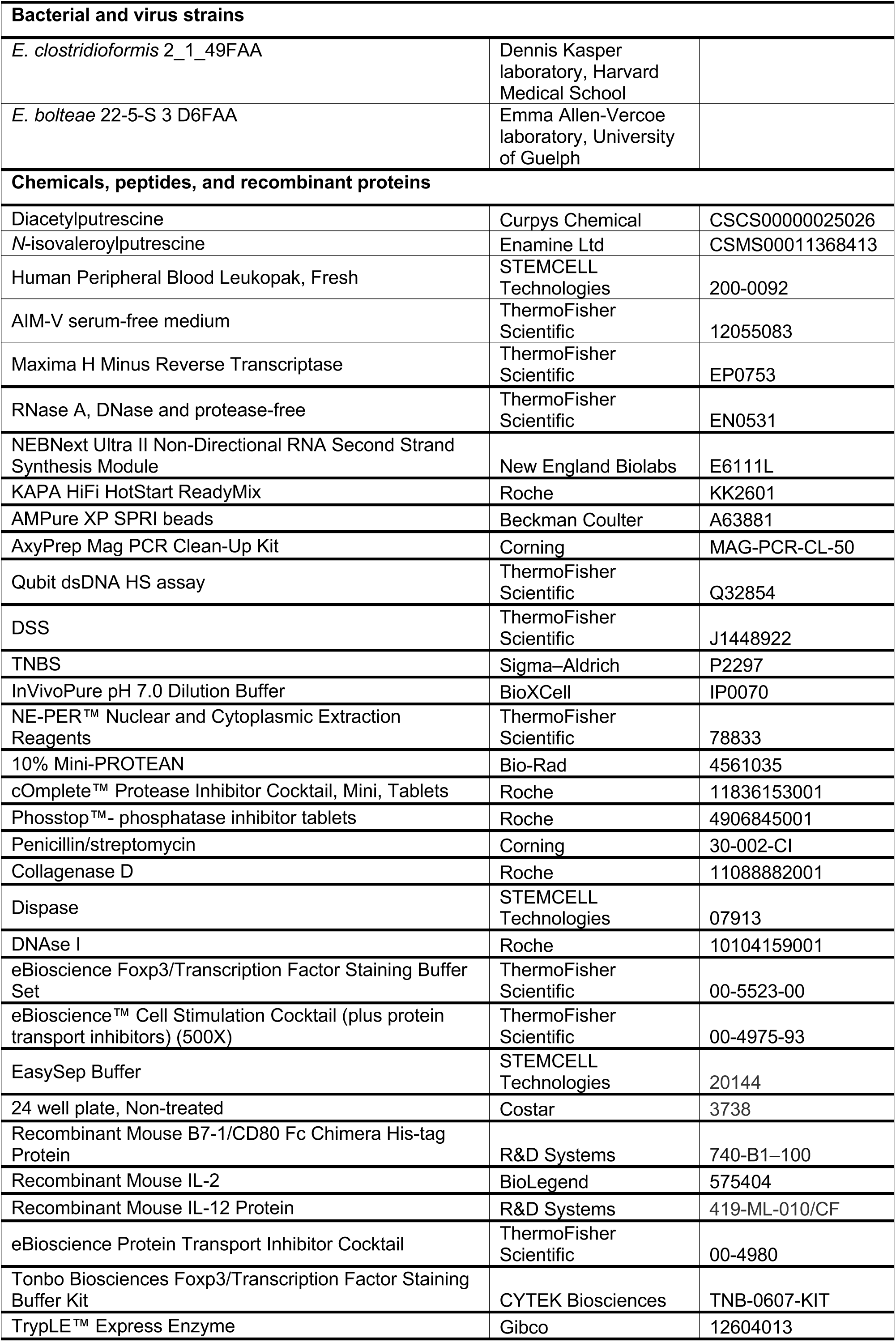
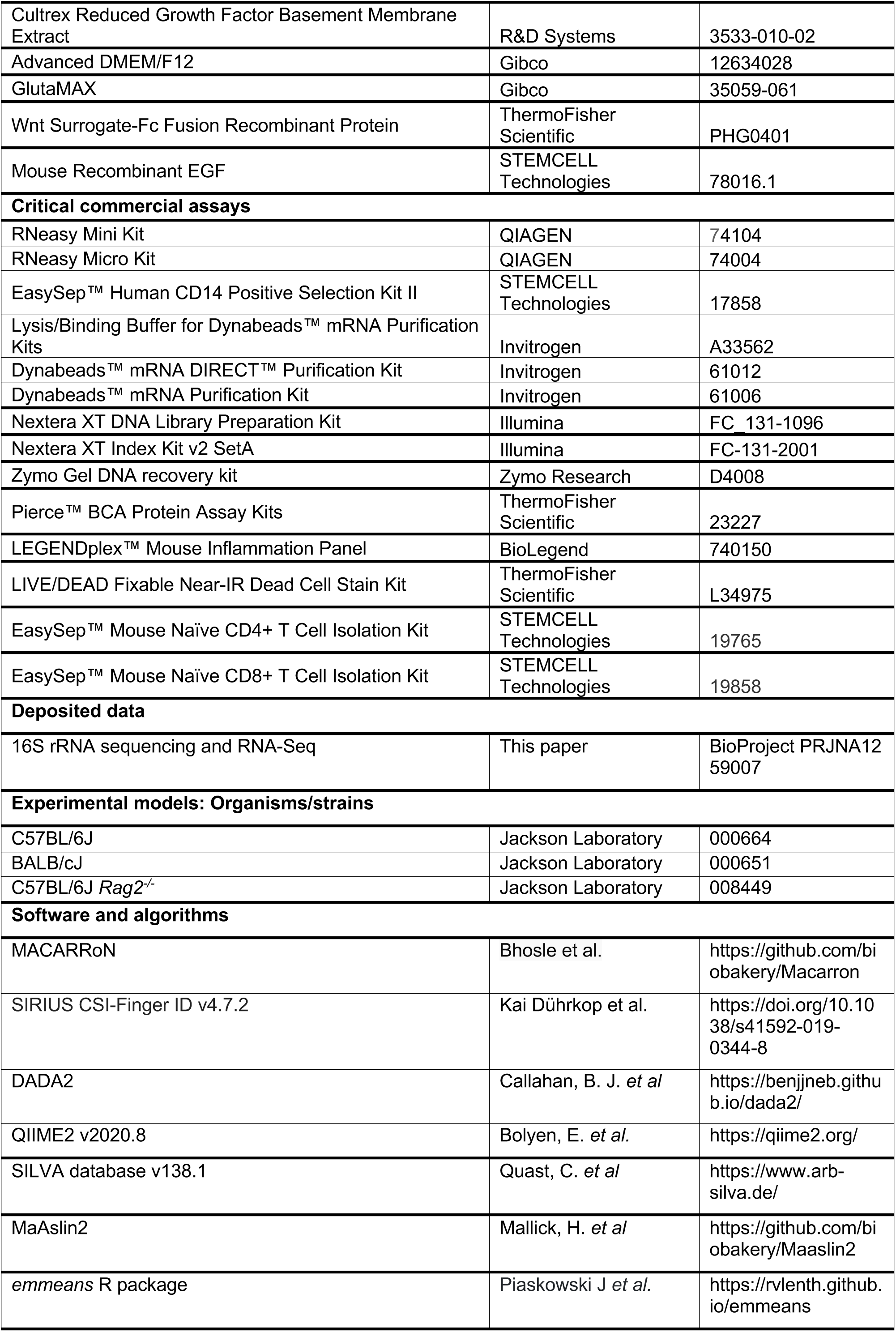
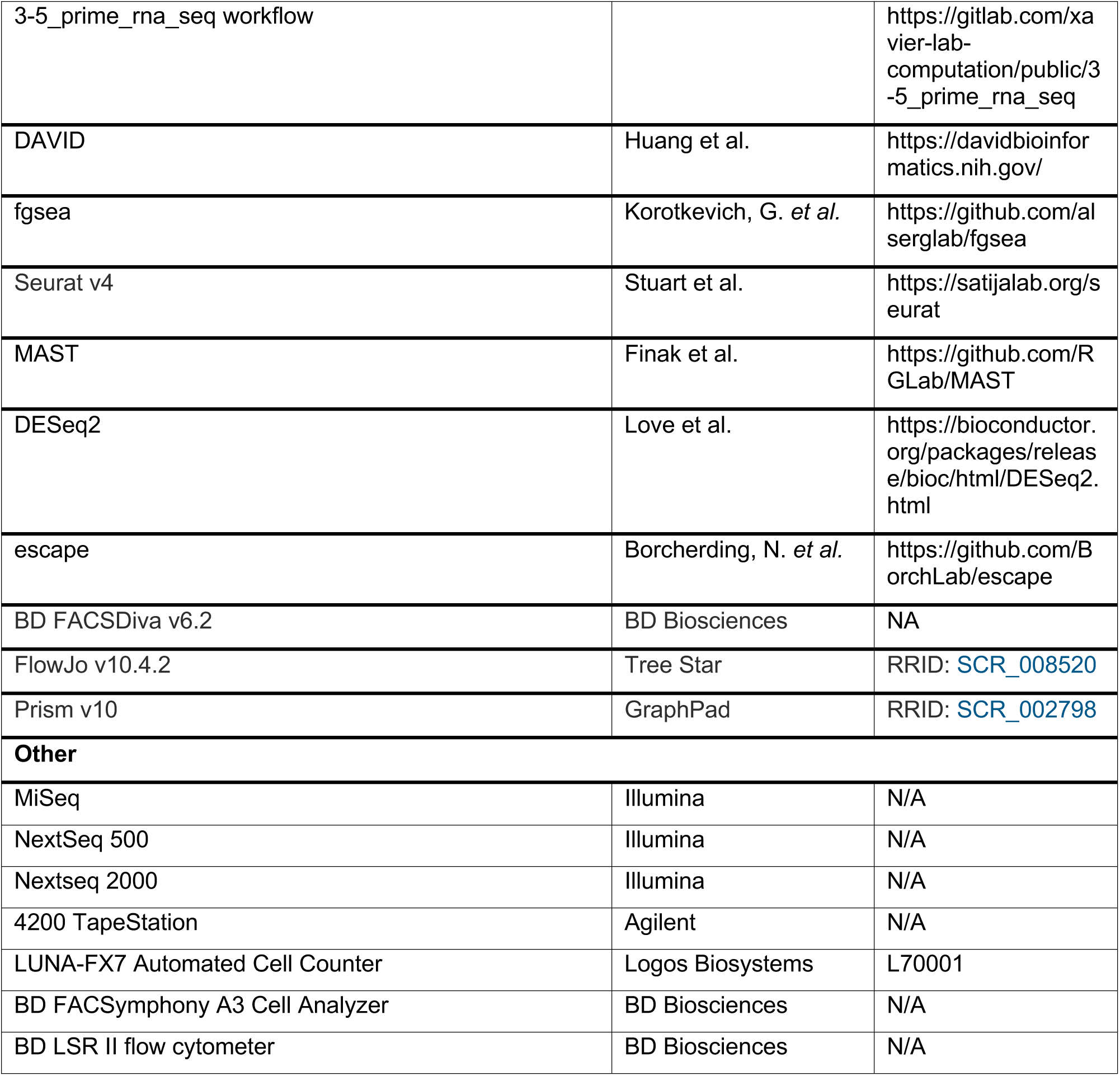

### Experimental model and subject details

#### Mice

C57BL/6J (Strain #: 000664) mice were purchased at 6 weeks of age from Jackson Laboratory were housed in semi-rigid gnotobiotic isolators in the Harvard T. H. Chan Gnotobiotic Center for Mechanistic Microbiome Studies.

Germ-free WT C57BL/6J mice were bred and maintained in semi-rigid gnotobiotic isolators in the Harvard T. H. Chan Gnotobiotic Center for Mechanistic Microbiome Studies.

BALB/cJ (Strain #: 000651) mice purchased at 6 weeks of age from Jackson Laboratory were housed in microisolator cages in the specific-pathogen-free (SPF) barrier facility at the Harvard T.H. Chan of Public Health.

C57BL/6J *Rag2^-/-^* (Strain #: 008449) male mouse^41^ was purchased from Jackson Laboratory, bred with C57BL/6 *Rag2^-/-^* female mice and housed in microisolator cages in the barrier facility at the Harvard T.H. Chan of Public Health. Mice were compared with littermate controls.

Mice were used experimentally between 6-12 weeks of age. Mice were randomized to experimental groups one week prior to the start of an experimental intervention to minimize cage-based or housing bias. Animal studies and experiments were approved and carried out in accordance with Harvard Medical School’s Standing Committee on Animals and the National Institutes of Health guidelines for animal use and care.

Male C57BL/6J mice were used for putrescine or NOP supplementation studies and the DSS colitis model. Female BALB/cJ mice were used for the TNBS colitis mode. Male and female C57BL/6 *Rag2^-/-^*mice were both used in these studies.

Mice used in the gnotobiotic and dextran sulfate sodium-induced colitis experiments, were reared under the same environmental conditions, including housing in semi-rigid gnotobiotic isolators and provided with Inotiv 2020SX Teklad Global Soy Protein-Free Extruded Rodent Diet. BALB/cJ and C57BL/6 *Rag2^-/-^* mice were housed in microisolator cages and provided with LabDiet 5058.

#### Bacterial strains and gnotobiotic colonization

Two human isolates *Enterocloster clostridioformis* 2_1_49FAA, *Enterocloster bolteae* 22-5-S 3 D6 FAA and *Escherichia coli* MG1655 were grown in pre-reduced brain heart infusion (BHI) media in an anaerobic chamber at 37°C. GF C57BL/6 male mice maintained in semi-rigid gnotobiotic isolators were orally gavaged once with ∼10^9^ CFU of *E. clostridioformis* 2_1_49FAA or *E. bolteae* 22-5-S 3 D6 FAA (in 100 μl volume).

### METHOD DETAILS

#### Identifying differentially abundant metabolic features in HMP2 and PRISM

Two publicly available IBD patient untargeted metabolomics datasets (the <u>H</u>uman <u>M</u>icrobiome <u>P</u>roject <u>2</u> (HMP2)^9^ and the <u>P</u>rospective <u>R</u>egistry in IBD <u>S</u>tudy at <u>M</u>GH (PRISM) Crohn’s Disease)^8^ were used for differential abundance (DA) analysis. The HMP2 fecal metabolomic dataset included 546 samples from 106 subjects (CD n=265 from 50 subjects, UC n=146 from 30 subjects, and control individuals without IBD n=135 from 26 subjects) and were downloaded from http://ibdmdb.org/. The PRISM CD (n=68) and non-IBD control (n=24) dataset were taken from the associated published supporting information.^8^

Fecal metabolomics data from the HMP2 (a longitudinal dataset) and PRISM (a cross-sectional dataset) were analyzed separately due to differences in study design. Prior to statistical model fitting, gut metabolome profiles of HMP2 and PRISM participants’ samples were median-normalized to reduce technical sample-to-sample variation and then prevalence-filtered to remove low-confidence features (requiring >30% non-zero values). Metabolite prevalence was computed over individual samples rather than at the subject level and log-transformed for variance stabilization (replacing zero values with half the smallest non-zero measurement on a per-feature basis). We used the same statistical models for HMP2 and PRISM as defined in their original papers.^8,9^ HMP2 includes a separate term for time-variable dysbiosis within diagnosis (based on ecological distance from non-IBD samples); it was fitted in the model, but the coefficient was not used in the work.

Differential abundance across disease phenotypes (CD versus non-IBD and UC versus non-IBD) within the HMP2 cohort was determined by evaluating the following linear mixed-effects model for each metabolite, implemented using the *nlme* package in R. In the model, the following covariates are included: IBD diagnosis as a categorical variable with non-IBD as the reference group; consent age as a continuous covariate; antibiotic exposure as a binary covariate; and subject as a random effect (to account for repeated sampling).

*feature ∼ (intercept) + diagnosis + diagnosis/dysbiosis + antibiotic use + consent age + (1 | subject)*

For transformed abundances of PRISM data, we used a linear model implemented in the R language to identify differentially abundant metabolites in CD. In the model, the following covariates are included: IBD diagnosis as a categorical variable with non-IBD as the reference group; age as a continuous covariate; and four medication exposures (antibiotics, immunosuppressants, mesalamine, steroids) each as a binary covariate.

*feature ∼ (intercept) + diagnosis + antibiotic + immunosuppressant + mesalamine + steroids + age*

The statistical significance (p-value) of metabolite-phenotype associations was assessed using Wald’s test and corrected for multiple hypothesis testing using the Benjamini-Hochberg FDR method.

#### Identifying microbe-associated metabolic features from gnotobiotic mouse stool samples

Stool pellets were collected from germ-free (GF), altered Schaedler Flora (ASF) and specified-pathogen free (SPF) C57BL/6 mice (6-9wks), snap-frozen immediately, and stored at -80°C until untargeted metabolomics sample preparation. The samples were collected at two independent time points. The first set of samples (GF n=6, ASF n=10, and SPF n=12) were analyzed by HILIC-pos mode. The second set (GF n=7, ASF n=8, and SPF n=6) was analyzed using HILIC-neg, C8-pos and C18-neg modes. The processed metabolic features are provided in Supplemental Table 15 and 16 and were used for microbe-associated feature testing.

Prior to statistical model fitting, mouse gut metabolomes were (1) median-normalized to reduce technical sample-to-sample variation; (2) prevalence-filtered to remove low-confidence features (requiring >10% non-zero values); and (3) log-transformed for variance stabilization (replacing zero values with half the smallest non-zero measurement on a per-feature basis).

Differential abundance metabolic features across gut microbial community (GF versus ASF and GF versus SPF) in the mouse fecal metabolomes was determined by evaluating the following linear mixed-effects model for each metabolite, implemented using the *nlme* package in R: gut microbiota type (GF, ASF, and SPF) as a categorical variable with GF as the reference group; sex as a categorical variable; and caging as a random effect.

*feature ∼ (intercept) + gut microbiota type + sex + (1 | cage)*

The statistical significance (p-value) of metabolite-phenotype associations was assessed using Wald’s test and corrected for multiple hypothesis testing using the Benjamini-Hochberg FDR method.

#### Identification of IBD-associated and microbe-associated unknown metabolic features

The identification process consisted of three main steps. 1) *Differential abundance (DA) analysis*: DA analysis was independently performed on the human cohort datasets (HMP2 and PRISM) and the gnotobiotic mouse metabolomics dataset to identify metabolic features associated with IBD and microbial status, respectively, as described above. 2) *Metabolic feature alignment across untargeted metabolomics datasets*: LC-MS features from the human (HMP2 and PRISM) and mouse datasets were aligned by matching peaks based on retention time and m/z values, using the Eclipse tool for feature matching.^42^ 3) *Covariance-based clustering*: Metabolic features from the HMP2 dataset were clustered into abundance covariance-based modules using the findMACMod function from MACARRoN^7^ implementation. At the end of the workflow, each HMP2 metabolic feature was annotated with IBD association statistics, microbial status linkage, and module membership.

We identified 59 modules with at least one DA known metabolite (anchor compound) and one IBD- and microbe-associated unknown feature (Supplemental Table 1, 2). We prioritized modules for further investigation, guided by the following properties: containing at least one DA anchor compound, a minimum of 25 DA microbial-associated unknown features in the module, and containing an anchor compound with prior biological knowledge relevant for IBD epithelial, myeloid, or T cell biology. Three modules met these criteria in both CD and UC, including one (module #11) with metabolic features that were decreased in IBD versus healthy controls and two (module #44 and #122) with numerous features increased in IBD versus healthy controls. These three modules contained stercobilin, glycerophospholipid, and polyamines as anchor compounds. We chose module #122, with the anchor compounds putrescine (P) and *N*-acetylputrescine (NAP) (abundance in HMP2 IBD cohort samples, Extended Data Fig. 1a), for further investigation.

#### Acylated putrescine prediction, synthesis and validation

##### MS/MS data acquisition and prediction of candidate microbe-associated putrescine derivatives

The 29 IBD-associated and microbe-associated unknown metabolic features were prioritized for MS/MS analysis. MS/MS spectra of the candidate features were generated using collision energies ranging from 10 to 50V in 10V increments in a Thermo IDX mass spectrometer (Thermo Fisher Scientific; Waltham, MA) with electrospray ionization in the positive ion mode using full scan analysis at 60,000 resolution followed by five MS-MS scans at 30,000 resolution and an isolation width of +/- 0.2 mass units. Additional MS settings were sheath gas 40, sweep gas 2, spray voltage 3.5 kV, capillary temperature 330°C, S-lens RF 40, heater temperature 300C, microscans 1, automatic gain control target 1e6, and maximum ion time 250 ms. Parsed MS-MS data (*.ms) were loaded into SIRIUS CSI-Finger ID version 4.7.2 ^43^. Molecular formula predictions generated with Orbitrap-specific settings (MS-MS isotope scorer: ignore, mass deviation: 5ppm, Candidates: 10, Candidates per ion: 1, possible ionizations: [M + H]+, [M + K]+, [M+Na]+). Structure elucidations were done using all included databases and the adducts [M + H]+, [M + K]+, [M + Na]+). The predictions were exported, and the top three structure annotations for each feature were evaluated. *N*-isovaleroyl P (N-isoVP) was identified among the predicted structures and validated using a commercial standard.

Concordant with our N-isoVP prediction, prior studies have demonstrated that the human gut microbiome produces fatty acid amide conjugates with host bioactive potential.^44,45^ Thus, we generated an *in silico* database of all possible combinations of fatty acid ranging from 2 to 40 carbons and P or NAP conjugates and matched their predicted *m/z* to the unknown stool features. This approach led to the prediction of three additional unknown features, *N*-propionoyl P (NPP), *N*-oleoyl P (NOP) and *N*-stearoyl P (NSP) in module #122.

##### Acylated putrescine validation experiment

Acylated putrescines were validated using authentic reference standards. The two commercially available standards are as follows: DAP - Cat#: CSCS00000025026, Curpys Chemical and N-isoVP – Cat#: CSMS00011368413, Enamine Ltd. Acylated putrescines were analyzed alongside reference materials from the HMP2, PRISM, and mouse experiments using the HILIC-positive LC-MS method described above. Retention times (RT) and m/z values of the standards were matched to those of the corresponding unknown features. Subsequently, MS/MS spectra were acquired for both the standards and the matched features in each biofluid at five collision energies (10 to 50 eV). The resulting fragment ion patterns were compared and used to further confirm the identity of the unknowns.

##### Acylated putrescine profiling in human gut microbiota depletion intervention study

Acylated putrescines were further assessed in a publicly available longitudinal human fecal metabolomics dataset from the Food and Resulting Microbial Metabolites (FARMM)^21^ study (Metabolomics Workbench project ID: PR001024), which includes an antibiotic and polyethylene glycol intervention. Acylated putrescines annotation in the FARMM study was performed using LC-MS analysis on a Thermo Q Exactive Orbitrap operated in positive ion mode, followed by feature alignment and annotation based on m/z and RT.

##### Validation of human microbiome-based filtering approach

Fecal metabolomics data acquired in HILIC-pos mode from FARMM^21^ longitudinal samples were downloaded from Metabolomics Workbench (project ID: PR001024). To identify microbe-associated metabolic features, paired pre- and post-microbiota depleted samples from diet-controlled inpatient groups (omnivore diet n=10 or exclusive enteral nutrition (EEN) diet n=10) were selected for downstream analysis to minimize environmental and dietary variability. Prior to statistical model fitting, the metabolomics data were median normalized and log transformed (with pseudo count 1 for zero values) and then fit with the following per-feature linear mixed effects model:

*feature ∼ (intercept) + antibiotic and polyethylene glycol treatment + sex + age + (1 | subject)*

The model included antibiotic and polyethylene glycol treatment and sex as categorical fixed effects, age as a continuous covariate and subjects as a random effect and were implemented using the *nlme* package in R. Model fitting was performed separately within each diet group (omnivore and EEN). Statistical significance (p-value) of metabolite-phenotype associations were assessed using Wald’s test and corrected for multiple hypothesis testing using the Benjamini-Hochberg FDR method. Microbe-associated metabolic features were defined as those significantly depleted (FDR-adjusted *p*-value < 0.2) following antibiotic and polyethylene glycol treatment.

##### Assessment of gut microbiota-dependent transformation of putrescine

Germ-Free (GF) (control n=6, putrescine n=6), altered Schaedler Flora (ASF) (control n=10, putrescine n=9), and SPF (control n=12, putrescine n=12) C57BL/6 mice were housed in the same environments and provided with drinking water with or without 1% w/v putrescine-dihydrochloride (Cat #: P5780, Sigma-Aldrich) for three weeks. Stool pellets (30-100mg) were collected at the end of the treatment, snap-frozen immediately, and stored at -80°C until sample submission for metabolomics.

SPF mice came from two separate rooms in the Jackson vivarium. For all subsequent experiments using SPF C57BL/6 mice (see below) in this study, mice were imported from the same, second room. Relative abundance levels of acylated putrescines were analyzed by HILIC-pos mode

##### Untargeted metabolomics sample preparation and mass spectrometry analysis

For the putrescine untreated group, a combination of four LC-MS methods was used to profile metabolites in the fecal homogenates, as previously described; two methods that measure polar metabolites, a method that measures metabolites of intermediate polarity (e.g., fatty acids and bile acids), and a lipid profiling method. For the putrescine treatment group, HILIC-positive method (positive ion mode MS analyses of polar metabolites) was used to profile fecal metabolites in the fecal homogenates. Snap-frozen stool samples were thawed on ice, and then aqueous homogenates were generated by homogenizing samples in 10 volumes of water (mg: µL) using a TissueLyser II (QIAGEN) bead mill set to two 2 min intervals at 20Hz. The homogenate for each sample was divided into two 10 µL and two 30 µL aliquots in 1.5mL centrifuge tubes for LC-MS sample preparation. 30 µL of homogenate from each sample was transferred into a 50 mL conical tube on ice to create a pooled reference sample. Subjects were randomized in the analysis queue in each method. Additionally, pairs of pooled reference samples were inserted into the queue at intervals of approximately 20 samples for quality control and data standardization. Samples were prepared for each method using extraction procedures that are matched for use with the chromatography conditions. Data were acquired using LC-MS systems comprised of Nexera X2 U-HPLC systems (Shimadzu Scientific Instruments) coupled to Q Exactive/Exactive Plus orbitrap spectrometers (ThermoFisher Scientific). The method details are summarized below.

###### LC-MS Method 1: HILIC-pos (positive ion mode MS analyses of polar metabolites)

LC-MS samples were prepared from stool homogenates (10 µL) by protein precipitation with the addition of nine volumes of 74.9:24.9:0.2 v/v/v acetonitrile/methanol/formic acid containing stable isotope-labeled internal standards (valine-d8, Isotec; and phenylalanine-d8, Cambridge Isotope Laboratories). The samples were centrifuged (10 min, 9,000g, 4°C), and the supernatants injected directly onto a 150 x 2-mm Atlantis HILIC column (Waters). The column was eluted isocratically at a flow rate of 250 µL/min with 5% mobile phase A (10 mM ammonium formate and 0.1% formic acid in water) for 1 min followed by a linear gradient to 40% mobile phase B (acetonitrile with 0.1% formic acid) over 10 min. MS analyses were carried out using electrospray ionization in the positive ion mode using full scan analysis over m/z 70–800 at 70,000 resolution and 3-Hz data acquisition rate. Additional MS settings are: ion spray voltage, 3.5 kV; capillary temperature, 350°C; probe heater temperature, 300°C; sheath gas, 40; auxiliary gas, 15; and S-lens RF level 40.

###### LC-MS Method 2: HILIC-neg (negative ion mode MS analysis of polar metabolites)

LC-MS samples were prepared from stool homogenates (30 µL) by protein precipitation with the addition of four volumes of 80% methanol containing inosine-15N4, thymine-d4 and glycocholate-d4 internal standards (Cambridge Isotope Laboratories). The samples were centrifuged (10 min, 9,000g, 4°C) and the supernatants were injected directly onto a 150 x 2.0-mm Luna NH2 column (Phenomenex). The column was eluted at a flow rate of 400 µL/min with initial conditions of 10% mobile phase A (20 mM ammonium acetate and 20 mM ammonium hydroxide in water) and 90% mobile phase B (10 mM ammonium hydroxide in 75:25 v/v acetonitrile/methanol) followed by a 10-min linear gradient to 100% mobile phase A. MS analyses were carried out using electrospray ionization in the negative ion mode using full scan analysis over m/z 60–750 at 70,000 resolution and 3 Hz data acquisition rate. Additional MS settings are as follows: ion spray voltage, -3.0 kV; capillary temperature, 350°C; probe heater temperature, 325°C; sheath gas, 55; auxiliary gas, 10; and S-lens RF level 40.

###### LC-MS Method 3: C18-neg (negative ion mode analysis of metabolites of intermediate polarity; for example, bile acids and free fatty acids)

Stool homogenates (30 µL) were extracted using 90 µL methanol containing 15R-15-methyl ProstaglandinA2, 15S-15-metyl ProstaglandinE1, 15S-15-metyl ProstaglandinE2, as internal standards (Cayman Chemical Co.) and centrifuged (10 min, 9,000g, 4°C). The supernatants (10 µL) were injected onto a 150 x 2.1-mm ACQUITY BEH C18 column (Waters). The column was eluted isocratically at a flow rate of 450 µL/min with 20% mobile phase A (0.01% formic acid in water) for 3 min followed by a linear gradient to 100% mobile phase B (0.01% acetic acid in acetonitrile) over 12 min. MS analyses were carried out using electrospray ionization in the negative ion mode using full scan analysis over m/z 70–850 at 70,000 resolution and 3 Hz data acquisition rate. Additional MS settings are as follows: ion spray voltage, -3.5 kV; capillary temperature, 320°C; probe heater temperature, 300°C; sheath gas, 45; auxiliary gas, 10; and S-lens RF level 60.

###### LC-MS Method 4: C8-pos

Lipids (polar and nonpolar) were extracted from stool homogenates (10 µL) using 190 µL isopropanol containing 1-dodecanoyl-2-tridecanoyl-sn-glycero-3-phosphocholine as an internal standard (Avanti Polar Lipids; Alabaster, AL). After centrifugation (10 min, 9,000g, ambient temperature), supernatants (10 µL) were injected directly onto a 100 x 2.1-mm ACQUITY BEH C8 column (1.7 mm; Waters). The column was eluted at a flow rate of 450 mL/min isocratically for 1 min at 80% mobile phase A (95:5:0.1 v/v/vl 10 mM ammonium acetate/methanol/acetic acid), followed by a linear gradient to 80% mobile phase B (99.9:0.1 v/v methanol/acetic acid) over 2 min, a linear gradient to 100% mobile phase B over 7 min, and then 3 min at 100% mobile phase B. MS analyses were carried out using electrospray ionization in the positive ion mode using full scan analysis over m/z 200–1,100 at 70,000 resolution and 3 Hz data acquisition rate. Additional MS settings are as follows: ion spray voltage, 3.0 kV; capillary temperature, 300°C; probe heater temperature, 300°C; sheath gas, 50; auxiliary gas, 15; and S-lens RF level 60.

##### Untargeted metabolomics data processing

Raw LC-MS data were acquired using the data acquisition computer interfaced to each LC-MS system and then stored on a robust and redundant file storage system (Isilon Systems) accessed via the internal network at the Broad Institute. untargeted data were processed using Progenesis QIsoftware (v 2.0, Nonlinear Dynamics) to detect and de-isotope peaks, perform chromatographic retention time alignment, and integrate peak areas. To remove redundant adducts and peaks, LC-MS peaks were clustered based on intensities and retention times. Clusters were generated containing peaks occurring within a retention time window of 0.025 min (0.015 min for C8-pos) and with intensities correlating with Spearman rank correlation coefficients above 0.8 and 0.85 for negative mode and positive methods respectively. The peak with the highest mean abundance within each cluster was kept as the representative ion and all other cluster members removed from the final dataset. Un-clustered peaks (singlets) were kept in the final dataset. Peaks of unknown identity were tracked by method, m/z and retention time. Identification of the resulting 48,557 untargeted metabolite LC-MS peaks, from the four LC-MS methods, was conducted by: i) matching measured retention times and masses to mixtures of reference metabolites analyzed in each batch; and ii) matching an internal database containing thousands of compounds that have been characterized using the Broad Institute methods, yielding 910 peaks with a match. Temporal drift was monitored and normalized with the intensities of peaks measured in the pooled reference samples.

##### 16S ribosomal RNA (rRNA) gene amplicon sequencing of mouse stool DNA

The 16S rRNA gene amplification protocol was adapted from the Earth Microbiome Project ^46^. The 16S rRNA V4 region of extracted DNA was amplified by PCR and then purified by the AxyPrep Mag PCR Clean-Up Kit (Cat#: MAG-PCR-CL-50, Corning) to remove free primers and primer dimers. The purified amplicon was quantified using a Qubit dsDNA HS assay (Cat#: Q32854, ThermoScientific Fisher) and an equal amount (by mass) was pooled together. The 16S rRNA V4 library sequencing was performed on a MiSeq instrument (Illumina, San Diego, CA) using 250bp paired-end reading by GENEWIZ from Azenta Life Science. The demultiplexed raw sequencing data were imported to the QIIME2 environment (version 2020.8)^47^ then low-quality bases were removed. The quality trimmed reads were joined, denoised, and checked for chimeras using DADA2 plug-in prior to taxonomic assignment^48^. Taxonomic assignment of each amplicon sequence variant (ASV) was performed using a pre-trained Naive Bayes classifier with the SILVA database (version 138.1).^49^ The feature table was further used for differential abundance analysis using MaAslin2.^50^ In this analysis, the differential abundance testing focused on the genus-level due to the resolution of 16S rRNA V4 region; low prevalence genera found in less than 10% of all samples, and genera that were low abundance (average relative abundance below 0.001%) were not included in the testing. Each genus-level was modeled as a function of treatment (categorical variable) with caging as a random effect. Genus levels with a corrected q-value of less than 0.25 are considered significant.

##### Mouse bone marrow-derived dendritic cell (BMDC) culture and compound treatment

Mouse BMDCs were generated from the bone marrow of C57BL/6 wild-type mice purchased from Jackson Laboratories. Bone marrow cells were isolated, red blood cell depleted via a hypotonic saline treatment, and pelleted cells were then resuspended and cultured in RPMI-1640 medium supplemented with 10% fetal bovine serum (FBS), 1% penicillin/streptomycin, and 40 ng/mL murine GM-CSF at 37 °C in a humidified atmosphere with 5% CO₂ for 7 days. On day 7, differentiated cells were gently scraped from the culture dishes, counted, and seeded into 96-well plates at a density of 100,000 cells per well in DMEM containing 10% FBS and 1% penicillin/streptomycin. Cells were allowed to adhere overnight under standard culture conditions. The following day, media was replaced with AIM-V serum-free medium (Cat#: 12055083, ThermoFisher Scientific) containing either 1%, 0.1%, or 0.01% (v/v) DMSO or polyamines/acylated putrescines at final concentrations of 100 µM, 10 µM, or 1 µM, respectively. For the 3′-DGE screening assays, cells were treated with the compounds for 4 hours at 37 °C and 5% CO₂. For full-length RNA-sequencing screening, cells were treated with either DMSO (vehicle control) or NOP for 6 hours under the same conditions. Following treatment, media was aspirated, cells were rinsed once with PBS, lysed in 50 µL Dynabead lysis buffer, and stored at −80 °C until RNA sequencing library construction.

##### Mouse colonic organoid culture and compound treatment

Mouse colonic crypts were isolated from C57BL/6 wild-type mice as previously described.^51^ Organoid culture and passaging were performed following a well-established protocol;^52^ For passaging, organoids were dissociated by incubation with TrypLE (Gibco) and embedded in basement membrane matrix (Matrigel; R&D Systems).

To evaluate the transcriptional effects of compounds, organoids were first cultured in L-WRN conditioned medium supplemented with 10% FBS and 1% penicillin/streptomycin for 3 days. Prior to compound treatment, the media was aspirated, organoids were rinsed with PBS once and then replaced with serum free media containing Advanced DMEM/F12 (Gibco, 12634028), 1% GlutaMAX (Gibco, 35059-061), Wnt Surrogate (200ng/ml; Thermo Scientific), mouse EGF (STEMCELL Technologies) 1% penicillin/streptomycin, and 10% condition medium containing R-spondin1 and Noggin. Organoids were then treated with vehicle control, polyamines, or acylated putrescines at final concentration of 100 µM, 10 µM, or 1 µM, respectively, for 24 hours at 37 °C and 5% CO₂. Following treatment, culture media was aspirated, cells were rinsed once with PBS. Total RNA was extracted using RNeasy Mini Kit (Qiagen, 74104) according to the manufacturer’s instructions and stored at −80 °C until RNA sequencing library preparation.

##### Identification of microbial features associated with NOP production

To identify microbial features associated with NOP production, paired metagenomic and metabolomic samples (n=461) from the HMP2 dataset were analyzed. Associations between microbial species and NOP abundance were assessed by calculating Spearman correlations between residual microbial species abundances and residual NOP abundances. Microbial species having significant positive correlations with NOP abundance were considered as NOP producing candidates. To investigate the potential biosynthetic basis of NOP production, we evaluated the association between NOP abundance and nonribosomal peptide synthetases (NRPS), which are enzymatic components of microbial biosynthetic gene clusters (BGCs) and are known to mediate fatty acid amide conjugation. Using the curated non-redundant core NRPS dataset reported by Elmassry et al.,^10^ Spearman correlations were computed between residual core NRPS abundances and residual NOP abundances in the HMP2 dataset. Residual values for both microbial features and NOP were derived from the linear mixed effects model described above for the HMP2 cohort. Residual values were used for the correlation analysis to identify associations that are independent of the model covariates by subtracting variation attributable to diagnosis, dysbiosis status, age, antibiotic usage, sample collection site and per-subject variation. Spearman correlation coefficients were assessed using two-tailed tests, and p-values were adjusted for multiple hypothesis testing using false discovery rate correction.

##### In vitro evaluation of NOP production

To evaluate microbial NOP production, in vitro assays were performed using two human isolates *Enterocloster clostridioformis* 2_1_49FAA and *Enterocloster bolteae* 22-5-S 3 D6 FAA. *Escherichia coli* MG1655 which lacks NRPS biosynthetic gene clusters was used as a control. The substrate preparation and *in vitro* assay were adapted from previously published methods.^19,53^ Putrescine was prepared as a 50mM stock solution in water, and labeled oleic acid ([U-^13^C_18_] OA) was prepared as a 50mM stock solution in ethanol and neutralized with sodium hydroxide solution at an equimolar (1:1) ratio of oleic acid to sodium hydroxide. Both stock solutions were stored at −80 °C. *E. clostridioformis* 2_1_49FAA, *E. bolteae* 22-5-S 3 D6 FAA and *E. coli* MG1655 were individually inoculated into pre-reduced brain heart infusion (BHI) media and were grown overnight in an anaerobic chamber at 37°C.

Overnight cultures were normalized to an OD_600_ = 0.1 by dilution into fresh pre-reduced BHI media. Bacterial cultures were supplemented with putrescine (final concentration 100 μM) and labeled oleic acid (final concentration 100 μM) and incubated anaerobically at 37°C for 48hrs. Bacterial cultures were shielded from light throughout the incubation period to protect against light-induced degradation of the labeled oleic acid substrate. Following incubation, bacterial cultures were serially diluted and plated for colony forming unit (CFU) enumeration. The remaining bacterial cultures were maintained on ice during harvest and centrifugation. Cell pellets were prepared by centrifugation at 20,000g for 1min at 4°C. Cell pellets were washed with ice-cold PBS and residual PBS was removed. Cell pellets were immediately frozen and stored at -80°C until sample submission for metabolomics.

##### In vivo evaluation of NOP production

GF C57BL/6 male mice maintained in semi-rigid gnotobiotic isolators were orally gavaged once with ∼10^9^ CFU of *E. clostridioformis* 2_1_49FAA or *E. bolteae* 22-5-S 3 D6 FAA (in 100 μl volume). Following oral gavage, mice were provided with drinking water with or without 1% w/v putrescine dihydrochloride for two weeks. Mice were maintained in semi-rigid gnotobiotic isolators during the experimental period. At the end of the treatment period, stool pellets were collected, immediately snap-frozen, and stored at -80°C until sample submission for metabolomics. Mono-colonization was confirmed by quantitative PCR. *E. clostridioformis* 2_1_49FAA, *E. bolteae* 22-5-S 3 D6 FAA cultures for inoculation of the mice were prepared as described above. Statistical analysis of the *in vivo* assays was performed using a linear model that include experiment batch as additional fixed effect to account for baseline difference in NOP abundance arising from technical variation. Estimated marginal means and pairwise contrasts were calculated using the *emmeans* R package.

##### Human Peripheral Blood Mononuclear Cell (PBMC)-Derived Monocyte Culture and NOP Treatment

Monocytes were isolated from human PBMCs from two healthy donors using the Human CD14 Positive Selection Kit II (STEMCELL Technologies) and cultured in AIM-V serum-free medium (ThermoFisher Scientific) overnight at 37 °C and 5% CO₂ to allow adherence. The purity of isolated monocytes, measured by flow cytometry, was 94% for donor 1 and 86% for donor 2. After an overnight recovery period, cells were treated with NOP at a final concentration of 10 µM, vehicle control, or lipopolysaccharide (LPS, 100 ng/mL; InvivoGen, tlrl-pb5lps) for 6 hours at 37 °C and 5% CO₂. After treatment, the media was aspirated, cells were rinsed once with PBS, and RNA was extracted using the RNeasy Micro Kit (QIAGEN). RNA samples were stored at −80 °C prior to full-length RNA-sequencing library preparation.

##### 3’ Digital Gene Expression (3’-DGE) RNA-sequencing library preparation

High throughput digital gene expression was performed based on modification of the 3’DGE protocol.^54^ After treatment, cells were lysed in Lysis Buffer (Invitrogen), to achieve 100,000 cells/100µL lysis buffer. A volume of 50µL lysate from each sample was used for mRNA purification with the Dynabeads mRNA Direct Purification kit (Invitrogen). Subsequently, reverse transcription was performed with PolydT primers containing unique sample barcodes for each well. Reactions were performed in 20µL reaction mix (8.25µL of mRNA, 0.5µL of 20mM dNTPs, 4µL of 5M Betaine 0.25µL of Maxima H Minus Reverse Transcriptase (ThermoFisher Scientific), 4µL of 5x buffer, 2µL of H2O, and 1µL of 5µM unique PolydT Primer) under the following conditions: 50°C for 30 min then 42°C for 60 min and final heat inactivate at 85°C for 5 min. The generated 1st strand cDNAs were pooled and treated with RNase A (ThermoFisher Scientific), followed by 2nd Strand cDNA synthesis with the NEBNext Ultra II Non-Directional RNA Second Strand Synthesis Module (New England Biolabs) following the kit instructions. The ds-cDNA products were cleaned up with the AMPure XP Reagent SPRI Beads (Beckman Coulter) and quantified and quality checked with a Qubit and 4200 Tapestation, respectively. Subsequently, the ds-cDNA was tagmented with the Nextera XT DNA Library Preparation Kit (Illumina) and amplified by limited cycle PCR using the universal P5 primer and unique P7 index primers. PCR products were gel purified to enrich the library for 300bp-600bp fragments. The final libraries were sequenced on the Illumina Nextseq 2000 with the paired end reads.

##### Full-length RNA-Sequencing library preparation

RNA-seq libraries were prepared according to a modification of the SmartSeq2 protocol.^55^ Briefly, cells were lysed in lysis/binding buffer (Invitrogen) to achieve 10^5^-10^6^ cells in 50-100μl lysis buffer. Subsequently, mRNA was isolated with Dynabeads mRNA Direct Purification kit (Invitrogen). RNA was reverse-transcribed with Maxima RT RNaseH-minus (ThermoFisher Scientific), and cDNA was amplified with KAPA HiFi HotStart Ready Mix (Roche) according to manufacturer instructions. After post-clean up with 0.8x AMPure XP SPRI beads (Beckman Coulter), the DNA was tagmented with Nextera XT DNA sample PreKit (Illumina) and Nextera XT Index Kit v2 SetA (Illumina). SPRI bead clean-up and gel purification were performed to extract and recover library fragments (250-800bp) with Zymo Gel DNA recovery kit (Zymo Research). The library was sequenced on an Illumina NextSeq 500 system.

##### Differential Gene Expression Analysis and GSEA

For both 3’-DGE sequencing and SMART-Sequencing data, raw sequence reads were processed using the 3-5_prime_rna_seq workflow (https://gitlab.com/xavier-lab-computation/public/3-5_prime_rna_seq). Briefly, we used the Illumina bcl2fastq conversion tool (version 2.19.0.316) to convert BCL files into FASTQ format. Reads were demultiplexed using unique molecular identifiers (UMIs)^56^. Genome alignment was performed using the Burrows-Wheeler Aligner (BWA)^57^ against the mm10/GRCm38 (mouse) reference genome. Aligned reads were assigned to RefSeq transcripts and then collapsed by UMI to quantify transcript counts. Finally, RefSeq IDs were mapped to gene symbols, and a gene-by-sample count matrix was generated based on UMI-collapsed counts.

Differential gene expression analysis was performed using DESeq2 version 1.44.0 with default parameters.^58^ Differentially expressed genes (false discovery rate (FDR) < 0.2 and abs (log2 fold change) > 1.5) were used for functional annotation and enrichment analysis using the Database for Annotation, Visualization, and Integrated Discovery (DAVID). To assess overall immune-related pathway-level transcriptional changes, Hallmark pathway^59^ enrichment analysis was performed using the fgsea R package.^60^ The hallmark gene sets were downloaded from the Molecular Signatures Database (MSigDB). The statistical significance of pathway enrichment was determined using normalized enrichment scores (NES). Pathways with an FDR < 0.05 were considered as differentially expressed pathways.

##### Differential HMP2 host gene expression analysis

Raw count matrices of the HMP2 host transcriptomic data were downloaded from the IBDMDB website (https://ibdmdb.org). Inflamed biopsy samples from patients with CD and UC, as well as uninflamed biopsy samples from non-IBD controls, were selected for differential expression analysis. Differential gene expression analysis was conducted using DESeq2 (v.1.44.0) incorporating IBD phenotype and biopsy location as covariates in the regression model. Gene set enrichment analysis was performed using fgsea^61^ package with the MSigDB Hallmark gene sets,^62^ following the same pipeline described above.

##### Single-Cell RNA-Sequencing (scRNA-seq) Analysis of IBD Cohorts

Three publicly available IBD cohort scRNA-seq datasets were obtained: (1) Kong et al.^32^ immune cell compartment data, downloaded from the Broad Single Cell Portal (accession ID: SCP1884), (2) Thomas et al.,^34^ pre-processed myeloid cell compartment data, downloaded from Zenodo (https://zenodo.org/records/14007626) and (3) Smillie et al.,^33^ immune cell compartment data, downloaded from the Broad Single Cell Portal (accession ID: SCP259).

For the Kong et al. and Smillie et al. datasets, pre-defined immune cell annotations from the original publications were used to identify dendritic cells (DC1, DC2) and inflammatory monocytes. Specifically, in Kong et al., cells annotated as DC1 were used as DC1, and cells annotated as DC2 CD1d^-^ and DC2 CD1d^+^ were combined to define the DC2 population. Cells annotated as monocytes S100A8^+^ and S100A9^+^ were defined as inflammatory monocytes. For the Smillie et al. dataset, cells annotated as DC1, DC2, and inflammatory monocytes were extracted for downstream analysis.

For the Thomas et al. dataset, the myeloid cells data were processed using Seurat^63^ (v. 4.3.0) in R (v 4.4.2). Briefly, UMI counts were log-normalized, 3000 most highly variable genes were selected. Data was scaled before principal component analysis (PCA). To correct for inter-patient batch effects, Harmony was applied using patient identity as the grouping variable. Clustering was performed on the Harmony-corrected embeddings using a shared nearest neighbor graph (20 dimensions, k = 50) and community detection at a resolution of 0.8. Clusters were annotated with supervised approach, using the marker genes that used in the original publication.^34^ Marker genes for the DCs and inflammatory monocytes clusters are provided in Supplemental Table 14.

Differential gene expression (DE) analysis was performed using MAST.^64^ Only inflamed tissue samples from IBD patients and noninflamed tissue samples from non-IBD subjects were used for DE analysis. The gene expression for each gene was fitted with a fixed effect model in MAST, to captures effects of disease state on gene expression while controlling for cell complexity (e.g., the number of genes detected per cell). P-values were obtained from likelihood ratio tests and FDR-corrected p-values < 0.2 were considered significant.

##### Derivation of inflammation and IBD therapy nonresponse gene expression scores

Inflammation score and IBD therapy nonresponse gene expression enrichment score were calculated as composite gene signature scores. Differentially expressed genes identified between histologically inflamed IBD resection and noninflamed/non-IBD gut tissues, as reported by Thomas et al.^34^ were used for inflammation score calculation. Gene signatures significantly enriched in inflamed IBD tissue and significantly associated with IBD therapy nonresponse, as reported by Friedrich et al.^35^ were used to compute enrichment scores representing IBD therapy nonresponse-associated gene co-expression signature. Mouse BMDC and human PBMC-derived monocytes transcriptome datasets were normalized by estimating size factors with DESeq2,^65^ followed by variance stabilizing transformation respectively. Gene signature enrichment analysis was performed using *enrichIt* function in the *escape* R package^66^ to calculate per sample enrichment scores for each gene signature. The enrichment scores were scaled between 0-10 for statistical analysis as in Thomas et al..^34^ Statistical analysis of the monocyte data was performed using a linear model with NOP treatment and donor as fixed effects to account for donor-to-donor variation. Estimated marginal means and pairwise contrast were calculated using *emmeans* R package.

#### Mouse models of Intestinal Inflammation and Related Experimental Methods

##### Dextran Sulfate Sodium (DSS) colitis model

###### Putrescine treatment

C57BL/6 GF and SPF male mice were reared in the same environmental conditions and received DSS (2.5% w/v for GF, 3% w/v for SPF, given morbidity and mortality issues of GF mice) in the drinking water for 5 days and followed by regular drinking water for 2 days. Putrescine (1% w/v) was supplemented to the drinking water throughout experiments, and the sham control group did not receive putrescine. Body weight was measured daily, and mice were euthanized on day 7 post DSS administration. Upon sacrifice, the colon was harvested, cleaned with PBS and then fixed with 4% paraformaldehyde (PFA) for histologic assessment.

###### N-oleoylputrescine treatment

Two different NOP treatment models were employed in the DSS-induced colitis model.

1. <u>Therapeutic DSS model</u>: C57BL/6J SPF male mice were given 3% w/v DSS in the drinking water for 5 days followed by regular drinking water for 7 days. 24mg/kg NOP or the same volume of H_2_O was orally gavaged starting on day 5 and continuing through day 11; oral gavage and daily body weight measurement were performed at approximately the same time (±2 hours). Upon sacrifice, the colon was harvested and then fixed with 4% PFA for histologic assessment.
2. <u>Prevention DSS model</u>: C57BL/6J SPF male mice received 3% w/v DSS in the drinking water for 5 days followed by regular drinking water for 2 days. 24mg/kg NOP or an equivalent volume of H_2_O was orally gavaged beginning 3 days prior to DSS administration (day -3) and continuing through day 7; oral gavage was done at approximately the same time each day (±2 hours) along with body weight measurement. Upon sacrifice, half of the longitudinally sectioned colon was fixed with 4% PFA for histologic assessment and the second half of the colon was used for immune cell profiling by flow cytometry. In addition, blood samples were collected via cardiac puncture into BD Microtainer blood collection tube (Ref#: 365967, Becton Dickinson) followed by centrifugation at 300g at 4°C for 5 min. The isolated serum sample was stored at −80 °C until analysis.

##### 2,4,6-trinitrobenzene sulfonic acid (TNBS)-induced colitis model

BALB/cJ female mice were fasted for 14-16 hrs prior to TNBS administration with free access to drinking water. 5% TNBS (Cat#: P2297, Sigma–Aldrich) was diluted to 30 mg/mL in absolute ethanol (40% v/v) and 100μl administered via enema by inserting a 3.5 French catheter 4 cm into the colon. 24mg/kg NOP or the same volume of H_2_O was orally gavaged daily starting at day 0 TNBS and continuing through day 5; oral gavage was done at approximately the same time each day (±2 hours) along with body weight measurement. Upon sacrifice, the colon was harvested and then fixed with 4% PFA for histologic assessment.

##### Anti-CD40-induced colitis model

C57BL/6 *Rag2^-/-^* mice were used at 12-13 weeks of age. At day 0, anti-CD40 colitis was induced by intraperitoneal injection (i.p.) of 200µg of anti-CD40 (Cat #:PE0016-2, IgG2 monoclonal Ab from BioXCell). Anti-CD40 mAb was diluted to 1 mg/mL in InVivoPure pH 7.0 dilution buffer (Cat #: IP0070, BioXCell). 24mg/kg NOP or the same volume of H_2_O was orally gavaged daily starting at day 0 anti-CD40 and continuing through day 7; oral gavage was done at approximately the same time each day (±2 hours) along with body weight measurement. Upon sacrifice, the colon was harvested and half of the most immediate portion of the proximal colon was collected and snap frozen, the rest of the colon tissue was fixed with 4% PFA for histologic assessment. The snap frozen proximal colon tissue was stored at −80 °C until analysis.

##### Histology

PFA-fixed colon tissues were stored in 70% ethanol prior to routine paraffin embedding, sectioning and hematoxylin and eosin (H&E) staining. Histologic colitis scores were determined by a pathologist (J.N.G.), who was blinded to the experimental parameters. Each of the four histologic parameters was scored as absent (0), mild (1), moderate (2), or severe (3): mononuclear cell infiltration, polymorphonuclear cell infiltration, epithelial hyperplasia, and epithelial injury. The scores for the parameters were summed to generate the histologic colitis score and were further quantified to include the percentage involvement by the disease process: (1) <10%; (2) 10–25%; (3) 30–50%; (4) >50% and presented as histologic colitis scores as follows: cumulative score * % involvement.^67^

##### Detection of NF-κB activation

Colons were excised, fat was removed, and intestines were opened longitudinally. After washing with PBS to remove residual debris, colonic tissue was incubated in PBS containing 5mM HEPES (ThermoFisher Scientific) and 1mM DTT for 10 min on ice. Tissues were then incubated in PBS with 5mM EDTA (Millipore Sigma) and 3% fetal bovine serum (FBS) (ThermoFisher Scientific) for two 15 min cycles at 37°C. The isolated mucosal cells were processed for nuclear protein isolation with NE-PER™ Nuclear and Cytoplasmic Extraction Reagents (Cat# 78833, ThermoFisher Scientific). Protein concentration of the nuclear fraction was determined using the BCA reagent (ThermoFisher Scientific), equal amounts of protein were separated by sodium dodecyl sulfate polyacrylamide gel electrophoresis (SDS-PAGE) (Biorad) and transferred to polyvinylidene fluoride membrane (PVDF, ThermoFisher Scientific). The membrane was incubated for 1h in 5% nonfat dry milk in TBS-T blocking buffer and all blots were probed with antibodies directed against NF-κB p65 subunit (Cat #: 8242S, Cell Signaling Technology) and Lamin B1(Cat #: 13435S, Cell Signaling Technology). Western blot band intensity was quantified using Image Lab software (Bio-Rad), and results were normalized to the corresponding loading control. Uncropped Western blot images corresponding to Figure 5h are provided in Supplemental Data 32.

##### Multiplex assay

Frozen proximal colon tissues from the anti-CD40-induced colitis model were thawed on ice and mechanically disrupted in PBS supplemented with protease inhibitors (Roche) and phosphatase inhibitors (Roche). Tissue homogenization was performed using a TissueRuptor (Qiagen), followed by centrifugation at 15,000 × g for 20 minutes at 4 °C to collect tissue supernatants. Levels of 13 proinflammatory cytokines and chemokines were measured using the LEGENDplex™ Mouse Inflammation Panel (BioLegend), following the manufacturer’s instructions. A volume of 25 μL of either tissue supernatant or serum was used per sample. LEGENDplex data were acquired on a BD LSR II flow cytometer (BD Biosciences) and analyzed using LEGENDplex Data Analysis Software (BioLegend).

##### Isolation of immune cells from colonic lamina propria

Colons were excised, fat was removed, and intestines were opened longitudinally. After washing with PBS to remove residual debris, colonic tissue was incubated in PBS containing 5mM HEPES (ThermoFisher Scientific) and 1mM DTT for 10 min on ice. Tissues were then incubated in PBS with 5mM EDTA (Millipore Sigma) and 3% fetal bovine serum (FBS) (ThermoFisher Scientific) for two 15 min cycles at 37°C. After vortexing for 15s, tissues were washed in PBS, cut into pieces < 1mm, and digested in RPMI 1640 (ThermoFisher Scientific) and 3% FBS, containing 0.5 mg/mL collagenase D (Roche), 0.03 mg/mL DNase I (Roche), 0.5mg/mL dispase (Stem Cell Technologies), and 1% penicillin/streptomycin (Corning). Digests were carried out in two rounds (25 min and 20 min) at 37°C on a rotating mixer, before vortexing 15s and passing through 70 um strainers into PBS with 5mM EDTA and 2% FBS. Cells were washed and resuspended in FACS buffer (PBS with 2% FBS and 1 mM EDTA). Cells were counted on the LUNA-FX7TM Automated Cell Counter and prepared for flow cytometry.

##### Flow cytometry

Single-cell suspensions were stained with fixable near-IR dead cell stain kit (ThermoFisher Scientific) in PBS containing CD16/32 antibody (93; BioLegend) to block non-specific binding to Fc receptors, according to manufacturer instructions. Next, cells were stained with antibodies against surface proteins (see list of reagents) in FACS buffer for 20 min on ice. Following surface staining, cells were washed twice with FACS buffer then fixed and permeabilized with eBioscience Foxp3/Transcription Factor Staining Buffer Set (ThermoFisher Scientific) according to manufacturer instructions. Intracellular proteins were then stained with antibodies in permeabilization buffer from the eBioscience Foxp3/Transcription Factor Staining Buffer Set for 1 hr at room temperature. Cells were washed twice, resuspended in FACS buffer, acquired on a BD FACSymphony A3 Cell Analyzer (BD Biosciences), and analyzed with FlowJo version 10.10 (BD Biosciences).

For cytokine detection, cells were cultured in RPMI 1640 with 10% FBS and eBioscience Cell Stimulation Cocktail (plus protein transport inhibitors) (ThermoFisher Scientific) for 4 h at 37°C then washed twice and fixed in BD Cytofix Fixation Buffer (BD Biosciences) for 30 min at room temperature, then fixed and permeabilized using the eBioscience Foxp3/Transcription Factor Staining Buffer Set, followed by intracellular staining as above.

##### *In vitro* activation of naïve CD4⁺ and CD8⁺ T cells

Naïve (CD44^lo^) CD8+ or naïve CD4+ T cells were activated as previously described with slight modifications.^39^ Briefly, spleens from C57BL/6J mice were pooled, dissociated into single-cell suspensions, and CD8⁺ T cells (STEMCELL Technologies) and CD4⁺ T cells (STEMCELL Technologies) were isolated by negative selection following manufacture protocols. The post-selection purity was consistently > 97%. Flat bottom 24-well tissue culture plates (Costar) were precoated with 10 μg/ml anti-CD3 (BioXCell) and 0.8 μg/ml B7–1/Fc Chimera (RnD Systems) for 3 hours at 37°C. 0.5 X 106 CD44low T cells in 2 ml of AIM-V serum-free medium (Thermo Fisher Scientific) supplemented with 1 mM HEPES, 1 mM sodium pyruvate, non-essential amino acids (Gibco), and 0.025 mM 2-mercaptoethanol (Thermo Fisher Scientific). Cells were cultured in the presence of 100 U/ml recombinant IL-2 (NIH), recombinant IL-12 (2.5 ng/ml for CD8+ and 10ng/ml for CD4+ T cells, R&D Systems), and NOP at 0, 1 and 2.5µM.

At 67 h, activated T cells were treated with eBioscience Protein Transport Inhibitor Cocktail (ThermoFisher Scientific) for 5 h at 37°C. Activated CD8+ and CD4+ T cells were harvested at 72 h and stained with antibodies against surface proteins (see list of reagents) in FACS buffer for 30 min at 4°C. CD16/32 antibody (BioLegend) was used to block non-specific binding to Fc receptor and cell viability was determined using Ghost Dye Red 780 (Tonbo Biosciences). Following surface staining, cells were washed twice with FACS buffer then fixed and permeabilized with Tonbo Biosciences Foxp3/Transcription Factor Staining Buffer Kit (CYTEK) according to manufacturer instructions for 30 min on ice. Then, cells were washed with Perm/Wash buffer twice followed by intracellular staining (see list of reagents) overnight at 4°C. Cells were washed twice, resuspended in FACS buffer. Data were acquired on a BD FACSymphony A3 Cell Analyzer (BD Biosciences) and analyzed with FlowJo version 10.10 (BD Biosciences).

### QUANTIFICATION AND STATISTICAL ANALYSIS

Data were analyzed with GraphPad Prism (version 10). Data are shown as mean with individual data points as noted. For comparison between two independent experimental groups, a two-tailed Mann-Whitney U test was used. For comparison between more than two groups, one-way ANOVA followed by Dunnett’s test was performed. Details of statistical analysis and sample size are provided in the figure legends and STAR methods. No samples were excluded from any experiments performed in this study unless in the case of technical failure. Experimenters were not blinded to experimental conditions. Differences of P < 0.05 were considered statistically significant.

## Supplemental information

### Supplemental Figure Legends

**Figure S1.**
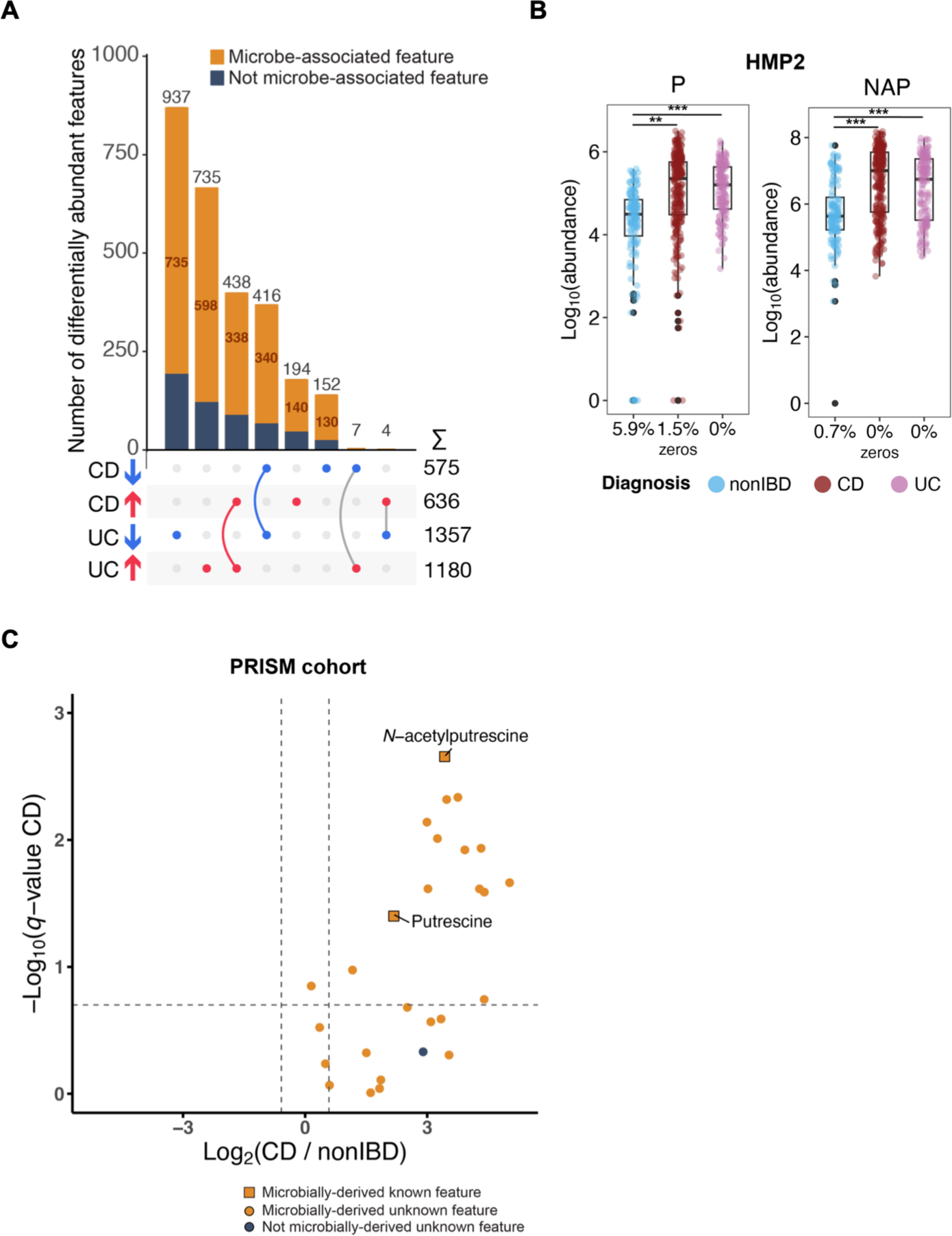
Differentially abundant metabolic features in HMP2 and independent validation of the putrescine module in the PRISM CD cohort. (A) UpSet plot showing the DA metabolic features (linear mixed-effects model, abs (log2 fold change) > log2(1.5), false discovery rate (FDR)-adjusted *P*<0.2; Method) in CD and UC after gnotobiotic microbiome filtering. The number of enriched and depleted metabolic features in CD and UC are displayed in the right bottom corner. Non-microbe-associated (navy blue) and microbe-associated features (orange). (B) Box plots of the relative abundances of two differentially abundant anchor compounds, putrescines (P) and N-acetylputrescine (NAP), in HMP2 fecal samples (non-IBD n=135, CD n=265, UC n=146). The percentage of zeros is shown on the x-axis. Statistical analysis was performed using a linear mixed model. \**q*-value (FDR-adjusted p-value) < 0.2, **q< 0.05, ***q<0.01. 1^st^, 2^nd^ and 3^rd^ quartiles of the data. Boxplot whiskers indicate the inner fences of the data with points outside the inner fences plotted as outliers. (C) Volcano plot depicting metabolic features (n=27) in PRISM CD fecal metabolomes that were aligned with metabolic features in module #122 in the HMP2 data and mouse fecal metabolomes. Twelve microbially-associated unknown features were consistently enriched in HMP2 CD, UC and PRISM CD (linear model, abs fold change > 1.5, FDR-adjusted p-value<0.2; Method).

**Figure S2.**
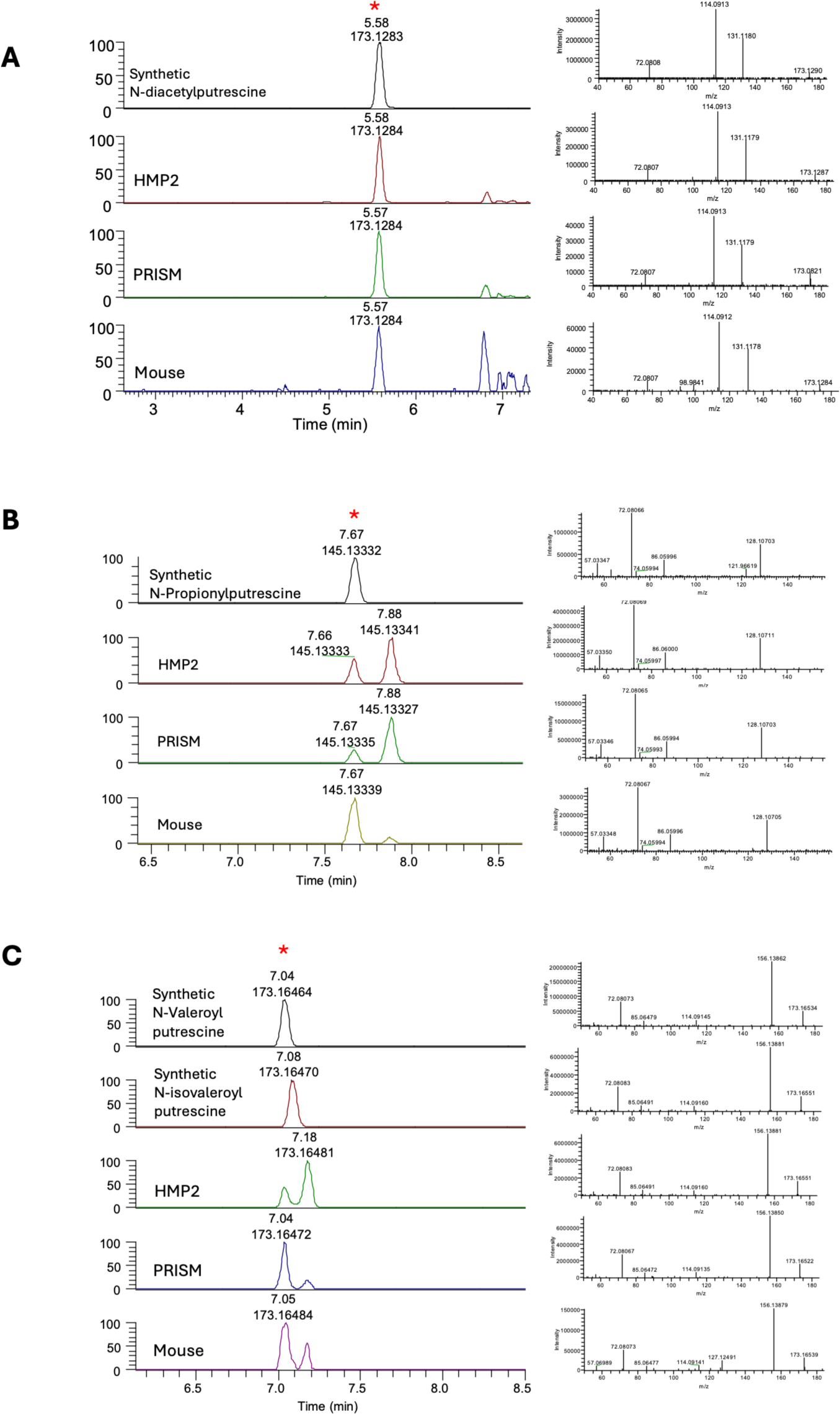
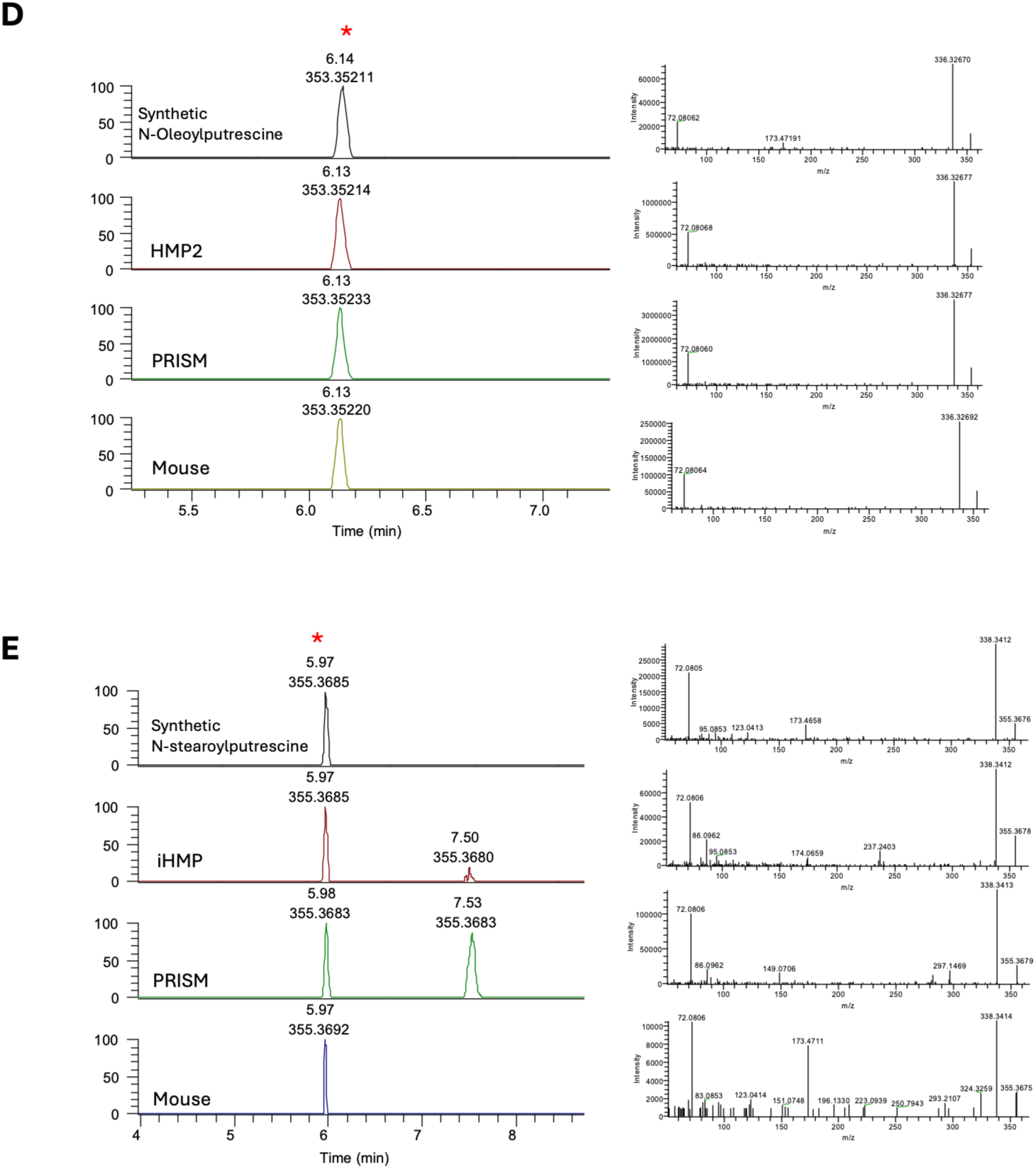
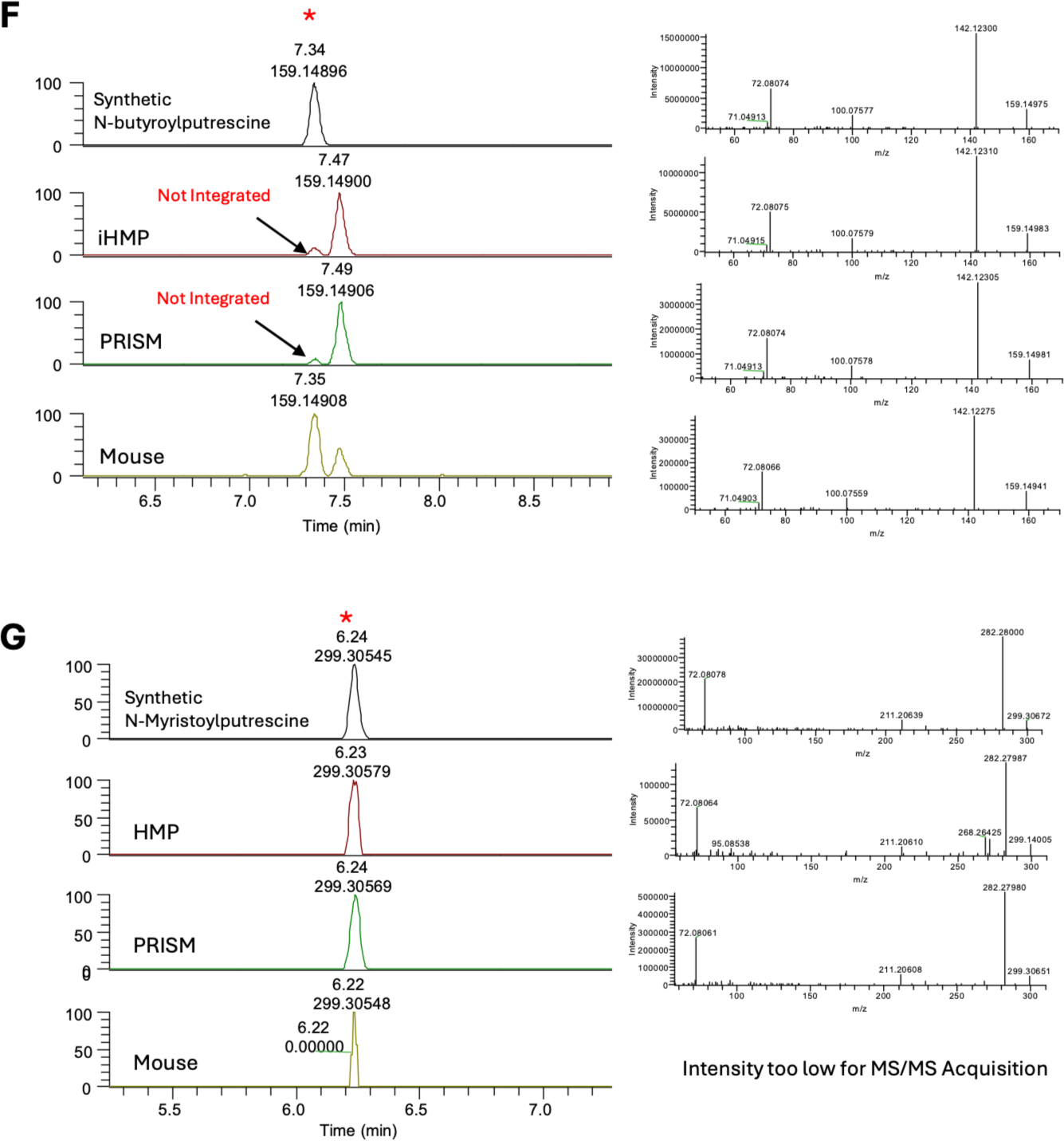
Validation of acylated putrescines in HMP2 and PRISM (human IBD cohorts) and mouse metabolomics data by retention time (RT) and MS/MS matching to synthetic standard. (A-G) Acylated putrescine standards. (A) Diacetylputrescine, (B) *N*-propionoylputrescine, (C) *N*-valeroylputrescine and *N*-isovaleroylputrescine, (D) *N*-oleoylputrescine, (E) *N*-stearoylputrescine, (F) *N*-butyroylputrescine and (G) *N*-myristoylputrescine detected in HMP2, PRISM and mouse stool samples using a Thermo QE Orbitrap in the positive mode.

**Figure S3.**
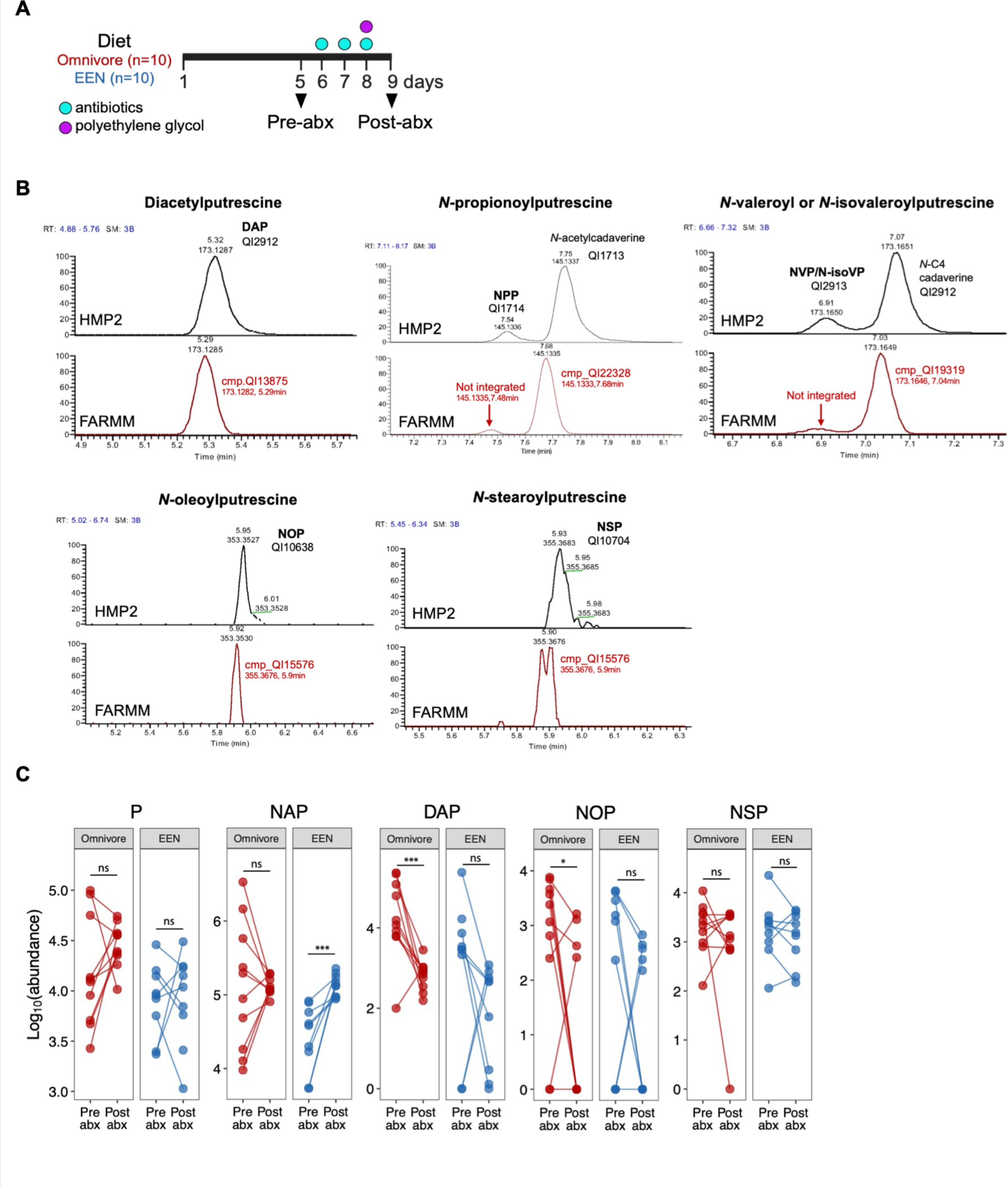
Human gut microbiota depletion alters fecal acylated putrescine abundance. (A) Partial overview of FARMM study design. To assess microbial influences on the intestinal metabolome, pre-antibiotic (day 5) and post-antibiotic treated (day 9) samples were used for human microbiome filtering analysis. Healthy inpatient subjects received defined diets throughout the study (omnivore n=10; EEN n=10 subjects). (B) Detection of acylated putrescines in the FARMM fecal metabolome. Diacetylputrescine (DAP), *N*-oleoylputrescine (NOP), and *N*-stearoylputrescine (NSP) were detected, whereas *N*-propionoylputrescine and *N*-valeroyl/*N*-isovaleroylputrescine were not detected in FARMM fecal samples. Metabolic feature IDs corresponding to each peak are indicated on the plots. (C) Fecal abundances of two known metabolites putrescine (P) and *N*-acetylputrescine (NAP) and acylated putrescine pre- and post-antibiotic and polyethylene glycol treatment detected in FARMM study. Circles represent individual subjects. Samples collected from the same subjects across microbiota depletion intervention are connected by lines. Statistical analysis was performed using a linear mixed model (Method). *q-value (FDR-adjusted *p*-value) < 0.2, **q< 0.05, ***q<0.01; ns, not significant.

**Figure S4.**
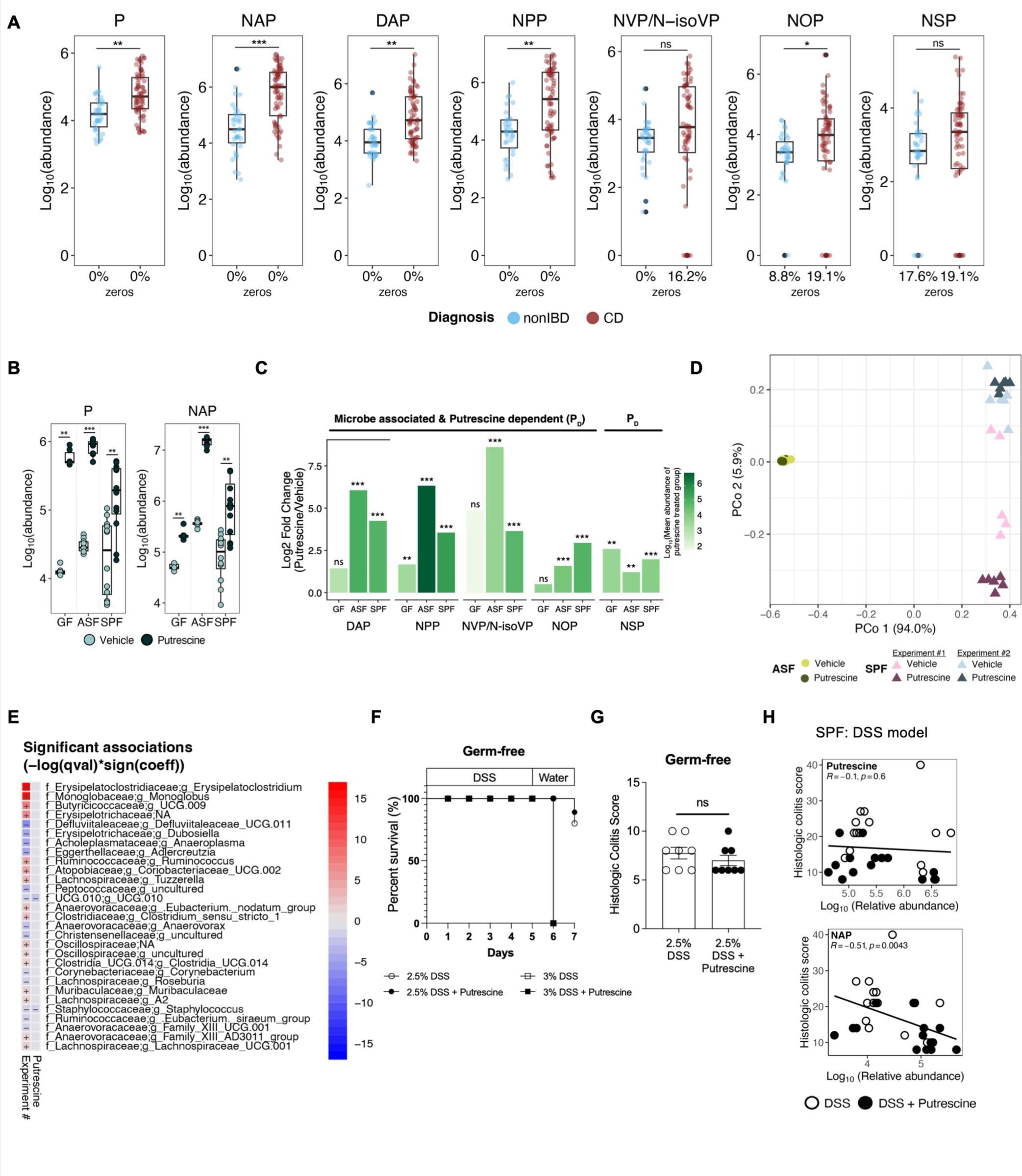
The gut microbiota utilizes putrescine and produces acylated putrescines that have the potential to mitigate intestinal inflammation and injury. (A) Box plots of the relative abundances of putrescine, *N*-acetylputrescine, and the 5 acylated putrescines in the PRISM fecal samples (non-IBD n=34, CD n=68). Circles represent metabolic feature abundance from individual sample. The percentage of zeros is shown on the x-axis. Statistical analysis was performed using a linear model. *q-value (FDR-adjusted p-value) < 0.2, **q< 0.05, ***q<0.01. (B) Box plots of the relative abundances of P and NAP from GF, ASF and SPF C57BL/6 mice with and without putrescine supplementation. Circles represent data from individual mice. *p<0.05, **p<0.01, ***p<0.001, ns (not significant), Wilcoxon test. (C) Bar plots showing the log2 fold change in relative abundance of DAP, NPP, NVP/N-isoVP, NOP and NSP in putrescine-treated mice compared to vehicle-treated mice with the same microbiome background, including germ-free (GF, control n=6; putrescine n=6), altered Schaedler flora (ASF, control n=10; putrescine n=9) and specific pathogen free (SPF, control n=12; putrescine n=12) C57BL/6 mice. *p<0.05, **p<0.01, ***p<0.001, ns (not significant), Wilcoxon test. Bar color indicates log10 transformed mean abundance of acylated putrescines in each putrescine-treated group. (D) Ordination plot using Principal Coordinates Analysis (PCoA) based on Bray-Curtis dissimilarity distances between samples from bacterial 16S rRNA gene amplicon V4 sequences of ASF (circle shape, vehicle n=6, putrescine n=9) and SPF (triangle shape, vehicle n=11, putrescine n=12). Separate colors were used for samples from different experiments for the SPF mice to highlight Jackson vivarium room differences. (E) MaAsLin2 analysis of differentially abundant amplicon sequence variants (ASV) in SPF mice with or without 1% w/v putrescine in the drinking water. (F) Survival curves of germ-free C57BL/6 mice treated with 2.5% or 3% w/v DSS in the drinking water (3% w/v DSS treatment ctrl n=5, putrescine n=5; 2.5% w/v DSS treatment ctrl n= 10, putrescine n= 9). (G) Histologic colitis scores of putrescine-supplemented (n=8) and control (n=8) germ-free C57BL/6 mice treated with 2.5% w/v DSS treatment. Data reflect two independent experiments. Circles represent data from individual mice and mean ± SEM are shown, ns not significant, two-tailed Mann-Whitney U test. (H) Spearman’s correlation coefficient and *P* values for the relative abundance of P and NAP and histologic colitis score respectively. Control mice (n=14), white circles, and putrescine supplemented mice (n=15), black circles.

**Figure S5.**
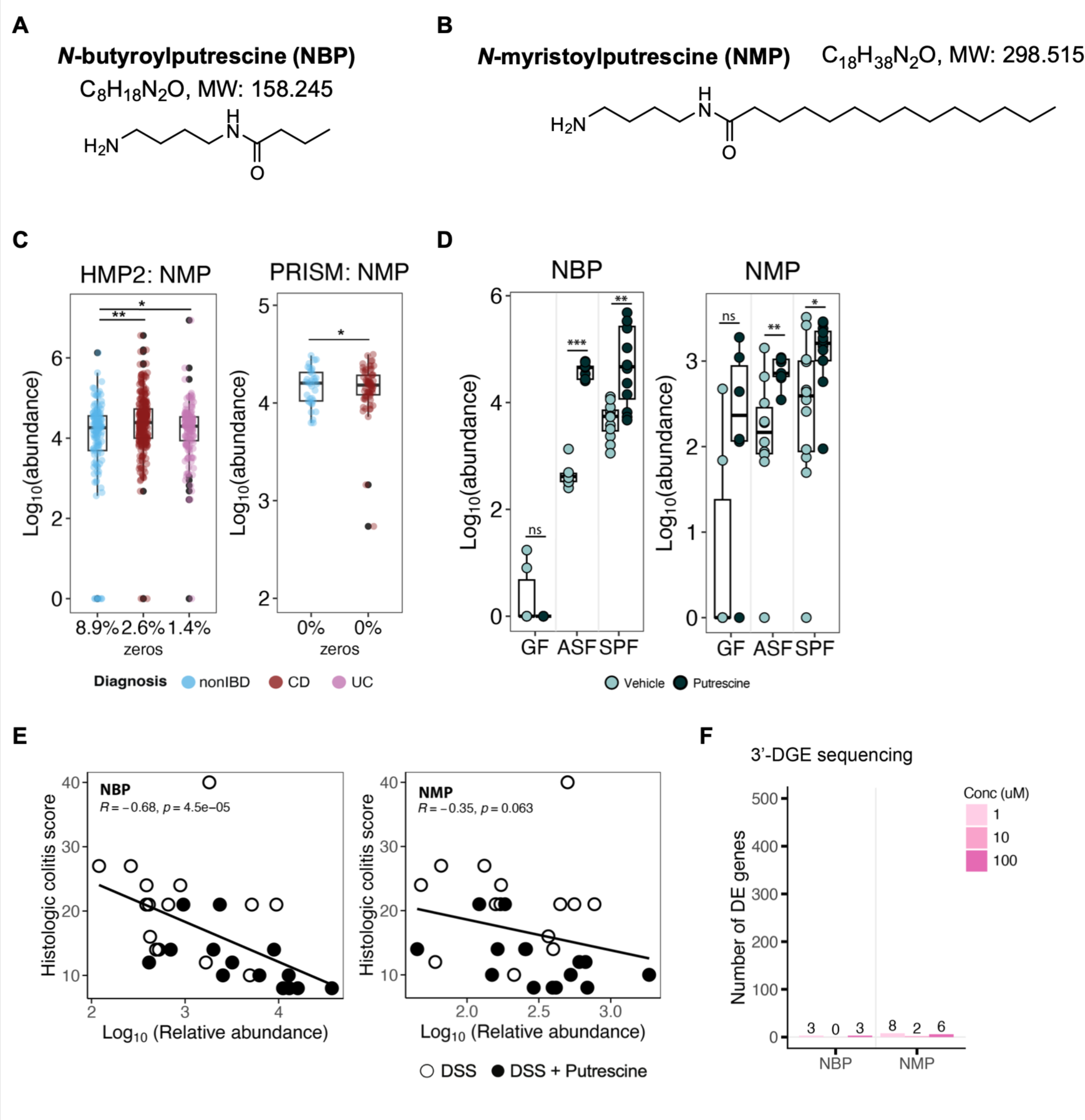
Other acylated putrescines not in module #122. (A-B) Chemical structures, formulas, and molecular weights of two acylated putrescines used in our studies, *N*-butyroylputrescine (NBP) (A) *N*-myristoylputrescine (NMP) (B) neither of which are in module #122. (C) Box plots of the relative abundances of NMP in HMP2 fecal samples (non-IBD n=135, CD n=265, UC n=146) and PRISM fecal samples (non-IBD n=34, CD n=68). The percentage of zeros is shown on the x-axis. Statistical analysis was performed using a linear mixed model for the HMP2 dataset and using a linear model for the PRISM dataset. *q-value (FDR-adjusted p-value) < 0.2, **q< 0.05, ***q<0.01. (D) Box plot of the relative abundances of NBP and NMP from germ-free (GF, control n=6; putrescine n=6), altered Schaedler flora (ASF, control n=10; putrescine n=9) and specific pathogen free (SPF, control n=12; putrescine n=12) C57BL/6 mice with or without putrescine supplementation. *p< 0.05, **p<0.01, ***p<0.001, ns not significant, Wilcoxon test. (E) Spearman’s correlation coefficient and *P* values for the relative abundance of NBP and NMP from 3% DSS-treated mice and histologic colitis score. (F) Number of differentially expressed genes in mouse bone marrow-derived dendritic cells (BMDCs) incubated with NBP or NMP (1, 10, 100 μM) that were profiled using 3’-Digital Gene Expression (3’-DGE) Sequencing. Control mice (n=14), white circles, and putrescine supplemented mice (n=15), black circles. Boxplot boxes (C, D) indicate the 1^st^, 2^nd^ and 3^rd^ quartiles of the data. Boxplot whiskers (C) indicate the inner fences of the data, with points outside the inner fences plotted as outliers.

**Figure S6.**
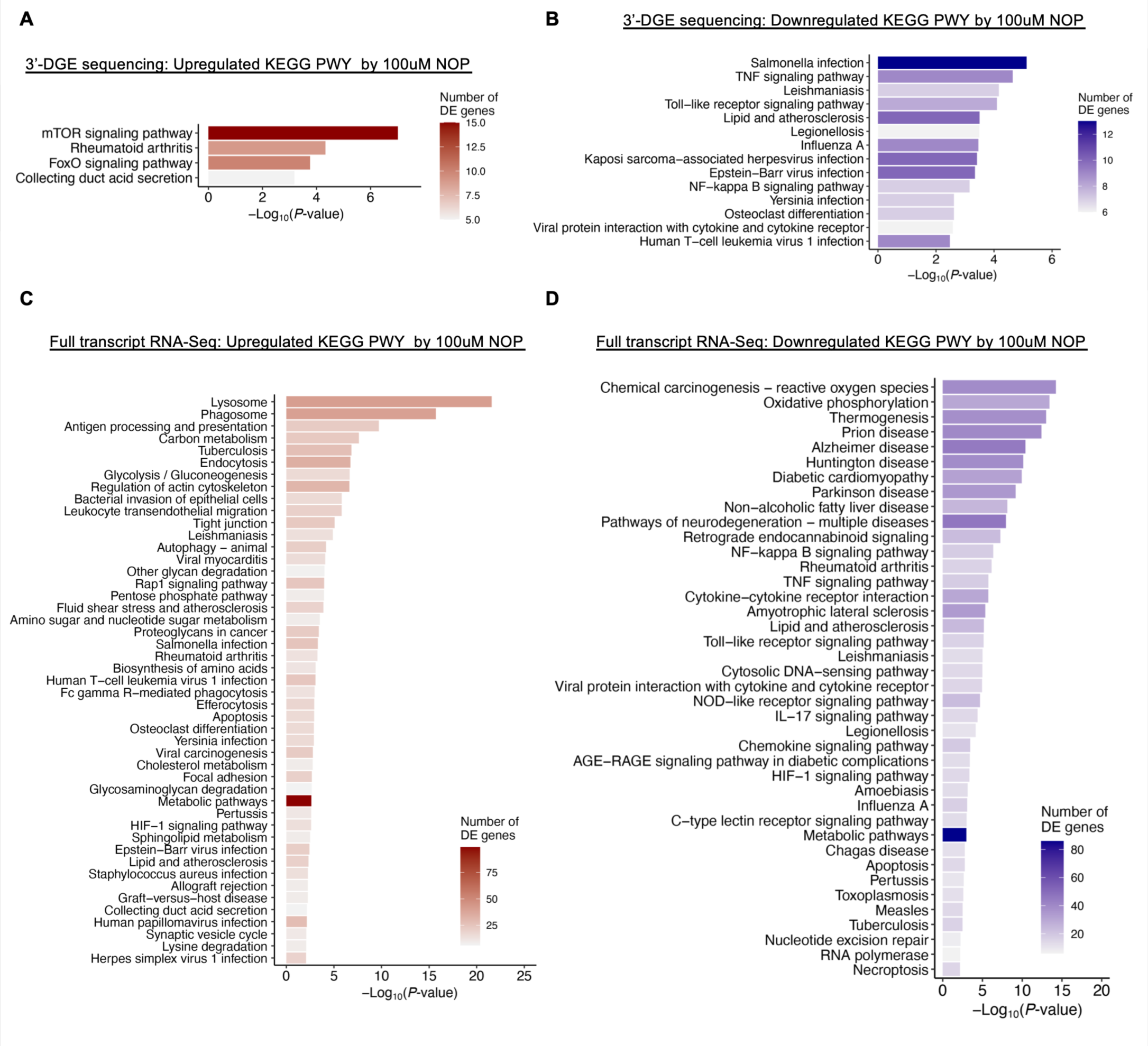
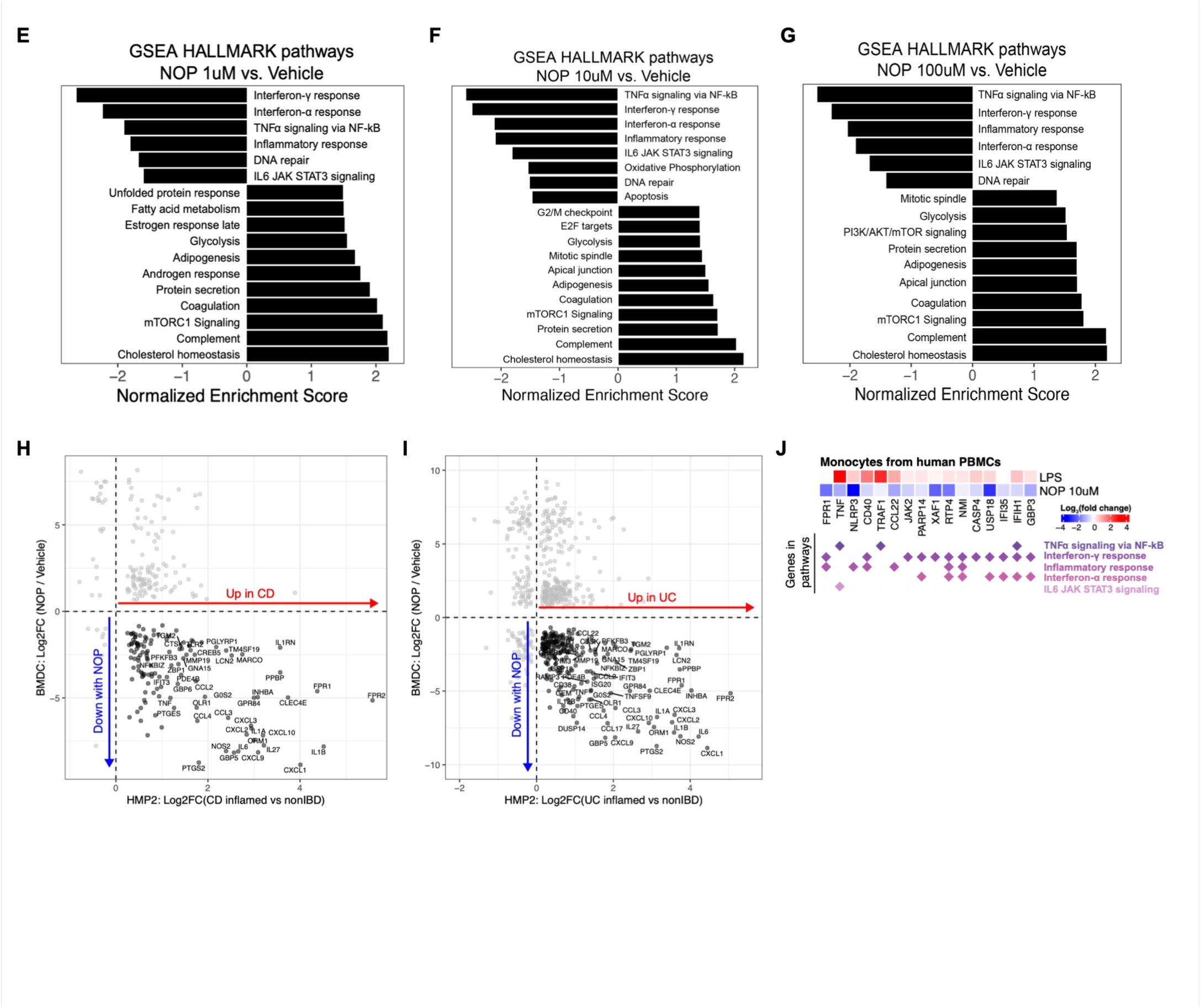
NOP treatment suppressed pro-inflammatory gene expression. (A-D) DAVID functional Kyoto Encyclopedia of Genes and Genomes (KEGG) analysis of differentially expressed genes (FDR-adjusted p-value < 0.05 & abs (log2 fold change) > log2(1.5)) by 100 μM NOP-treated mouse BMDC. (A) Upregulated KEGG pathways profiled by 3’-DGE Sequencing. (B) Downregulated KEGG pathways profiled by 3’-DGE Sequencing. (C) Upregulated KEGG pathways profiled by full transcript RNA sequencing. (D) Downregulated KEGG pathways profiled by full transcript RNA sequencing. (E-G) Hallmark Gene Set Enrichment Analysis (GSEA) of BMDC gene expression comparing NOP treatment at 1 μM (E), 10 μM (F) and 100 μM (G) NOP with vehicle control. Key inflammatory pathways that are implicated in IBD were significantly downregulated across all NOP concentrations tested. (H-I) Log2 fold change comparison of transcript expression between NOP-treated BMDC and HMP2 biopsy tissue RNA-seq data. The fold change comparison between 100 μM NOP-treated BMDC and CD inflamed vs. non-IBD (H), UC inflamed vs. non-IBD (I) are shown respectively. Differentially expressed genes (Wald’s test, FDR-adjusted *P* < 0.05) in both the BMDC and HMP2 dataset are displayed. Genes that were upregulated in HMP2 IBD inflamed tissue samples and were down regulated by NOP treated BMDC are indicated by black dots. (J) Heatmap showing gene expression patterns of human PBMC-derived monocytes incubated with NOP or LPS that are upregulated in inflamed CD or UC tissues (Wald’s test, FDR-adjusted *P* < 0.05) from the HMP2 cohort but downregulated following NOP treatment in mouse BMDCs (see main Figure 3) (Wald’s test, FDR-adjusted *P* < 0.05). Genes associated with specific inflammatory pathways are indicated by diamond symbols.

**Figure S7.**
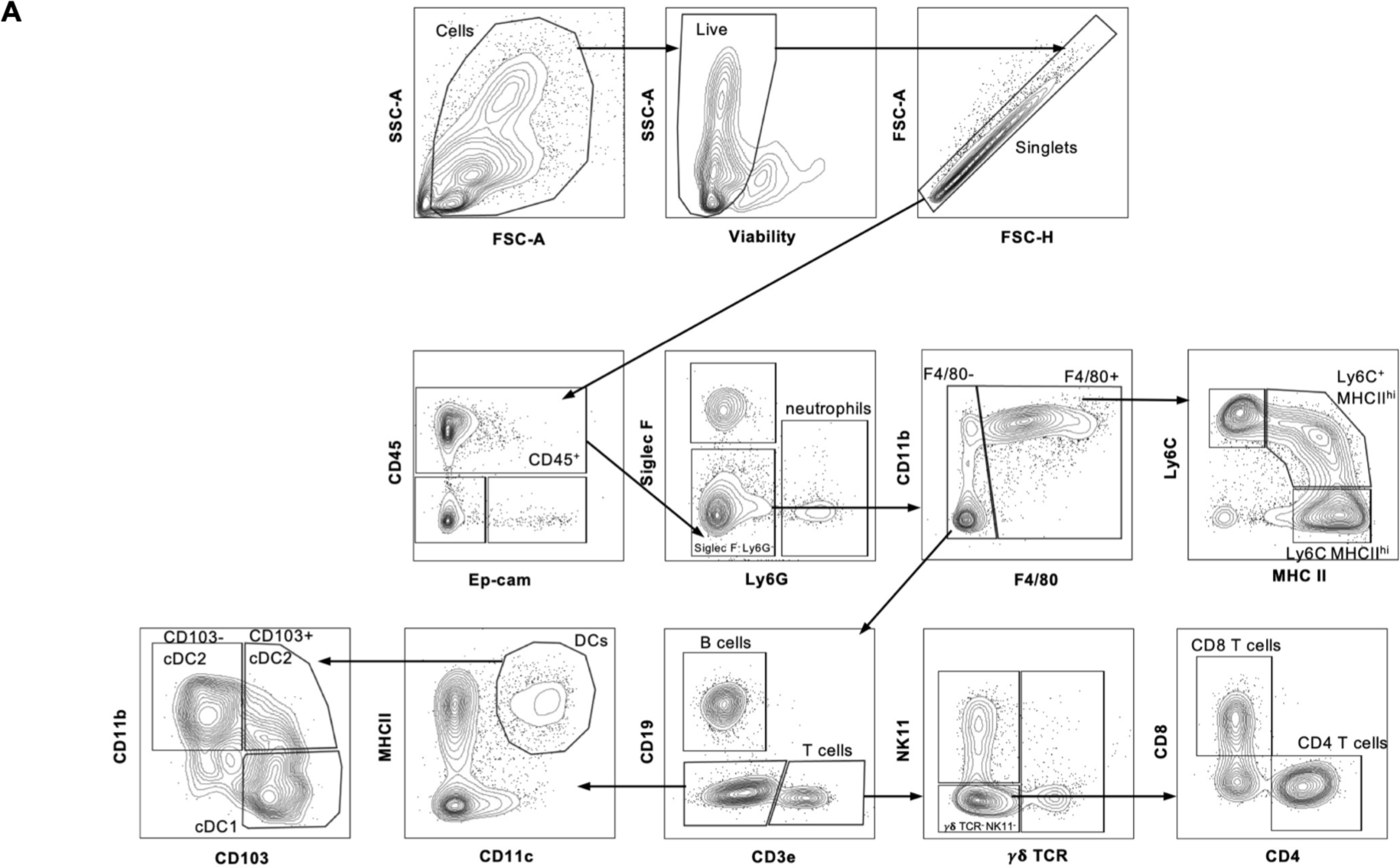
Gating strategy for flow cytometry.

**Figure S8.**
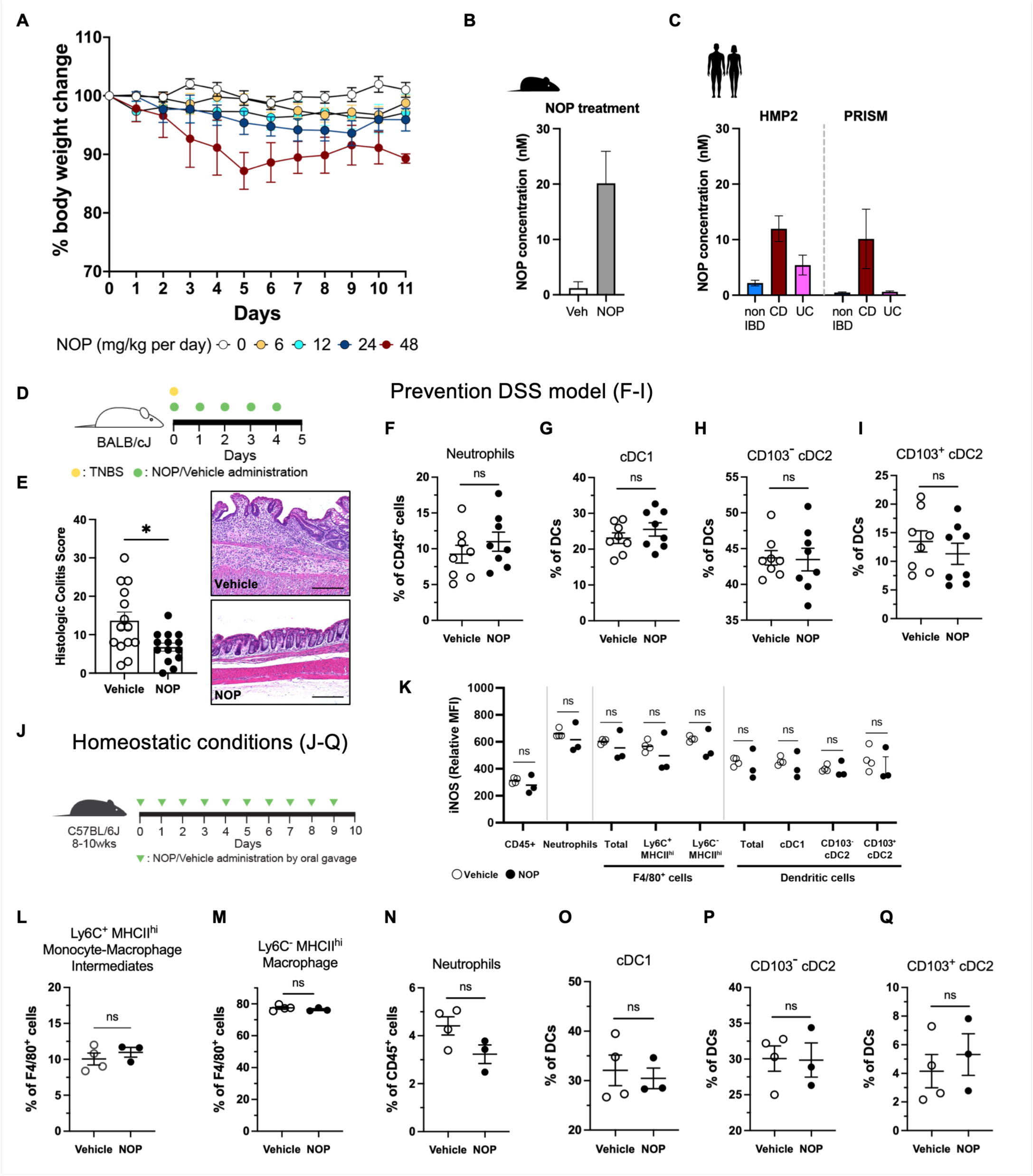
Dose-response evaluation of NOP *in vivo* and its immunomodulatory effects under intestinal inflammatory and homeostatic conditions. (A) Body weight changes of C57BL/6 mice treated with daily NOP oral gavage (0mg/kg n=5; 6mg/kg n=4; 12mg/kg n=4; 24mg/kg n=5; 48mg/kg n=5). Circles represent mean ± SEM. (B-C) Average estimated NOP concentrations (nM) measured in fecal samples from mice and two human IBD cohorts (HMP2 and PRISM). (B) Healthy SPF C57BL/6J mice were orally gavaged with NOP (24mg/kg/day; n=3) or vehicle (n=4) for 10 consecutive days. Single experiment. (C) HMP2 fecal samples (non-IBD n=135; CD n=265; UC n=146) and PRISM fecal samples (non-IBD n=34; CD n=68; UC n=53). Mean ± SEM are shown. NOP concentration in stool samples was calculated using an external calibration curve generated with synthetic NOP standard. (D) Schematic of 2,4,6-trinitrobenzene sulfonic acid (TNBS)-induced intestinal injury model. (E) Histologic colitis scores of vehicle-treated (n=14) or NOP-treated (n=14) mice with representative photomicrographs. Scale bars, 200µm; 3 independent experiments. (F-I) Flow cytometry (FC) analysis of the prevention DSS model (vehicle n=8; NOP n=8, 2 independent experiments). (F) FC analysis of neutrophils, cDC1 (G), CD103^-^ cDC2 (H), CD103^+^ cDC2 (I). (J-Q) FC analysis of the effect of NOP under homeostatic conditions (vehicle n=4; NOP n=3; single experiment). (J) Experimental scheme. (K) Intracellular iNOS expression in colonic lamina propria CD45+ cells, neutrophils, F4/80+ cells and dendritic cells. (L) Ly6C^+^MHCII^high^ monocyte-macrophage intermediates. (M) Ly6C^-^MHCII^high^ macrophages. (N) neutrophils. (O) cDC1. (P) CD103^-^ cDC2. (Q) CD103^+^ cDC2. Circles represent data from individual mice, mean ± SEM are shown ns (not significant), *p<0.05. Two-tailed Mann-Whitney U test (E-I and K-Q).

**Figure S9.**
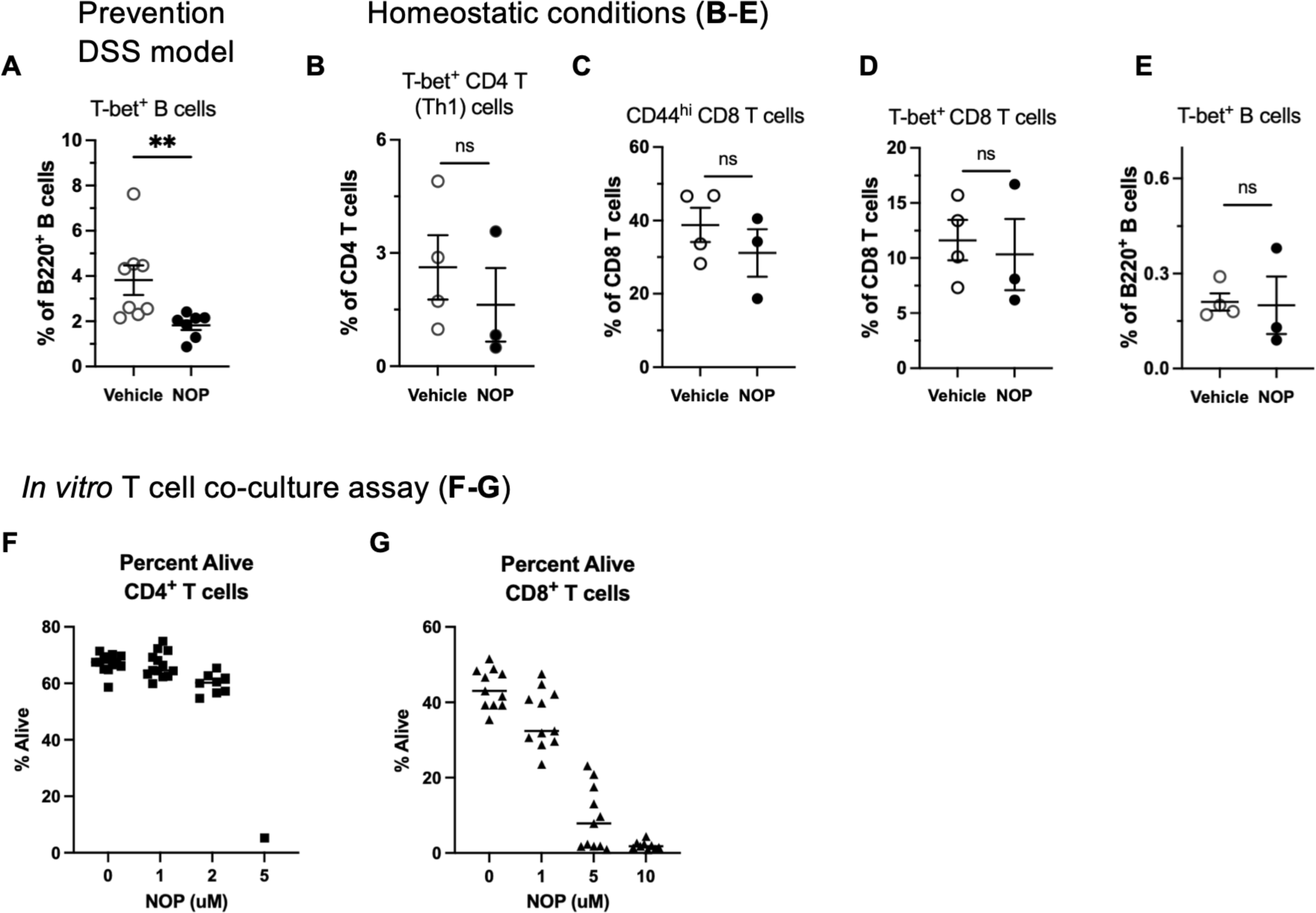
Adaptive lymphocyte immunomodulatory effects of NOP under intestinal inflammatory and homeostatic conditions. (A) Flow cytometry (FC) analysis of T-bet^+^ B cells in the prevention DSS model (vehicle n=8; NOP n=8, 2 independent experiments). (B-E) FC analysis of Th1 cells (B), CD44^hi^ CD8 T cells (C), T-bet^+^ CD8 T cells (D), and T-bet^+^ B cells (E) following NOP treatment under homeostatic conditions (vehicle n=4; NOP n=3; single experiment). Circles represent data from individual mice, mean ± SEM are shown *p<0.05, **p<0.01, ns (not significant). Two-tailed Mann-Whitney U test (A-E). (F-G) Viability of CD4⁺ (F) and CD8⁺ (G) T cells cultured in vitro for 3 days with anti-CD3, B7-1, IL-2, and IL-12, in the presence or absence of NOP.

**Figure S10.**
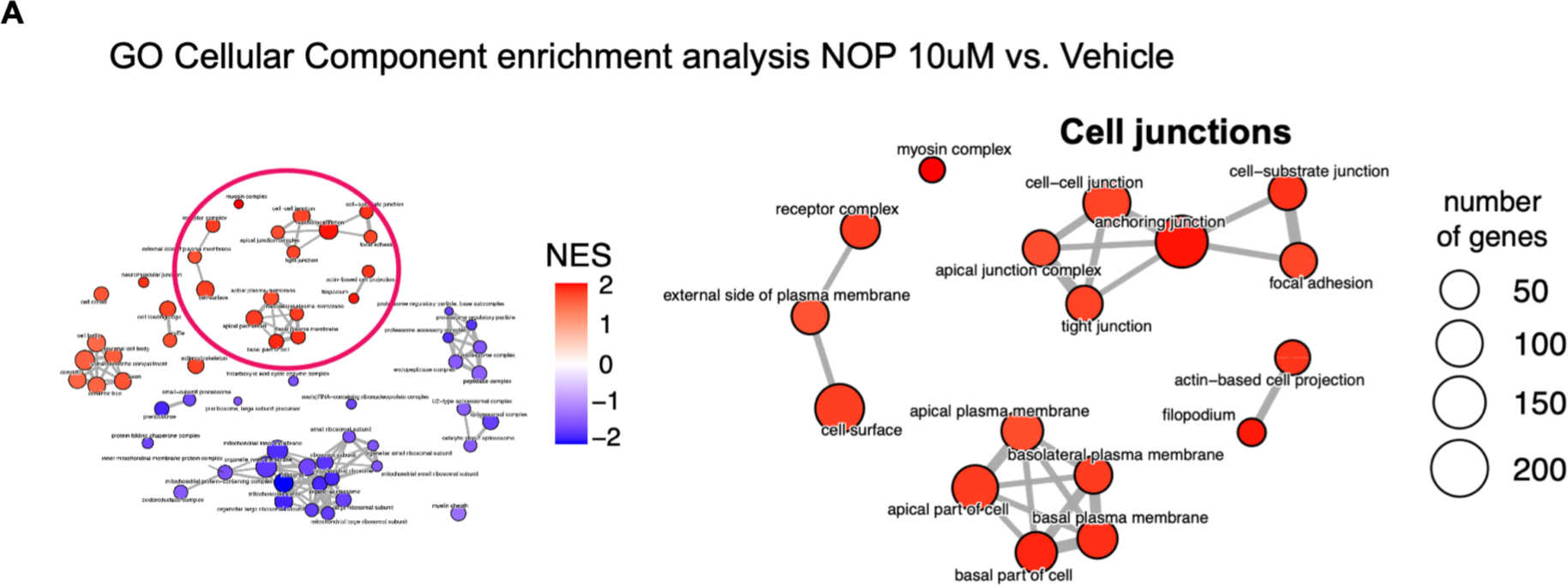
NOP transcriptional effects on intestinal barrier integrity in mouse colonic organoids. (A) Pathway enrichment network analysis. Gene Ontology (GO) Cellular component enrichment analysis of full-transcript organoid RNA-Seq comparing 10 μM NOP treatment to vehicle control. Pathways related to intestinal barrier integrity, including cell junctions, tight junction, and cell surface enriched upon NOP treatment; these enriched pathways are indicated by red circle with zoomed-in inset. Each circle represents a pathway, and circle size corresponds to the number of genes in that pathway. Normalized Enrichment Score (NES) is represented by color.

### Supplemental Tables

**Supplemental Table 1: Metabolic features in prioritized HMP2 CD modules.** Metabolic features belong to modules that contain at least one differentially abundant (DA, abs fold change > 1.5, false discovery rate (FDR)-adjusted *p*-value<0.2) known metabolite, anchor compound, and IBD- and microbe-associated unknown feature in HMP2 CD patients. Disease association (CD vs. non-IBD, non-IBD as a reference) was tested with a linear mixed effects model (Methods). Features are sorted by module number. For each feature, coefficient estimates, the associated two-tailed p-values, FDR-adjusted p-values (q-values), log2 fold change values, and module identification number are provided. For the features that were aligned with mouse stool metabolic features, mouse feature id, mass-to-charge (m/z) ratio, retention time (RT) and microbial association annotation are provided.

**Supplemental Table 2: Metabolic features in prioritized HMP2 UC modules.** Table fields follow the same format as Supplemental Table 1. Disease association (UC vs. non-IBD, non-IBD as a reference) was tested with a linear mixed effects model (Methods).

**Supplemental Table 3**: Metabolic features shared across gnotobiotic-HMP2, and FARMM-HMP2 datasets acquired in HILIC-positive mode.

**Supplemental Table 4: IBD-associated and microbially-associated unknown features in module #122.** List of IBD-associated unknown features (linear mixed-effects model, abs fold change > 1.5, false discovery rate (FDR)-adjusted *p*-value<0.2; Method) in module #122 containing putrescine and *N*-acetylputrescine as anchor compounds. Table fields follow the same format as Supplemental Table 1 and 2.

**Supplemental Table 5: Metabolic feature alignments between HMP2 module #122, mouse stool metabolomes and PRISM CD metabolomes.** Feature ID, m/z and RT values of metabolic features aligned between HMP2 module #122, mouse stool metabolomics and PRISM CD metabolomes are provided. Disease association (CD vs. non-IBD) from PRISM dataset was tested with a linear model (Method). For each aligned feature, coefficient estimates, the associated two-tailed p-values and FDR-adjusted *p*-values (q-values), log2 fold change values are provided.

**Supplemental Table 6: Summary table of acylated putrescines in HMP2, mouse metabolome, and PRISM.**

**Supplemental Table 7: 16S rRNA V4 amplicon relative abundance analysis of the gut microbiota in ASF and SPF mice with and without putrescine supplementation.** Sample metadata include microbiome type, putrescine treatment (Y/N), cage, and experiment number.

**Supplemental Table 8: Relative abundance levels of putrescine, *N*-acetylputrescine and acylated putrescines in gnotobiotic mouse (GF, ASF, and SPF) fecal samples with and without putrescine supplementation.** The provided relative abundance values were median-normalized. Sample metadata include microbiome type and putrescine treatment (Y/N).

**Supplemental Table 9: Relative abundance levels of putrescine, *N*-acetylputrescine and acylated putrescines in DSS-treated SPF mouse fecal samples with and without putrescine supplementation.** The provided relative abundance values were median-normalized. Metadata for each sample, including information on putrescine treatment and associated colitis scores, are also provided.

**Supplemental Table 10: Differentially expressed genes from 100 μM NOP treatment in BMDCs profiled by 3’ Digital Gene Expression Sequencing.** List of differentially expressed genes (FDR-adjusted p-value < 0.05, abs fold change > 1.5) from 100 μM NOP treatment compared with vehicle treatment in BDMC. For each differentially expressed gene, log2 fold change values, the Wald test p-value and FDR-adjusted p-values are provided.

**Supplemental Table 11: DAVID functional Kyoto Encyclopedia of Genes and Genomes (KEGG) analysis of 100 μM NOP-treated BMDCs profiled by 3’-DGE Sequencing.** DAVID KEGG pathway analysis of the differentially expressed (DE) genes (FDR-adjusted p-value < 0.05, abs fold change > 1.5) from 100 μM NOP-treated BMDCs compared with vehicle treatment. Significantly overrepresented pathways (FDR-adjusted p-value < 0.05) and DE gene names belonging to those pathways are provided.

**Supplemental Table 12: Enrichment analysis of differentially expressed Hallmark pathways from NOP-treated BMDC co-cultures.** Significant differentially expressed pathways (FDR-adjusted p-value < 0.05) with NOP treatment compared with vehicle treatment and the associated p-value, FDR-adjusted *p*-values and normalized enrichment score (NES) from gene set enrichment analysis are provided. The comparisons were between 1 μM NOP vs. vehicle, 10 μM NOP vs. vehicle and 100 μM NOP vs. vehicle treatment in BMDCs.

**Supplemental Table 13: Enrichment analysis of differentially expressed Hallmark pathways in HMP2 host intestinal biopsies.** Significant differentially expressed pathways (FDR-adjusted p-value < 0.05), p-value, FDR-adjusted *p*-values and NES values from gene set enrichment analysis for comparisons between inflamed IBD and non-IBD intestinal biopsies. The comparisons were between biopsies from CD and non-IBD subject and UC and non-IBD subject.

**Supplemental Table 14: Marker genes for dendritic cell and inflammatory monocyte identification.**

**Supplemental Table 15: Per-sample metabolite relative abundance profiles of gnotobiotic fecal samples acquired in HILIC-pos mode.** Untargeted fecal metabolomes of GF, ASF and SPF mice that were maintained in the same environment. Sample metadata include microbiome type, cage, and sex.

**Supplemental Table 16: Per-sample metabolite relative abundance profiles of gnotobiotic fecal samples acquired in HILIC-neg, C18-neg and C8-pos modes.** Untargeted fecal metabolomes of GF, ASF and SPF mice that were maintained in the same environment. Sample metadata include microbiome type, cage, and sex.

### Supplemental Data

General experimental procedures for NPP, NBP, NVP, NMP, NOP, and NSP synthesis Experimental procedures for NOP hydrochloride synthesis

**S1-S30**: ^1^H and ^13^C NMR and HRESIMS spectra data of intermediates during synthesis and NPP, NBP, NVP, NMP, NOP, and NSP synthetic standards

**S31**: ^1^H NMR spectra data of NOP hydrochloride synthetic standard

**S32**: Uncropped Western blot images corresponding to Data 5h

#### General experimental procedures for NPP, NBP, NVP, NMP, NOP and NSP synthesis

Compound 1: *N*-propionoylputrescine
Compound 2: *N*-butyroylputrescine
Compound 3: *N*-valeroylputrescine
Compound 4: *N*-myristoylputrescine
Compound 5: *N*-oleoylputrescine
Compound 6: *N*-stearoylputrescine

##### Reagents, instruments, and techniques

High resolution electrospray ionization mass spectrometry (HRESIMS) was carried out using an Agilent 6530 LC-q-TOF mass spectrometer (Agilent Technologies, Santa Clara, CA) equipped with an Agilent 1290 UHPLC system. NMR spectra were obtained using Bruker Advance (^1^H: 400 MHz, ^13^C: 100 MHz) or Bruker NEO NMR system (^1^H: 600 MHz, ^13^C: 150 MHz) (Bruker, Billerica, MA) with CD_3_OD or CDCl_3_ (Cambridge Isotope Laboratories, Inc., Tewksbury, MA). HPLC for purification was carried out on an Agilent 1200 system equipped with an ELSD using a Phenomenex Luna C_8_ (2) column (5 μm, 250 × 10 mm). Thin layer chromatography (TLC) was performed using silica gel 60 F_254_ aluminum, together with the detection of ultraviolet and permanganate staining. All chemicals were HPLC grade.

###### General Procedure A

To a solution of *tert*-Butyl (4-aminobutyl)carbamate (50 mg, 0.27 mM) in CH_2_Cl_2_ (5 mL) was added DMAP (1.0 equiv), EDC HCl (2.7 equiv), and fatty acid (2.3 equiv). After the reaction was carried out for 24 hr at room temperature, the mixture was diluted with CH_2_Cl_2_ and washed with 0.5 N HCl, 1% K_2_CO_3_, and brine. The organic layer was dried over Na_2_SO_4_, filtered, and evaporated in vacuo. The crude material was purified by reversed-phase HPLC (Phenomenex, Luna C8 (2), 5 μm, 250 × 10 mm) with H_2_O-MeOH gradient to provide the chemicals in (**a**).

*<u>tert</u>*<u>-Butyl (4-propionamidobutyl)carbamate (**1a**):</u> amorphous yellow oil; ^1^H-NMR (CDCl_3_, 600 MHz) *δ* (ppm): 5.82 (1H, s, NH), 4.67 (1H, s, NH), 3.24 (2H, q, *J* = 6.0Hz), 3.12 (2H, q, *J* = 6.0 Hz), 2.18 (2H, q, *J* = 7.2 Hz), 1.51 (4H, m), 1.42 (9H, s), 1.13 (3H, t, *J* = 7.2 Hz); ^13^C-NMR (CDCl_3_, 150 MHz) δ (ppm): 174.0, 156.3, 79.3, 40.2, 39.2, 29.8, 28.5, 27.7, 26.8, 10.0; ESI-MS m/z 267.2 [M+Na]^+^.

*<u>tert</u>*<u>-Butyl (4-butyramidobutyl)carbamate (**2a**):</u> amorphous yellow oil; ^1^H-NMR (CDCl_3_, 600 MHz) δ (ppm): 5.81 (1H, s, NH), 4.67 (1H, s, NH), 3.25 (2H, q, *J* = 6.0 Hz), 3.12 (2H, m), 2.13 (2H, t, *J* = 7.2 Hz), 1.65 (2H, m), 1.51 (4H, m), 1.42 (9H, s), 0.92 (3H, t, *J* = 7.2 Hz); ^13^C-NMR (CDCl_3_, 150 MHz) δ (ppm): 173.2, 156.3, 79.3, 40.2, 39.1, 38.8, 28.5, 27.7, 26.8, 19.3, 13.9; ESI-MS m/z 281.2 [M+Na]^+^.

*<u>tert</u>*<u>-Butyl (4-valeramidobutyl)carbamate (**3a**):</u> amorphous yellow oil; ^1^H-NMR (CDCl_3_, 600 MHz) δ (ppm): 5.76 (1H, s, NH), 4.64 (1H, s, NH), 3.26 (2H, m), 3.13 (2H, m), 2.15 (2H, t, *J* = 7.8 Hz), 1.60 (2H, m), 1.51 (4H, m), 1.43 (9H, s), 1.33 (2H, m), 0.90 (3H, t, *J* = 7.2 Hz); ^13^C-NMR (CDCl_3_, 150 MHz) δ (ppm): 173.4, 156.3, 79.4, 40.2, 39.2, 36.7, 28.5, 28.0, 27.8, 26.9, 22.5, 13.9; ESI-MS m/z 295.2 [M+Na]^+^.

*<u>tert</u>*<u>-Butyl (4-myristamidobutyl)carbamate (**4a**):</u> amorphous yellow oil; ^1^H-NMR (CDCl_3_, 600 MHz) δ (ppm): 5.72 (1H, s, NH), 4.63 (1H, s, NH), 3.25 (2H, q, *J* = 6.0 Hz), 3.12 (2H, m), 2.14 (2H, t, *J* = 7.8 Hz), 1.61 (2H, q, *J* = 7.8 Hz), 1.51 (4H, m), 1.43 (9H, s), 1.24 (20H, s), 0.87 (3H, t, *J* = 7.2 Hz); ^13^C-NMR (CDCl_3_, 150 MHz) δ (ppm): 173.4, 156.3, 79.4, 40.2, 39.2, 37.0, 32.1, 29.8-29.5, 28.5, 27.8, 26.9, 26.0, 22.8, 14.3; ESI-MS m/z 421.4 [M+Na]^+^.

*<u>tert</u>*<u>-Butyl (4-oleamidobutyl)carbamate (**5a**):</u> amorphous yellow oil; ^1^H-NMR (CDCl_3_, 600 MHz) δ (ppm): 5.74 (1H, s, NH), 5.33 (2H, m), 4.64 (1H, s, NH), 3.25 (2H, q, *J* = 6.0 Hz), 3.12 (2H, q, *J* = 6.0 Hz), 2.14 (2H, t, *J* = 7.2 Hz), 1.99 (4H, q, *J* = 6.6 Hz), 1.60 (2H, m), 1.51 (4H, m), 1.43 (9H, s), 1.29-1.25 (20H,m), 0.87 (3H, d, *J* = 6.6 Hz); ^13^C-NMR (CDCl_3_, 150 MHz) δ (ppm): 173.4, 156.3, 130.1, 129.9, 79.3, 40.2, 39.2, 37.0, 32.0, 29.9-29.3, 28.5, 27.78, 27.3 (2 x CH_2_), 26.8, 25.9, 22.8, 14.2; ESI-MS m/z 475.4 [M+Na]^+^.

*<u>tert</u>*<u>-Butyl (4-stearamidobutyl)carbamate (**6a**):</u> amorphous yellow oil; ^1^H-NMR (CDCl_3_, 600 MHz) δ (ppm): 5.71 (1H, s, NH), 4.63 (1H, s, NH), 3.26 (2H, q, *J* = 6.0 Hz), 3.12 (2H, q, *J* = 6.0 Hz,), 2.15 (2H, t, *J* = 7.8 Hz), 1.61 (2H, m), 1.51 (4H, m), 1.43 (9H, s), 1.25 (28H, s), 0.87 (3H, t, *J* = 7.8 Hz); ^13^C-NMR (CDCl_3_, 150 MHz) δ (ppm): 173.4, 156.3, 79.4, 40.2, 39.2, 37.0, 32.1, 29.8-29.5, 28.5, 27.8, 26.9, 26.0, 22.8, 14.3; ESI-MS m/z 477.4 [M+Na]^+^.

###### General Procedure B

To a solution of (**a**) in anhydrous CH_2_Cl_2_ (5 mL), TFA (5 mL) was added and stirred for 1 h at room temperature. Then excess reagent and solvent were removed under vacuum. The resulting product was neutralized by saturated NaHCO_3_, extracted with CH_2_Cl_2_ (2 x 10 mL), dried over Na_2_SO_4_, filtered, and evaporated in vacuo. The residue was purified using silica gel flash chromatography (CH_2_Cl_2_/MeOH gradient) to yield final compound.

*<u>N</u>*<u>-propionoylputrescine (**1**):</u> amorphous yellow oil; ^1^H-NMR (CD_3_OD, 600 MHz) δ (ppm): 3.21 (2H, t, *J* = 7.2 Hz), 2.94 (2H, t, *J* = 7.2 Hz), 2.20 (2H, q, *J* = 7.2Hz), 1.66 (2H, m), 1.58 (2H, m), 1.12 (3H, t, *J* = 7.2 Hz);^13^C-NMR (CD_3_OD, 150 MHz) δ (ppm): 177.4, 40.5, 39.6, 30.3, 27.6, 26.0, 10.7; ESI-HRMS *m/z*: [M+H]^+^ calculated for C_7_H_17_N_2_O 145.1334; found 145.1335.

*<u>N</u>*<u>-butyroylputrescine (**2**):</u> amorphous yellow oil; ^1^H-NMR (CD_3_OD, 600 MHz) δ (ppm): 3.21 (2H, t, *J* = 7.2 Hz), 2.94 (2H, t, *J* = 7.2 Hz), 2.16 (2H, t, *J* = 7.2Hz), 1.67 (2H, m), 1.63 (2H, m), 1.58 (2H, m), 0.94 (3H, t, *J* = 7.2 Hz); ^13^C-NMR (CD_3_OD, 150 MHz) δ (ppm): 176.4, 40.5, 39.6, 39.2, 27.6, 26.0, 20.5, 14.1; ESI-HRMS *m/z*: [M+H]^+^ calculated for C_8_H_19_N_2_O 159.1492; found 159.1491.

*<u>N</u>*<u>-valeroylputrescine (**3**):</u> amorphous yellow oil; ^1^H-NMR (CD_3_OD, 600 MHz) δ (ppm): 3.20 (2H, t, *J* = 7.2 Hz), 2.94 (2H, t, *J* = 7.2 Hz), 2.19 (2H, t, *J* = 7.2 Hz), 1.67 (2H, m), 1.57 (4H, m), 1.34 (2H, m), 0.93 (3H, t, *J* = 7.2 Hz); ^13^C-NMR (CD_3_OD, 150 MHz) δ (ppm): 176.6, 40.4, 39.6, 37.0, 29.3, 27.5, 26.0, 23.5, 14.3; ESI-HRMS *m/z*: [M+H]^+^ calculated for C_9_H_21_N_2_O 173.1647; found 173.1648.

*<u>N</u>*<u>-myristorylputrescine (**4**):</u> amorphous yellow oil; ^1^H-NMR (CD_3_OD, 600 MHz) δ (ppm): 3.20 (2H, t, *J* = 7.2 Hz), 2.94 (2H, t, *J* = 7.2 Hz), 2.18 (2H, t, *J* = 7.2 Hz), 1.66 (2H, m), 1.59 (4H, m), 1.29 (20H, m), 0.90 (3H, t, *J* = 7.2 Hz); ^13^C-NMR (CD_3_OD, 150 MHz) δ (ppm): 176.7, 40.5, 39.5, 37.3, 33.2, 30.9 – 30.5, 27.6, 27.2, 26.0, 23.9, 14.6; ESI-HRMS *m/z*: [M+H]^+^ calculated for C_18_H_39_N_2_O 299.3054; found 299.3055.

*<u>N</u>*<u>-oleoylputrescine (**5**):</u> amorphous yellow oil; ^1^H-NMR (CD_3_OD, 600 MHz) δ (ppm): 5.35 (2H, m), 3.79 (2H, t, *J* = 7.2 Hz), 3.29 (2H, m), 3.15 (2H, t, *J* = 7.2 Hz), 2.59 (2H, t, *J* = 7.2 Hz), 2.03 (4H, m), 1.65 (2H, m), 1.38 – 1.29 (22H, m), 0.90 (3H, t, *J* = 7.2 Hz); ^13^C-NMR (CD_3_OD, 150 MHz) δ (ppm): 177.8, 131.1, 130.9, 56.6, 43.7, 42.2, 36.9, 36.6, 33.2, 31.0 – 30.3, 28.3 (2 x CH_2_), 26.2, 25.8, 23.9, 14.60; ESI-HRMS *m/z*: [M+H]^+^ calculated for C_22_H_45_N_2_O 353.3526; found 353.3522.

*<u>N</u>*<u>-stearoylputrescine (**6**):</u> amorphous yellow oil; ; ^1^H-NMR (CD_3_OD, 600 MHz) δ (ppm): 3.20 (2H, t, *J* = 7.2 Hz), 2.94 (2H, t, *J* = 7.2 Hz), 2.18 (2H, t, *J* = 7.2 Hz), 1.66 (2H, m), 1.58 (4H, m), 1.29 (26H, m), 0.90 (3H, t, *J* = 7.2 Hz); ^13^C-NMR (CD_3_OD, 150 MHz) δ (ppm): 176.7, 40.3, 39.5, 37.3, 33.2, 30.9 – 30.5, 27.7, 27.2, 26.0, 23.9, 14.6; ESI-HRMS *m/z*: [M+H]^+^ calculated for C_22_H_47_N_2_O 355.3683; found 355.3689.

##### Synthesis of *N*-oleoylputrescine hydrochloride

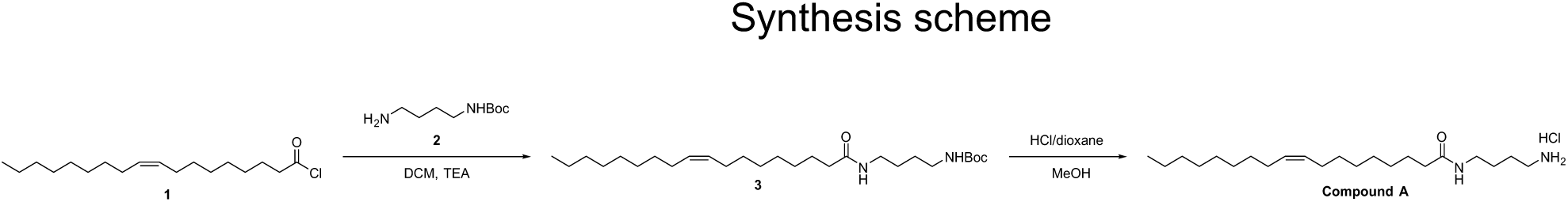

<u>Step1</u>

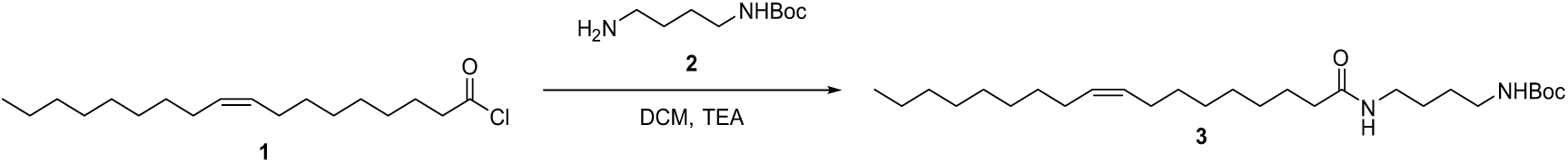

*N*-oleoylputrescine hydrochloride (NOP) was prepared by BioDuro LLC; tert-Butyl (4-aminobutyl)carbamate and oleic acid conjugation, followed by tert-butyloxycarbonyl (Boc) cleavage and adding hydrogen chloride. To a solution of *tert*-Butyl (4-aminobutyl)carbamate **(2)** (6.26g, 33.23mmol) in anhydrous dichloromethane (DCM, 50 mL) at 0 °C was added a solution of oleoyl chloride (**1**) (10.0 g, 33.23 mmol) in anhydrous DCM (50 mL) and triethylamine (TEA, 6.71 g, 66.46 mmol) dropwise. The mixture was stirred at 25 °C for 3 h. The reaction was quenched with water (100 mL), extracted with dichloromethane (DCM, 100 mL x 3). The combined organic phases were washed with brine, dried with anhydrous sodium sulfate and filtered. The filtrate was concentrated under reduced pressure to provide a crude product, which was purified through a reverse-phase chromatography column using a 75%-95% acetonitrile in water gradient to afford tert-butyl (4-oleamidobutyl)carbamate (**3**) (7.4 g, yield: 49.2%) as a white solid. ESI-MS *m/z* = 353.4 [M-Boc] ^+^, 475.4 [M+Na] ^+^

<u>Step2</u>

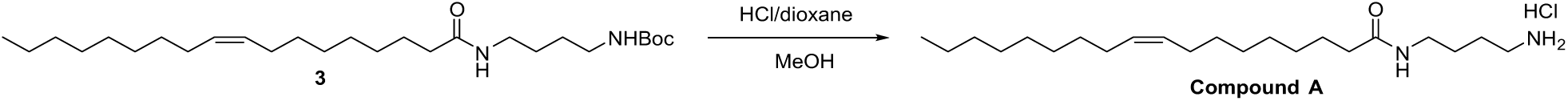

To a solution of tert-butyl (4-oleamidobutyl)carbamate (**3**) (7.4 g, 16.34 mmol) in methanol (60 mL) was added HCl/1,4-dioxane (15 mL, 4 M in dioxane). The reaction mixture was stirred at 25 °C for 2h. The reaction mixture was concentrated under the reduced pressure. The residue was diluted with water and then lypholized to afford N-(4-aminobutyl) oleamide hydrochloric acid (**Compound A**) (5.2 g, yield: 81.2%) as a white solid.

<u>Instrumentation</u>: ^1^H NMR spectra was recorded on a Bruker Advance III (400 MHz) instrument and mass spectrometry was recorded on Agilent Technologies 1200 Series, G6110A.

Data for ^1^H NMR are reported as follows: chemical shift (δ ppm), integration, multiplicity (s = singlet, br s = broad singlet, d = doublet, t = triplet, q = quartet, m = multiplet), and coupling constant (Hz). ^1^H NMR (DMSO-*d_6_*, 400 MHz) δ 7.08 (t, *J* = 5.2 Hz, 1H), 7.73 (brs, 3H), 5.36-5.28 (m, 2H), 3.03 (dd, *J_1_* = 6.4 Hz, *J_2_* = 12.8 Hz, 2H), 2.79-2.74 (m, 2H), 2.04-1.97 (m, 6H), 1.52-1.39 (m, 6H), 1.24 (s, 20H), 0.85 (t, *J* = 6.8 Hz, 3H); ESI-MS *m/z* 353.4 [M+H]^+ 1^H and ^13^C NMR and HRESIMS spectra data of intermediate during synthesis and NPP, NBP, NVP, NMP, NOP, and NSP synthetic standards generated in the laboratory of Prof. Jon Clardy lab at Harvard Medical School.

**Supplemental Data 1:**
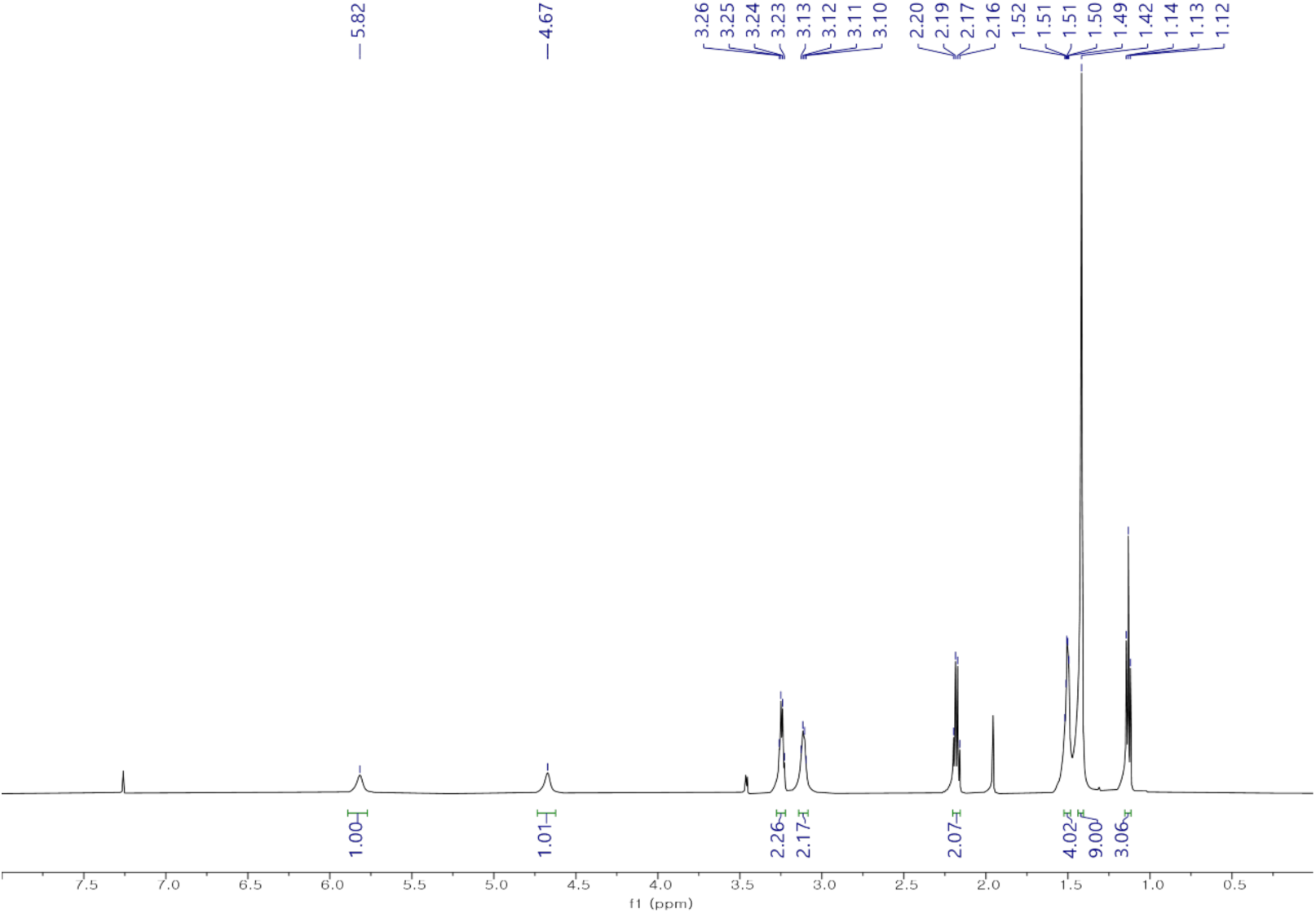
^1^H NMR spectrum of compound **1a** in CDCl_3_ (600 MHz).

**Supplemental Data 2:**
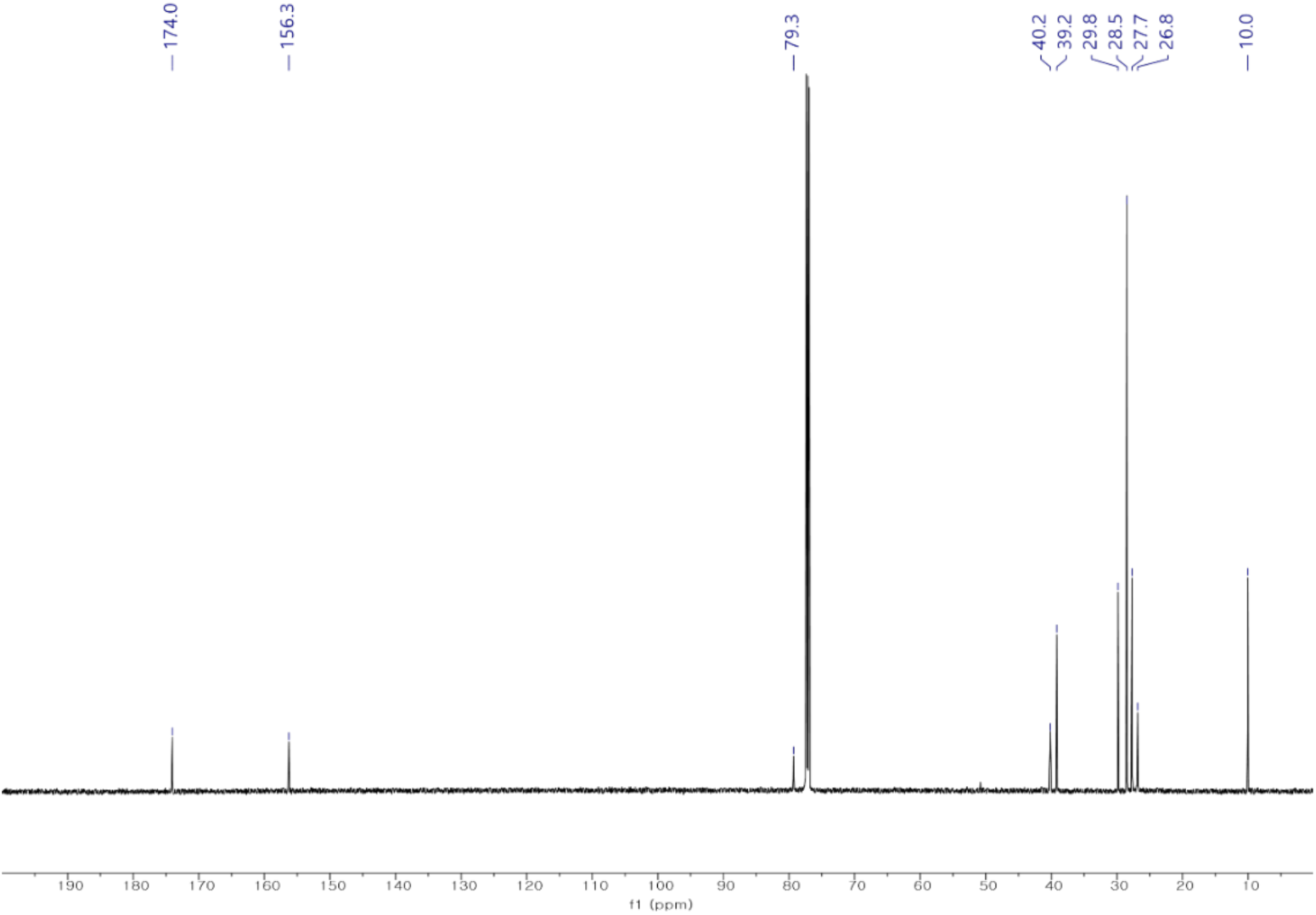
^13^C NMR spectrum of compound **1a** in CDCl_3_ (150 MHz).

**Supplemental Data 3:**
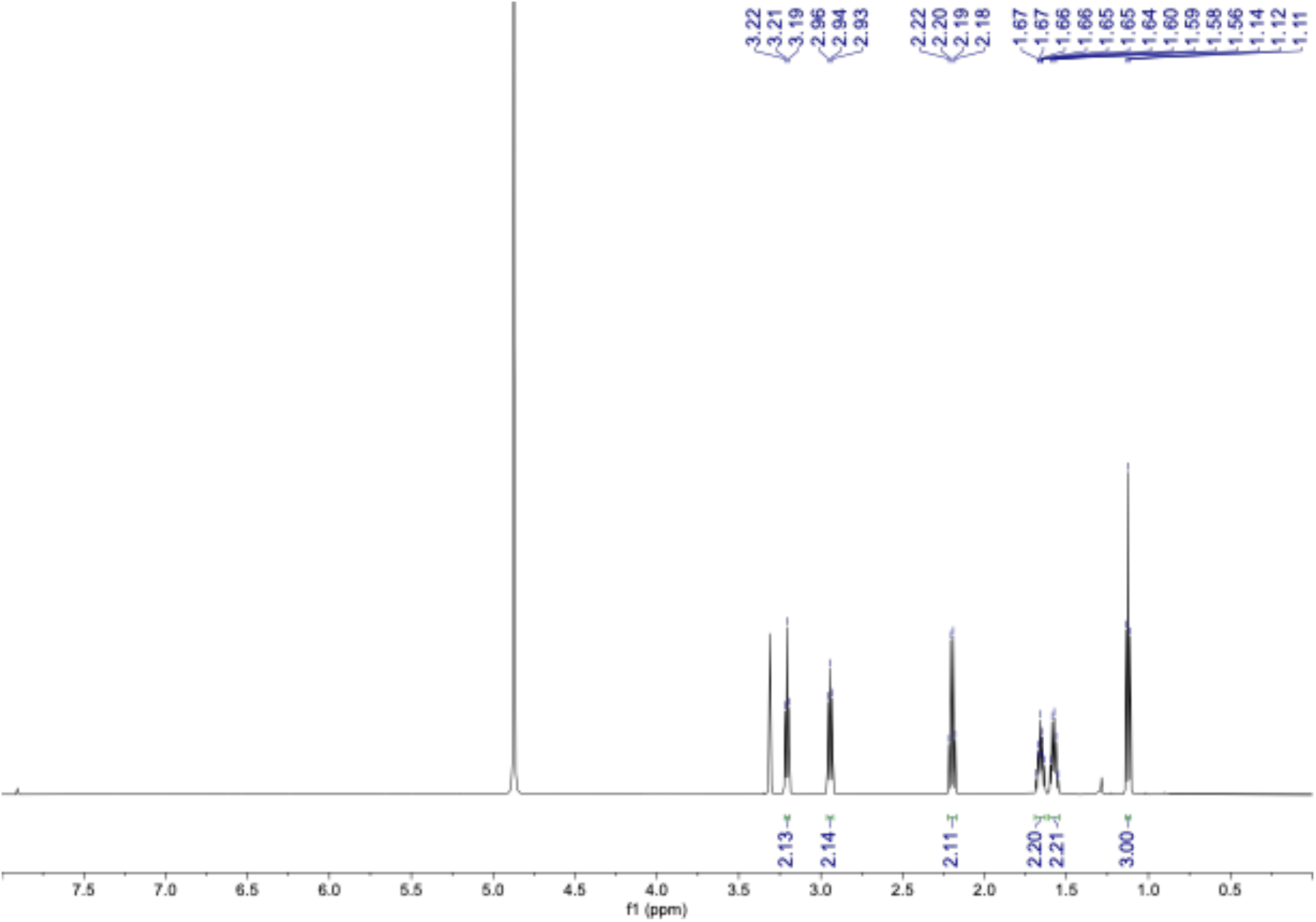
^1^H NMR spectrum of compound **1** in CD_3_OD (600 MHz).

**Supplemental Data 4:**
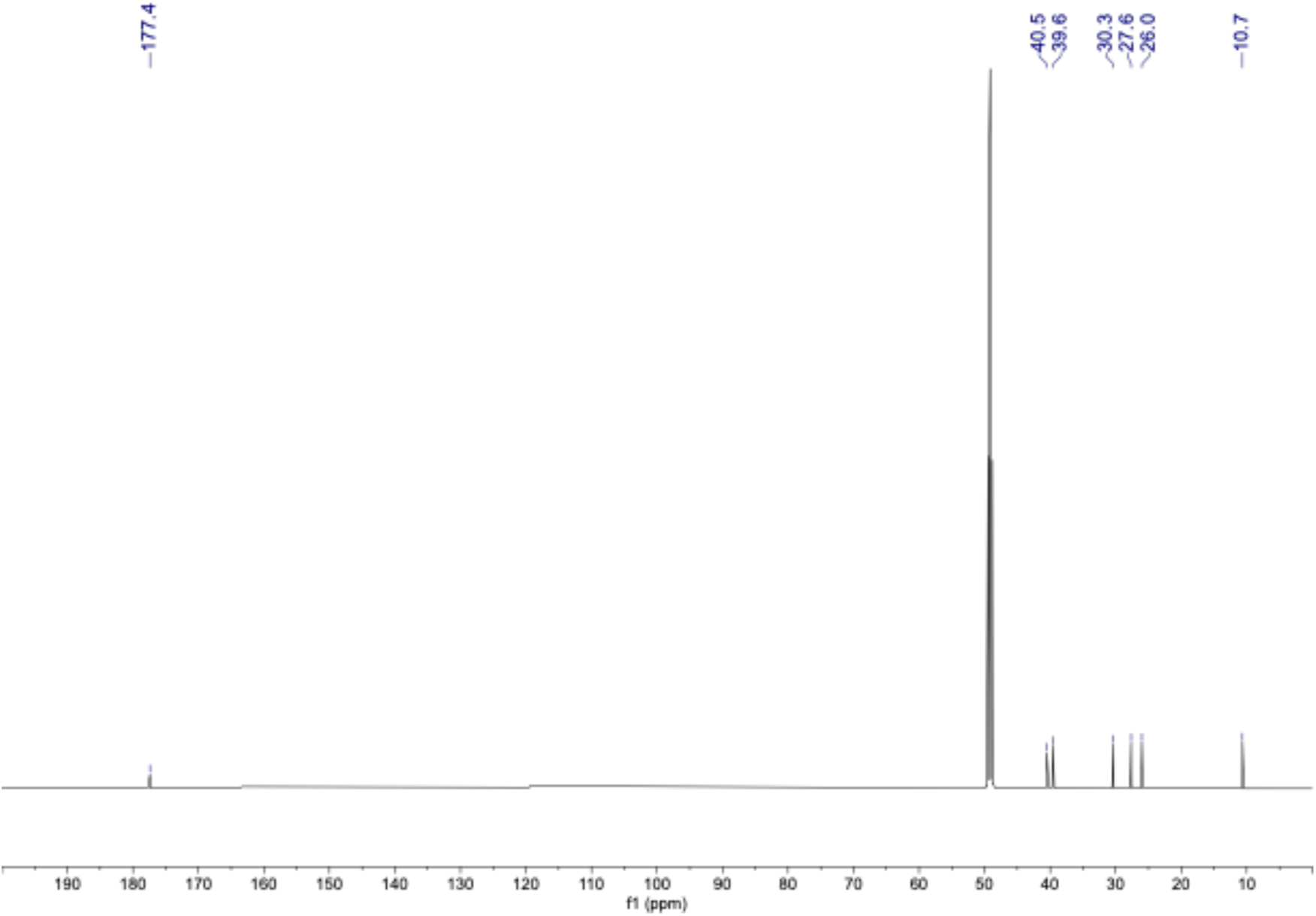
^13^C NMR spectrum of compound **1** in CD_3_OD (150 MHz).

**Supplemental Data 5:**
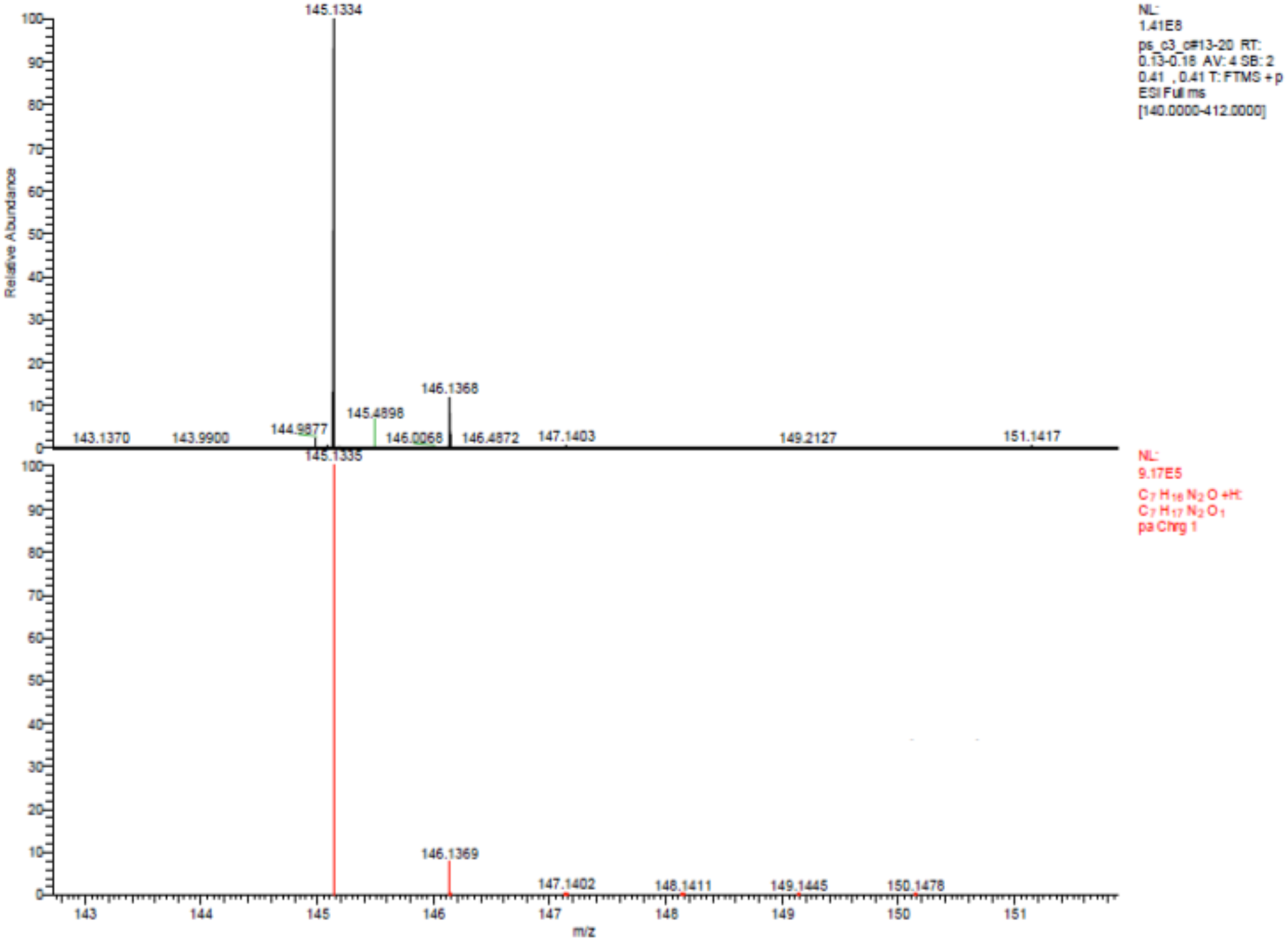
HRESIMS spectrum of compound **1**.

**Supplemental Data 6:**
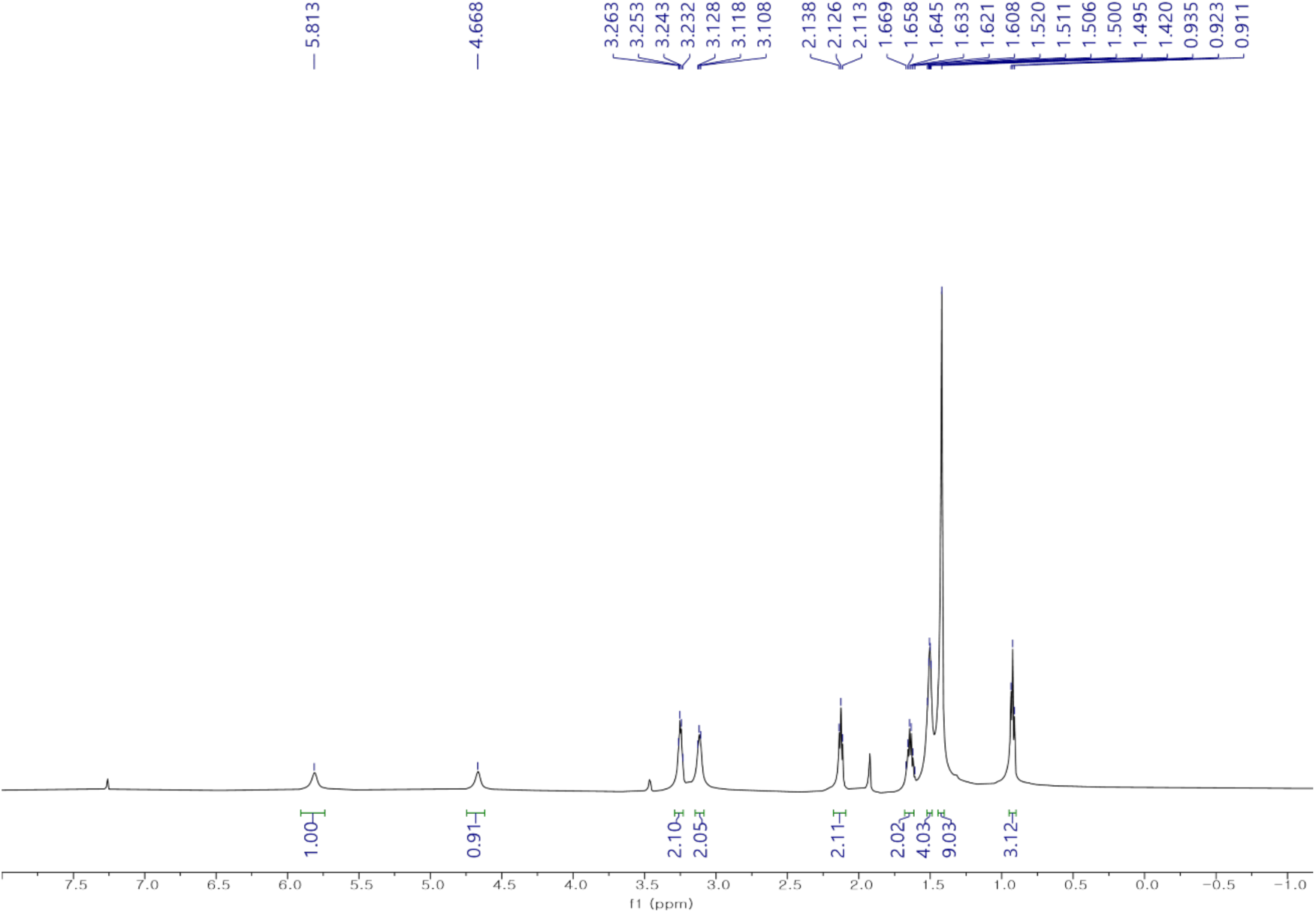
^1^H NMR spectrum of compound **2a** in CDCl_3_ (600 MHz).

**Supplemental Data 7:**
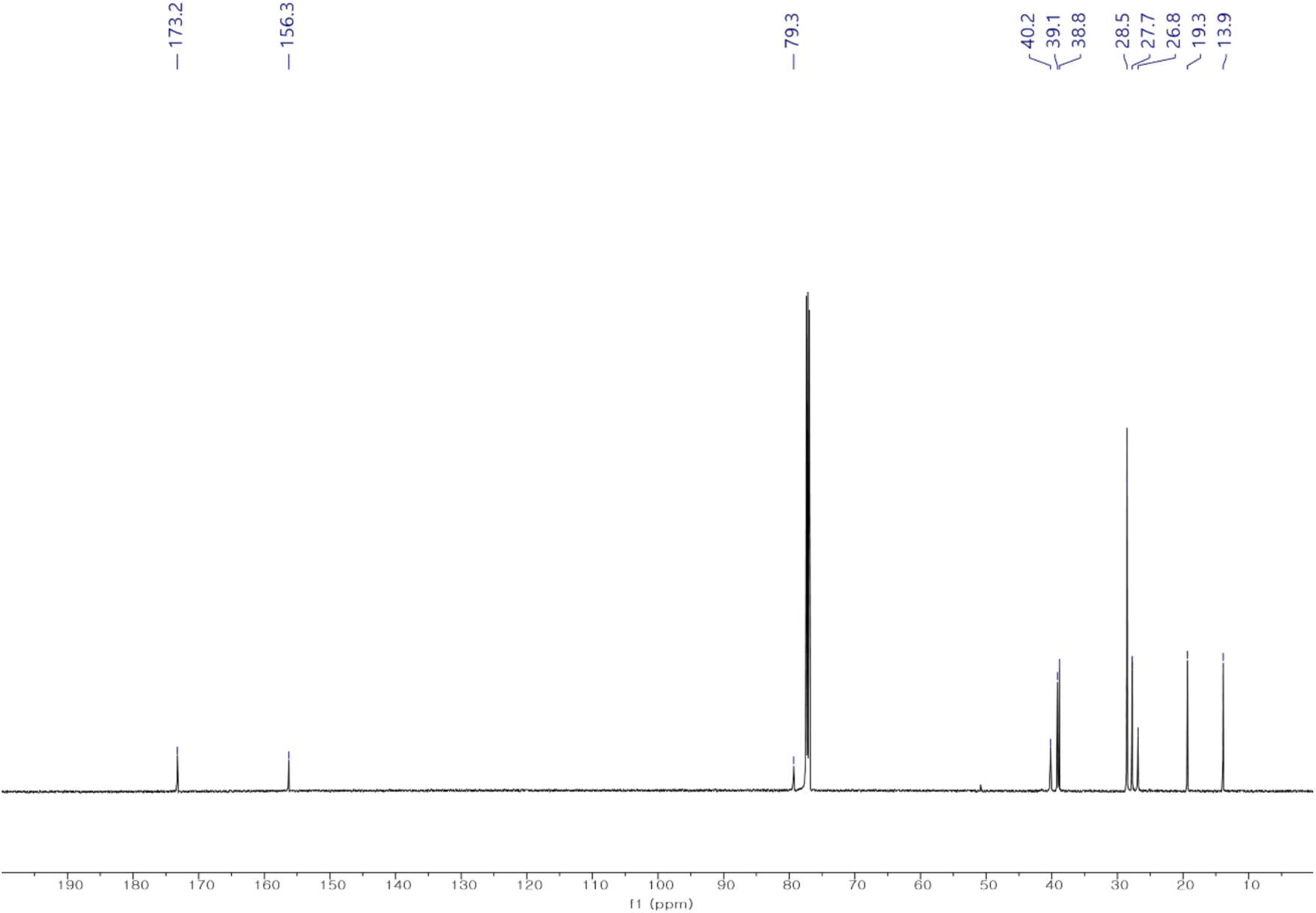
^13^C NMR spectrum of compound **2a** in CDCl_3_ (150 MHz).

**Supplemental Data 8:**
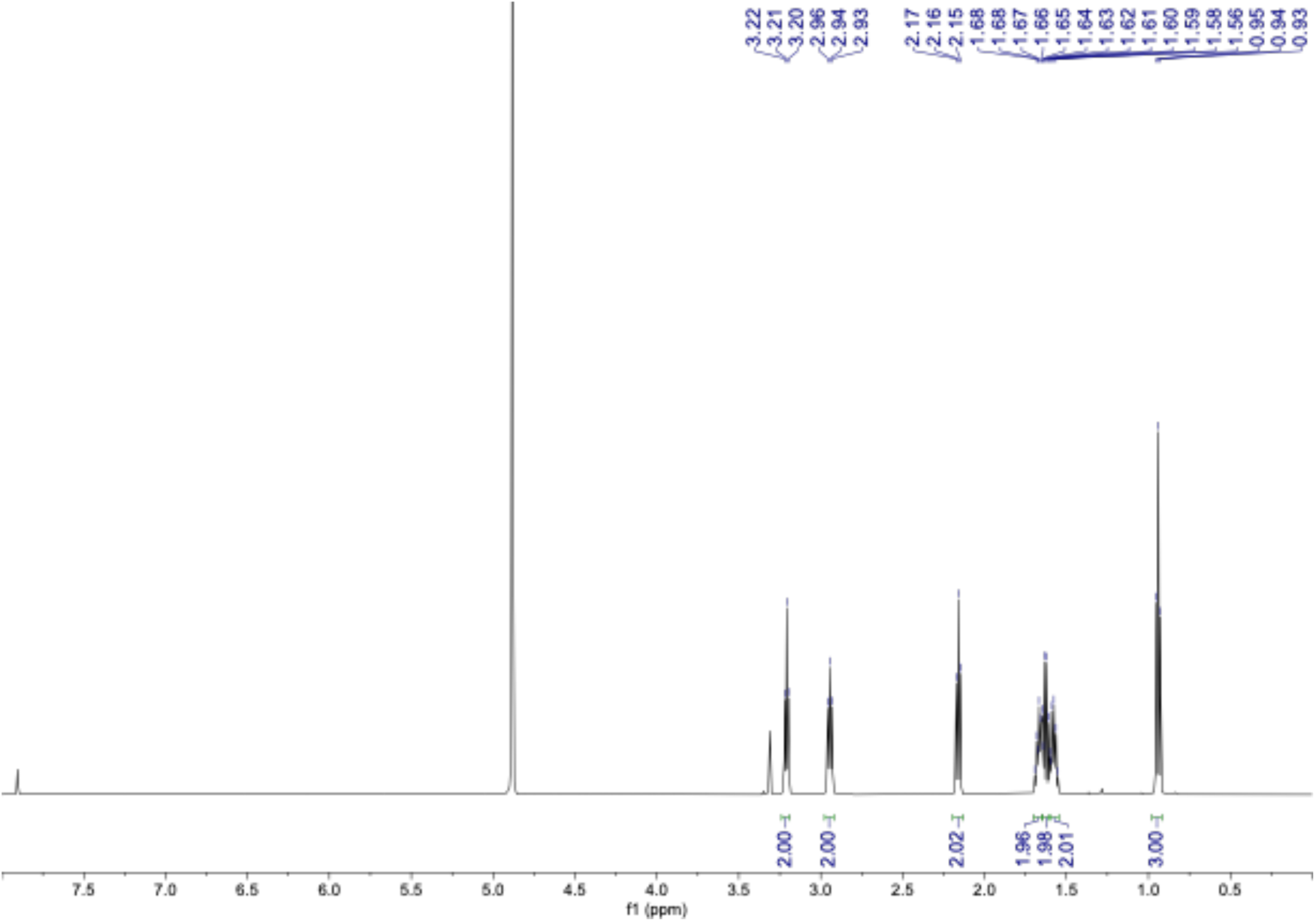
^1^H NMR spectrum of compound **2** in CD_3_OD (600 MHz).

**Supplemental Data 9:**
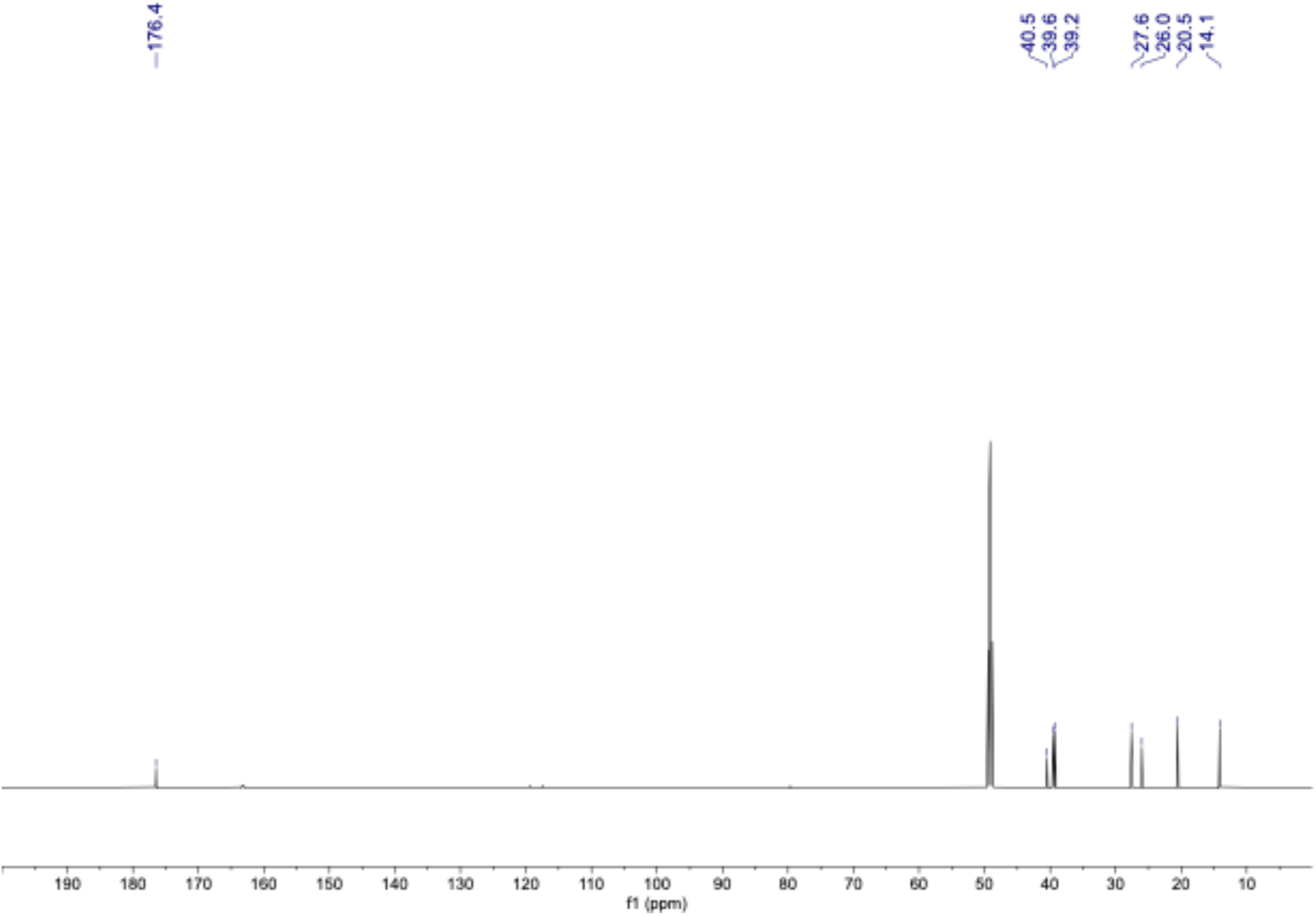
^13^C NMR spectrum of compound **2** in CD_3_OD (150 MHz).

**Supplemental Data 10:**
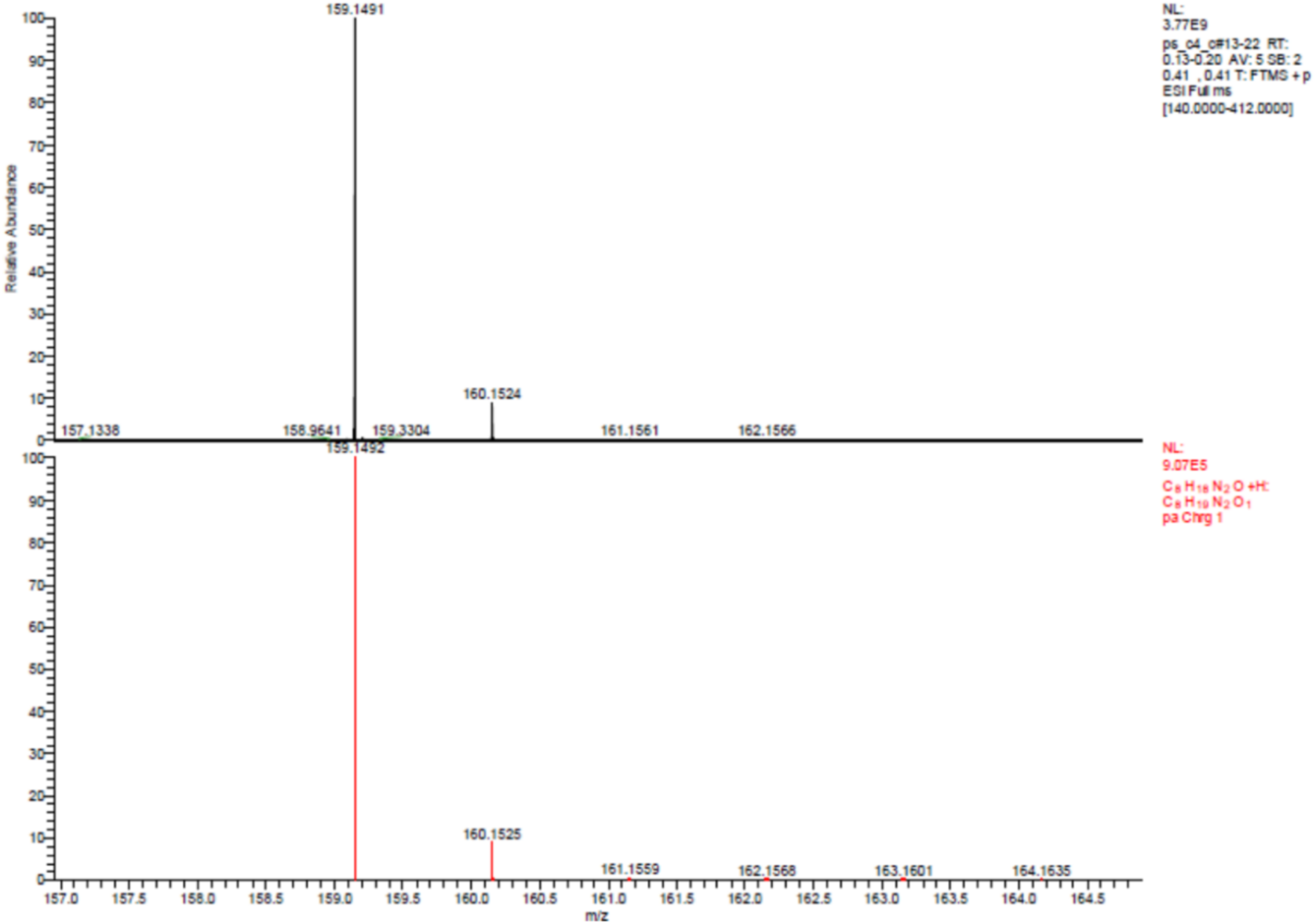
HRESIMS spectrum of compound **2**.

**Supplemental Data 11:**
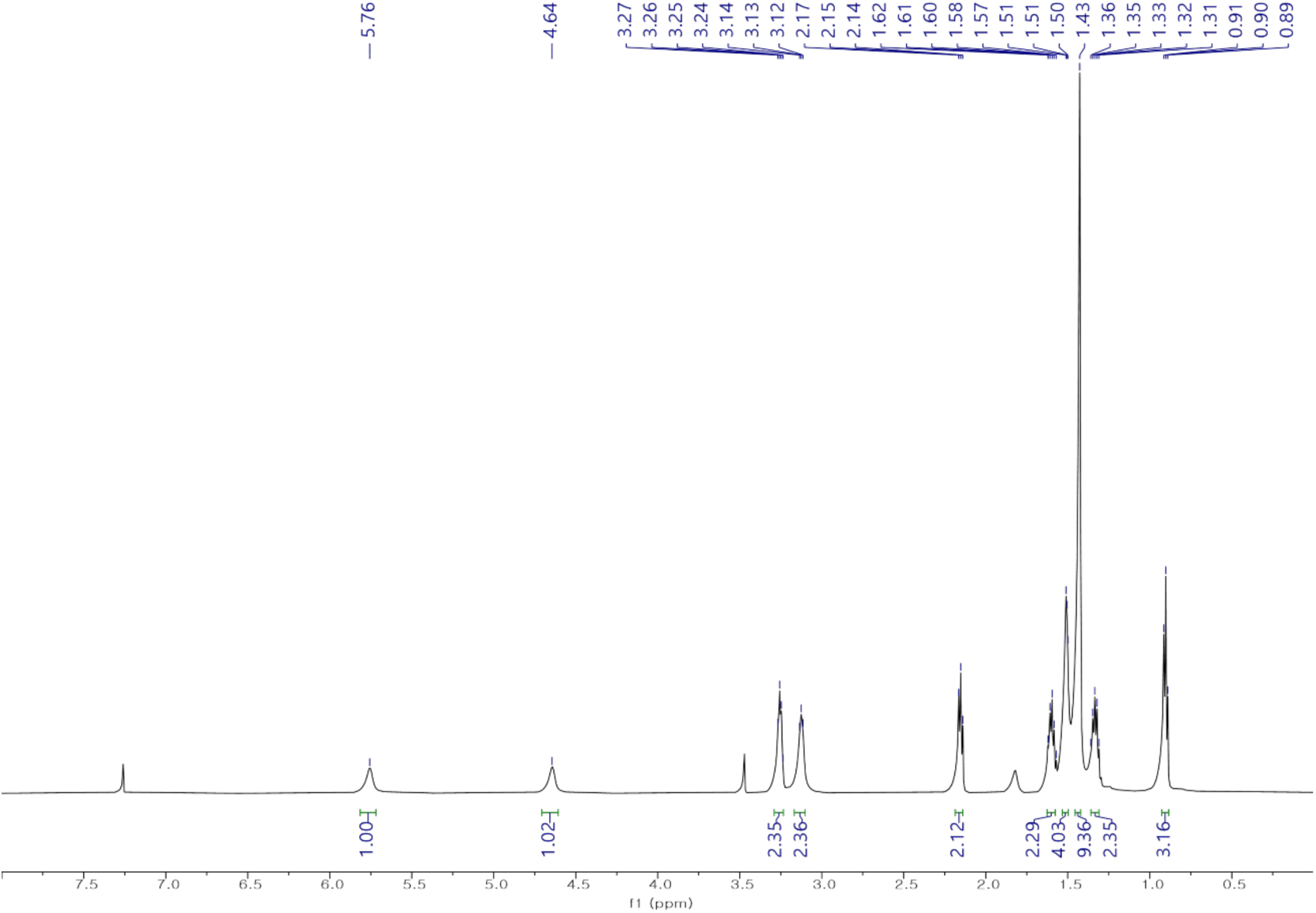
^1^H NMR spectrum of compound **3a** in CDCl_3_ (600 MHz).

**Supplemental Data 12:**
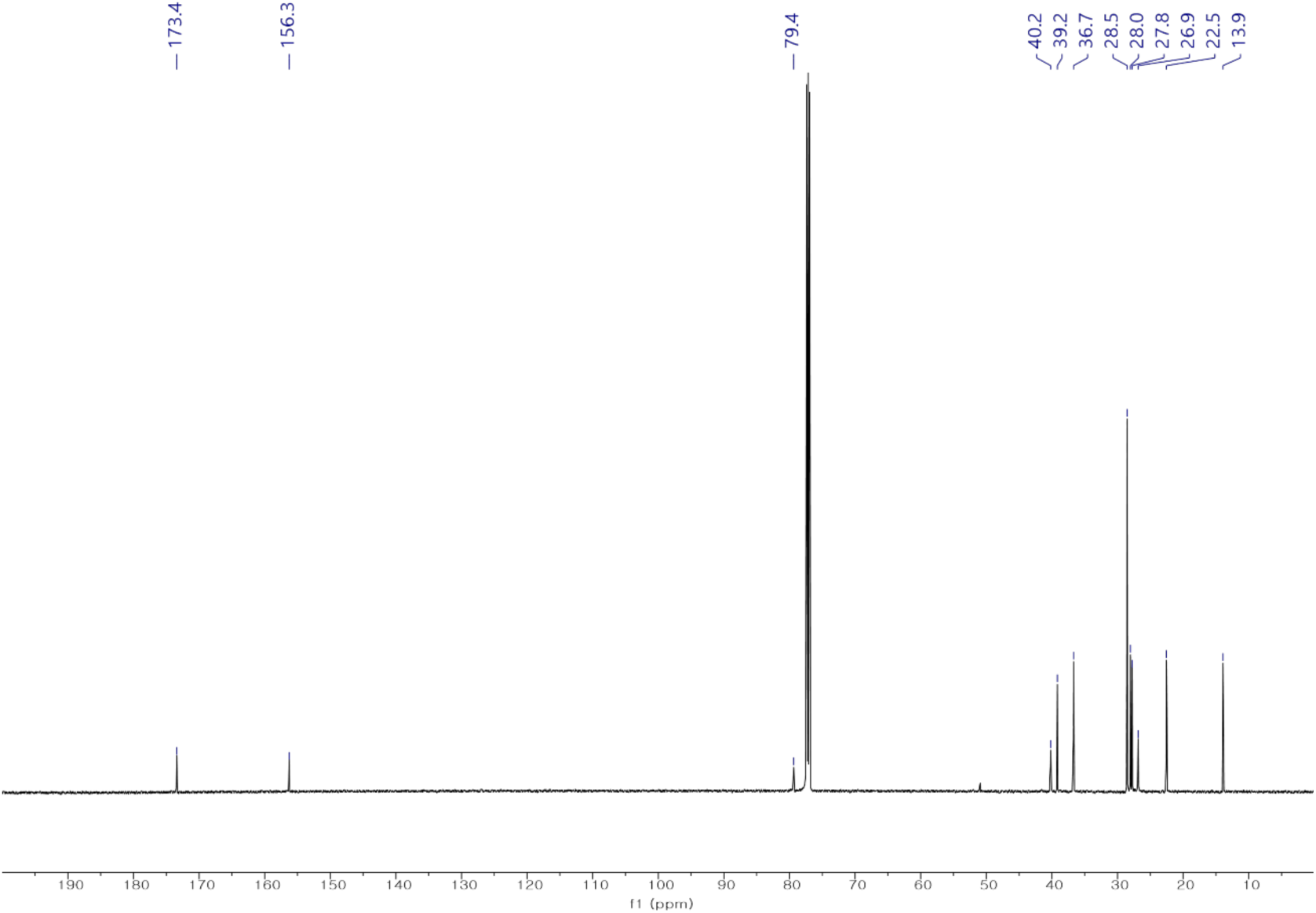
^13^C NMR spectrum of compound **3a** in CDCl_3_ (150 MHz).

**Supplemental Data 13:**
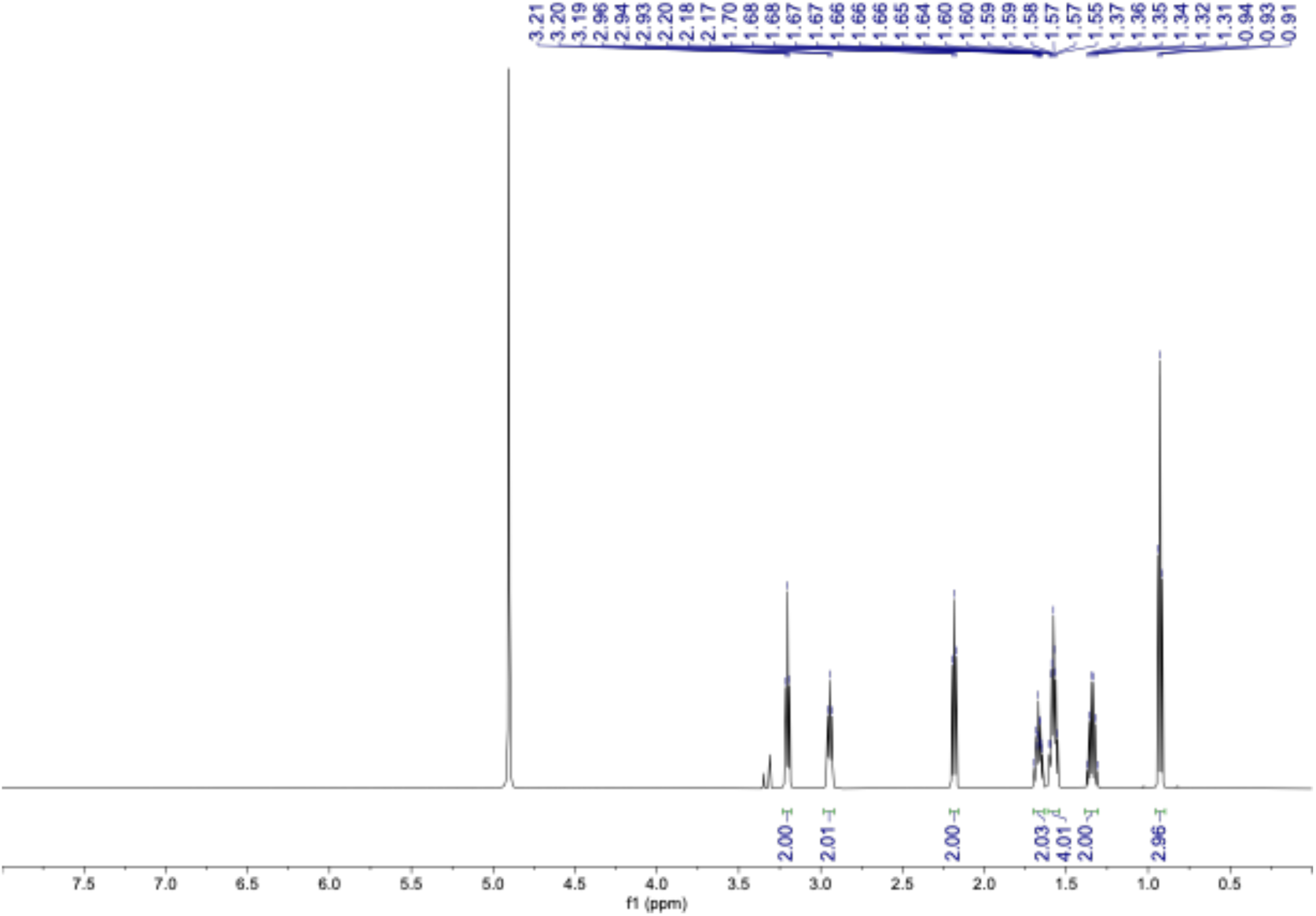
^1^H NMR spectrum of compound **3** in CD_3_OD (600 MHz).

**Supplemental Data 14:**
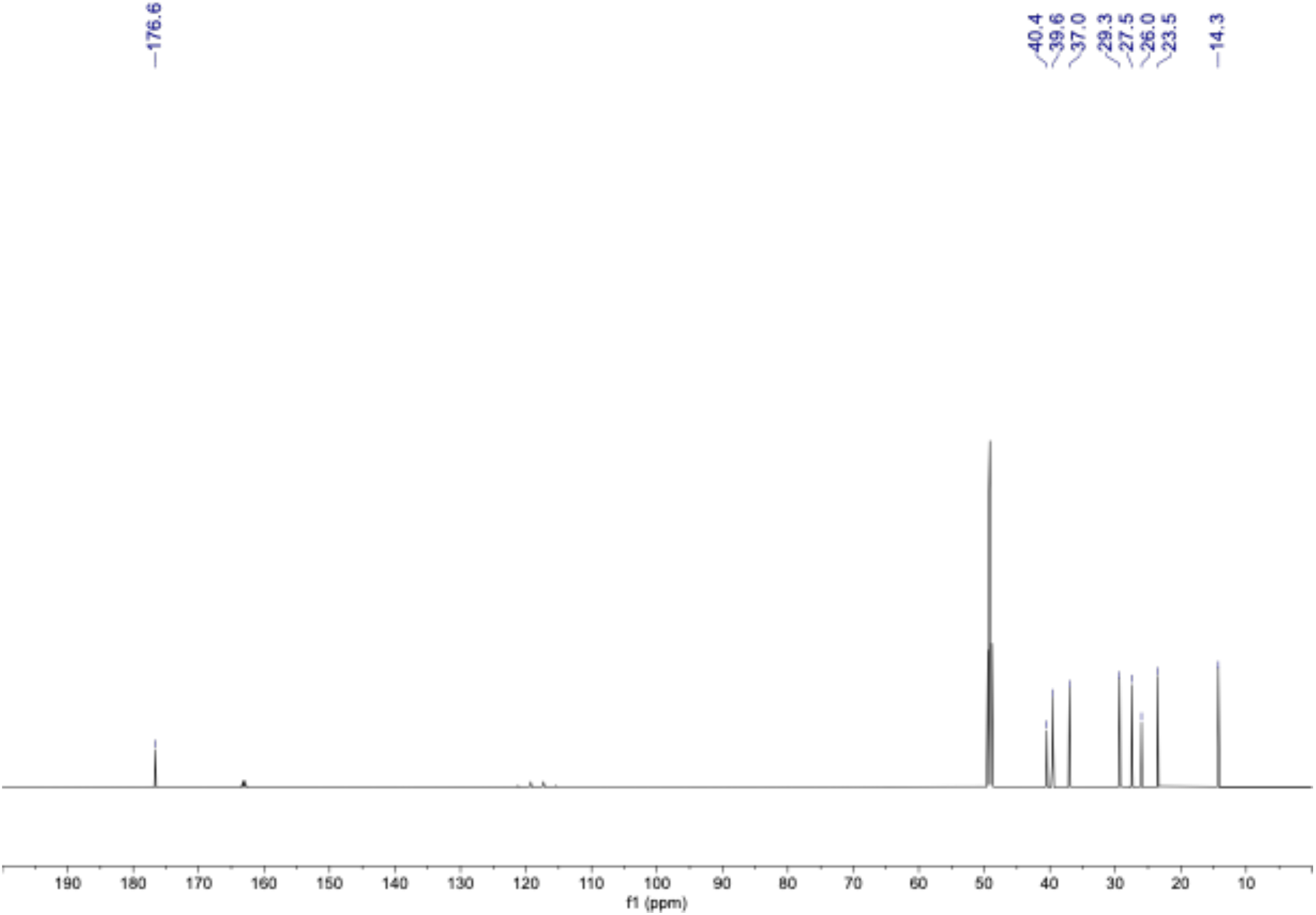
^13^C NMR spectrum of compound **3** in CD_3_OD (150 MHz).

**Supplemental Data 15:**
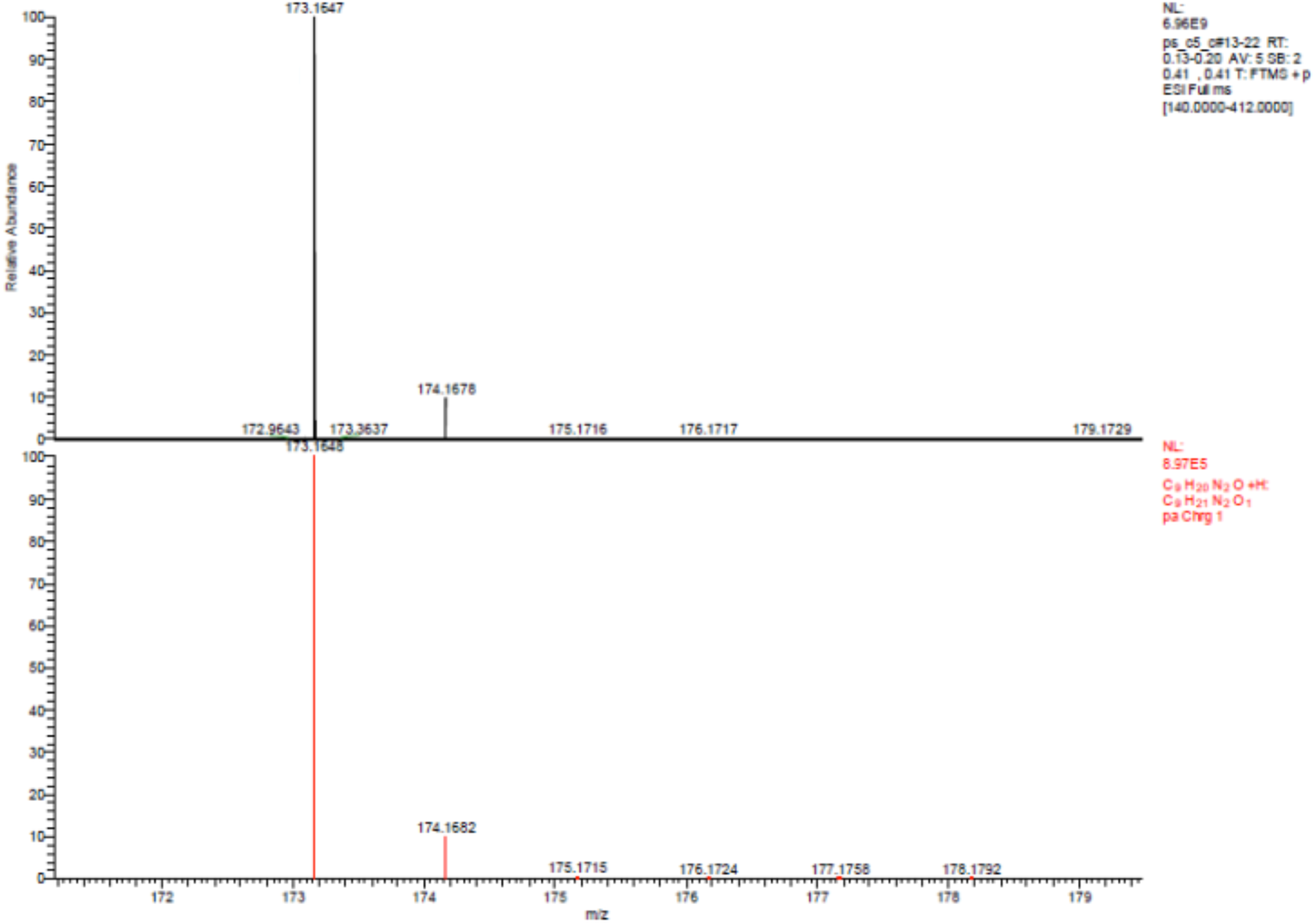
HRESIMS spectrum of compound **3**.

**Supplemental Data 16:**
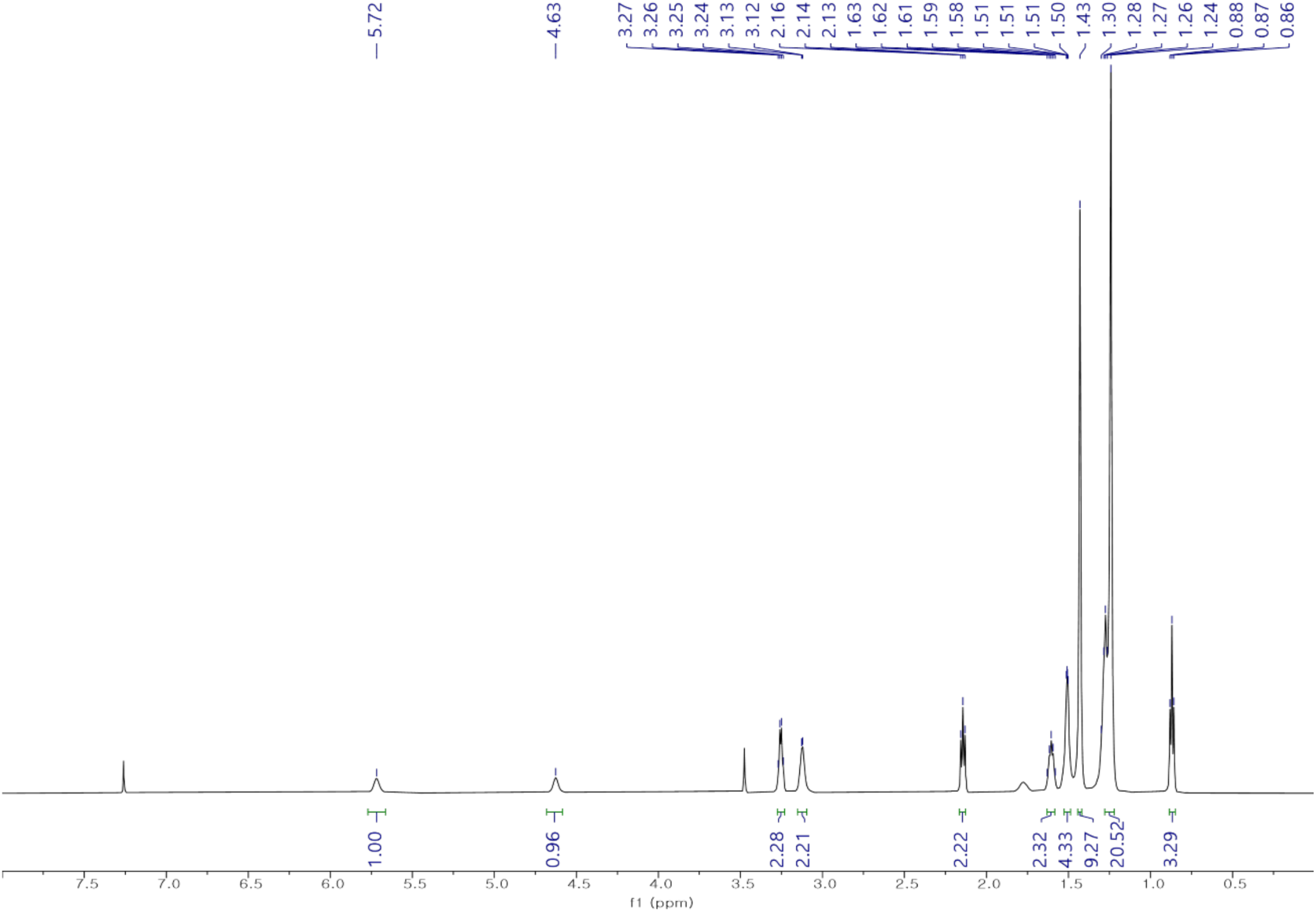
^1^H NMR spectrum of compound **4a** in CDCl_3_ (600 MHz).

**Supplemental Data 17:**
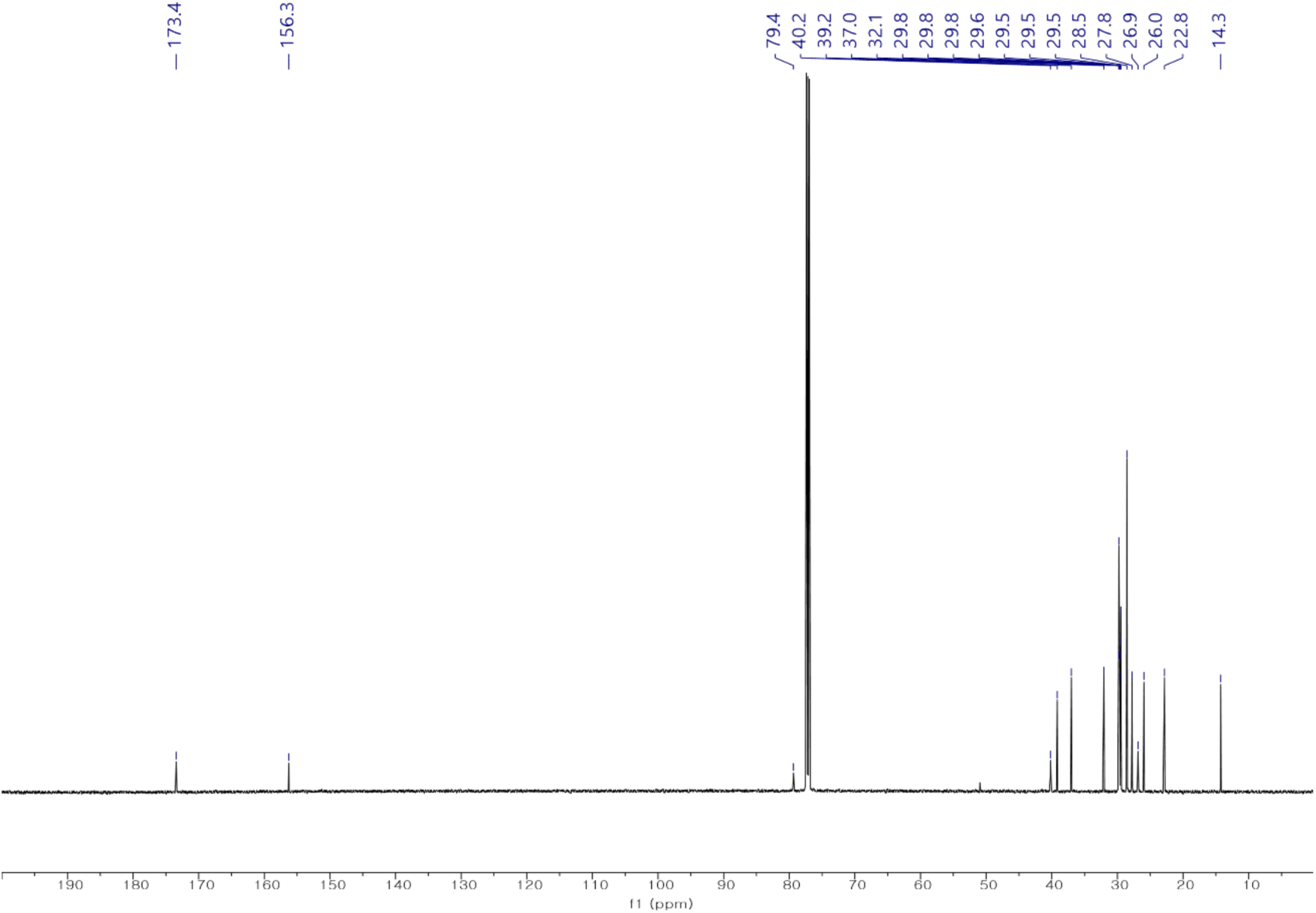
^13^C NMR spectrum of compound **4a** in CDCl_3_ (150 MHz).

**Supplemental Data 18:**
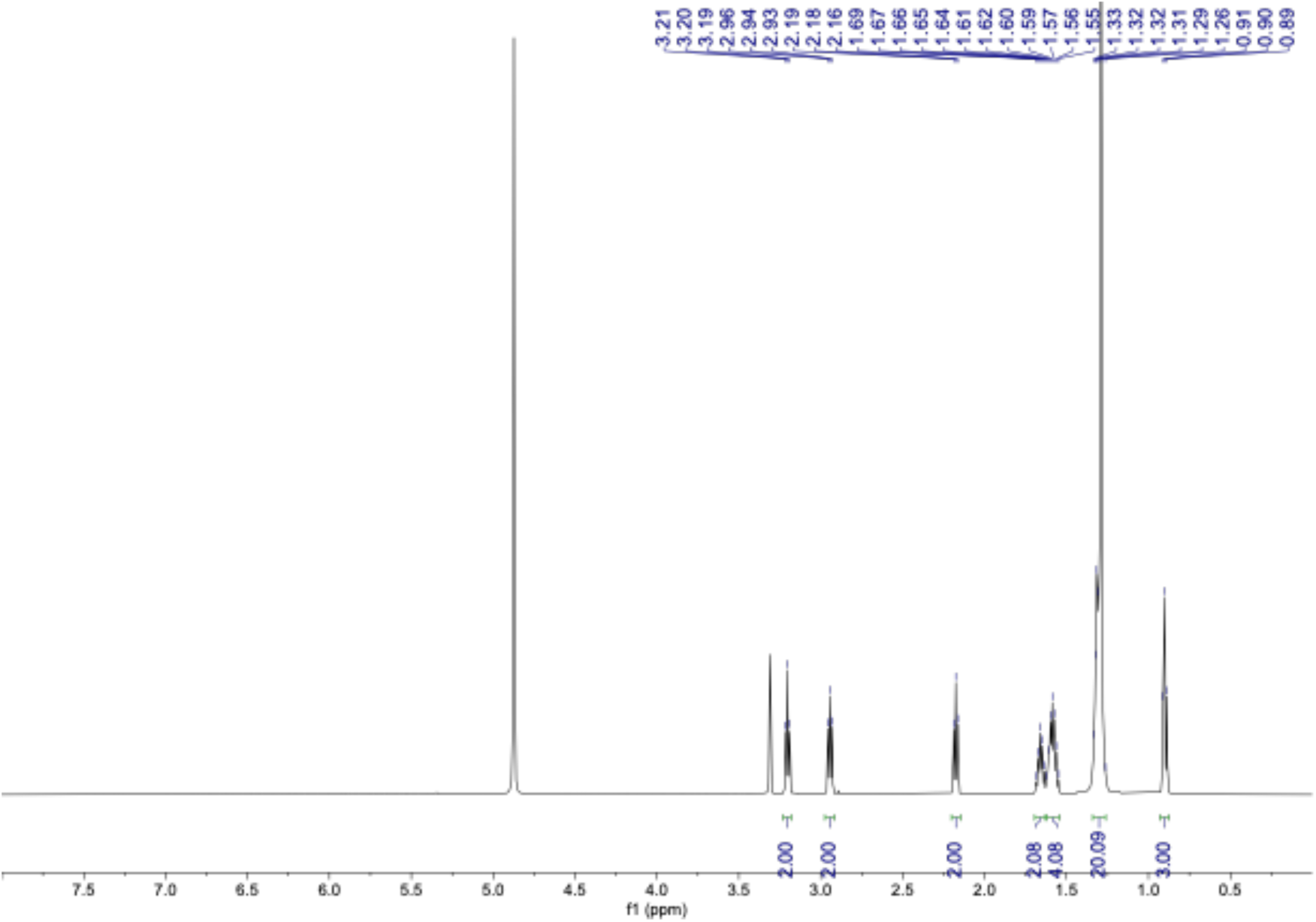
^1^H NMR spectrum of compound **4** in CD_3_OD (600 MHz).

**Supplemental Data 19:**
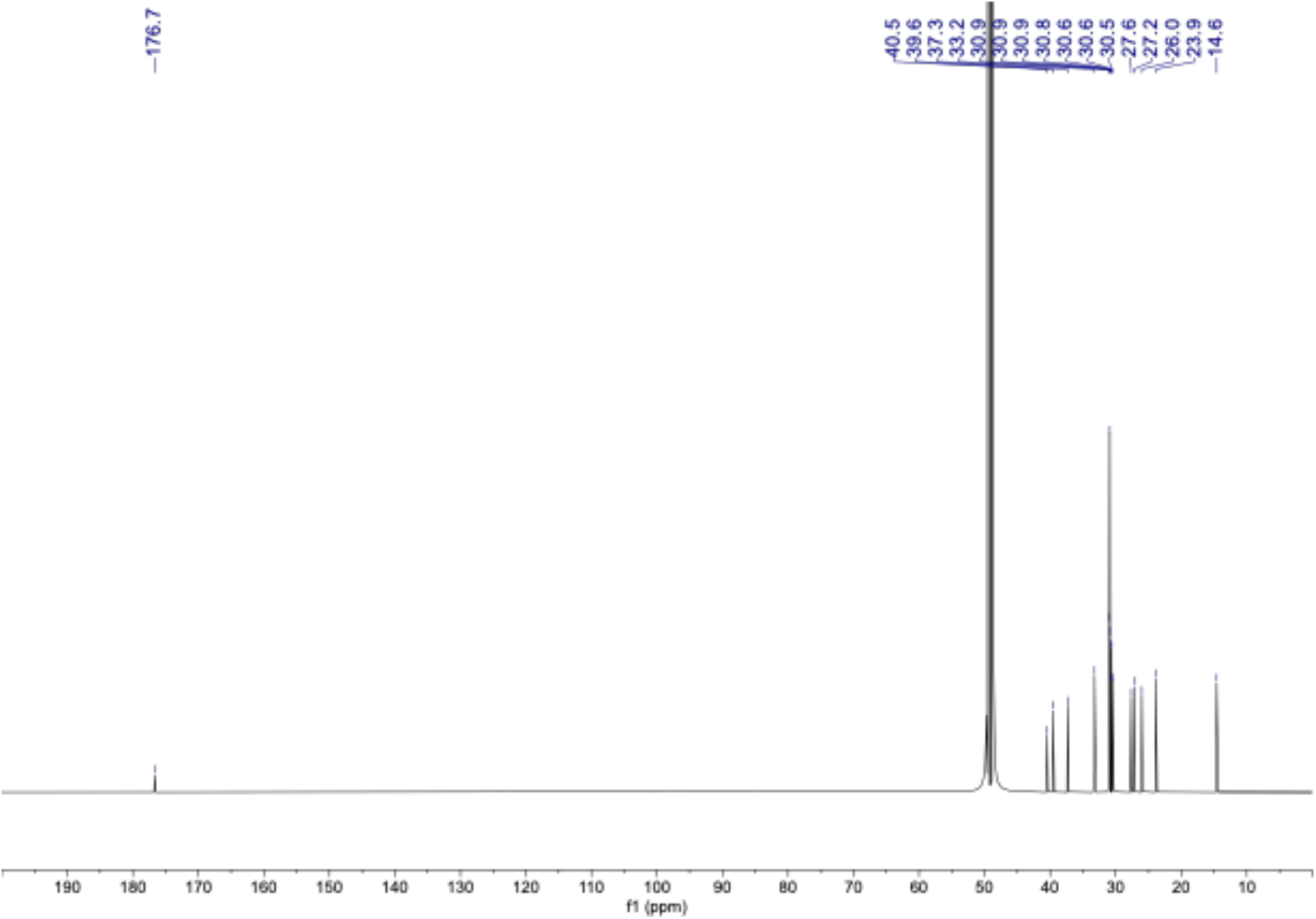
^13^C NMR spectrum of compound **4** in CD_3_OD (150 MHz).

**Supplemental Data 20:**
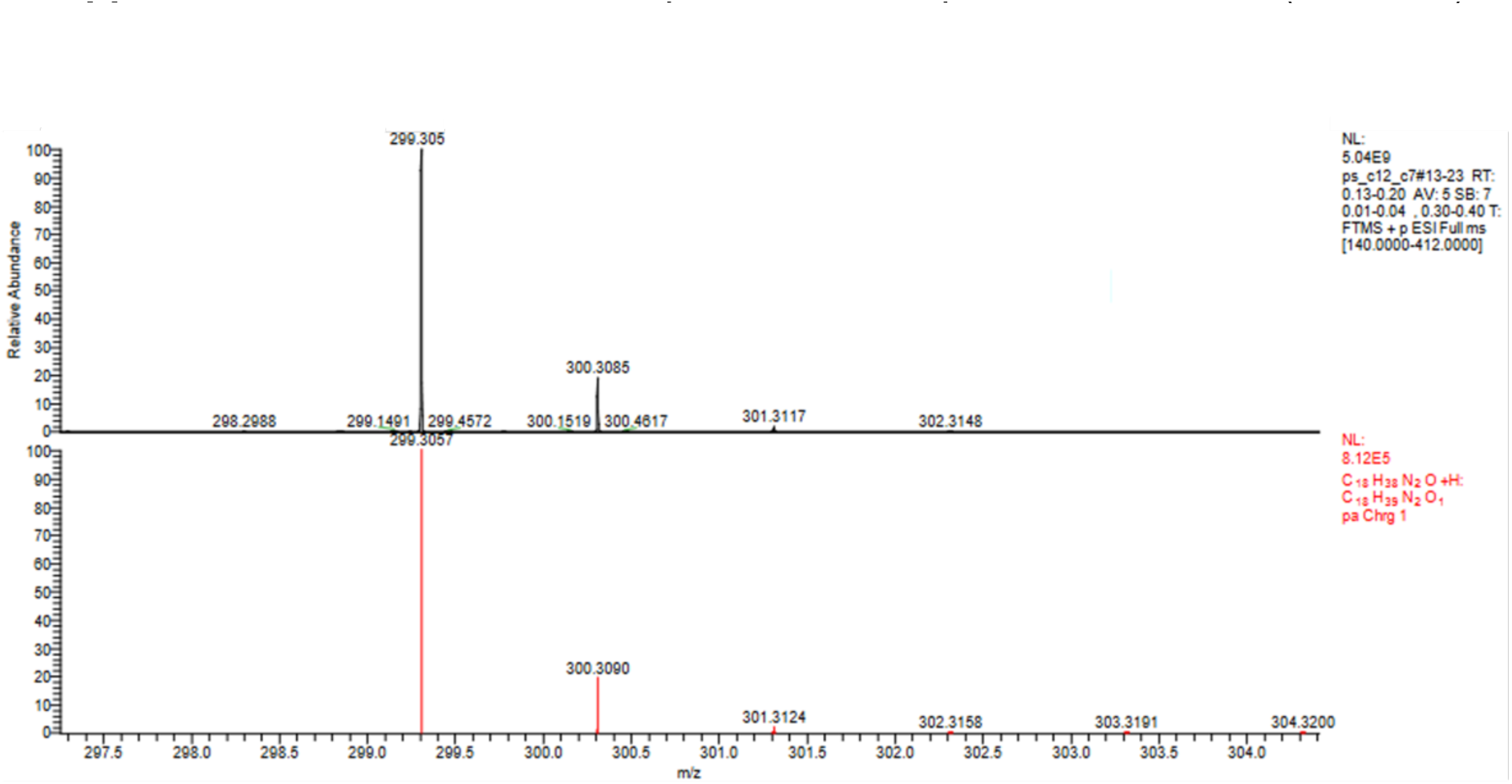
HRESIMS spectrum of compound **4**.

**Supplemental Data 21:**
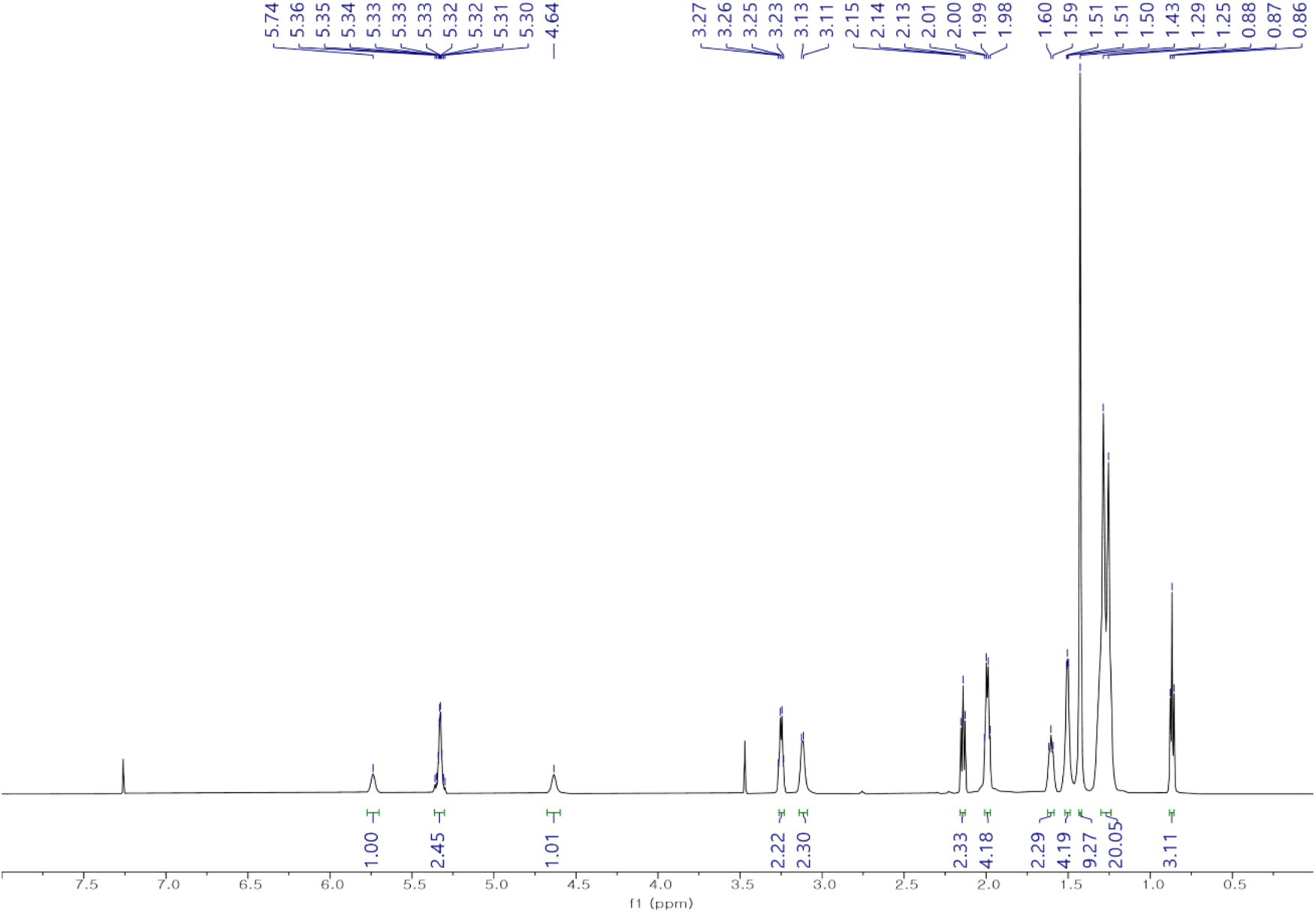
^1^H NMR spectrum of compound **5a** in CDCl_3_ (600 MHz).

**Supplemental Data 22:**
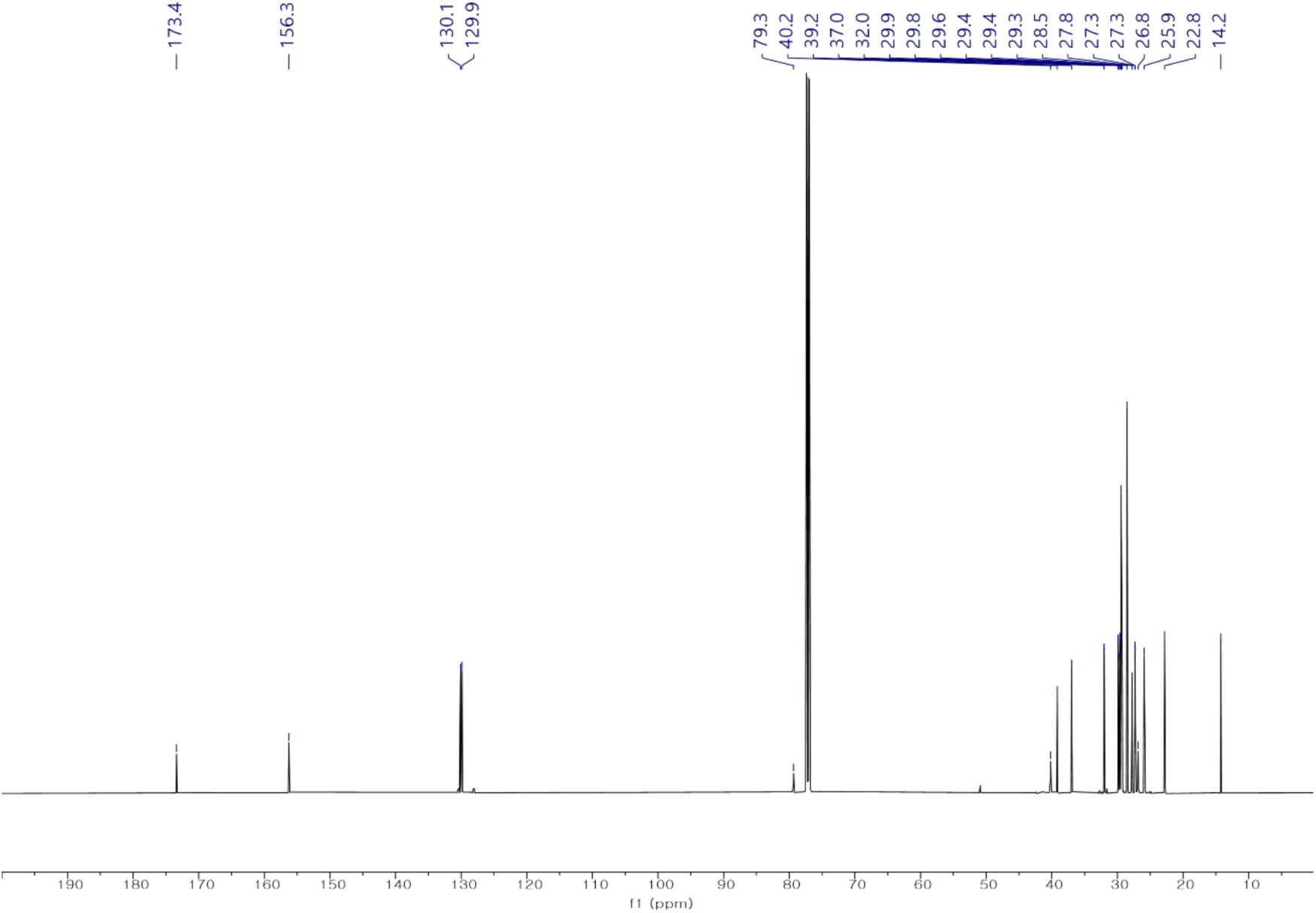
^13^C NMR spectrum of compound **5a** in CDCl_3_ (150 MHz).

**Supplemental Data 23:**
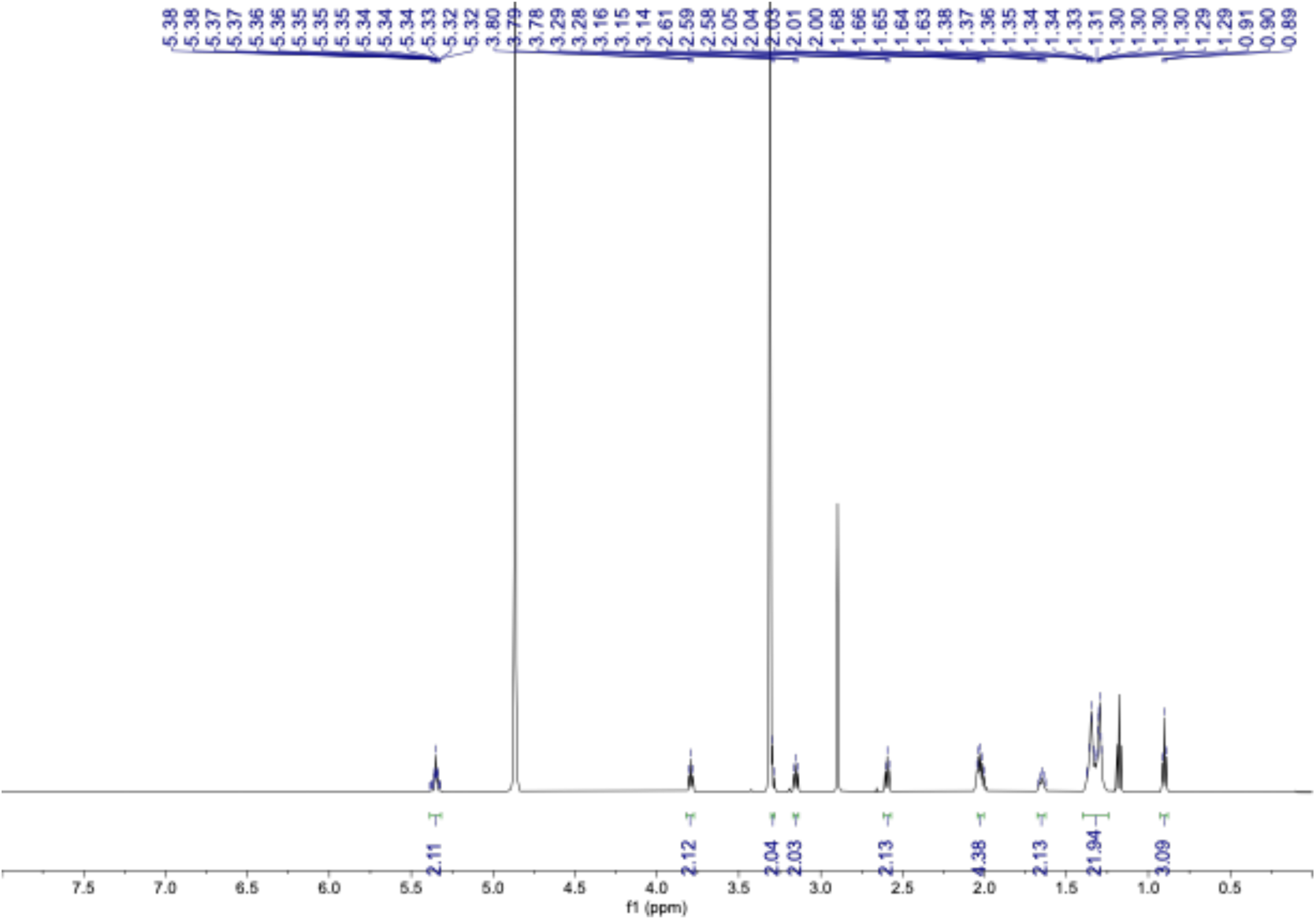
^1^H NMR spectrum of compound **5** in CD_3_OD (600 MHz).

**Supplemental Data 24:**
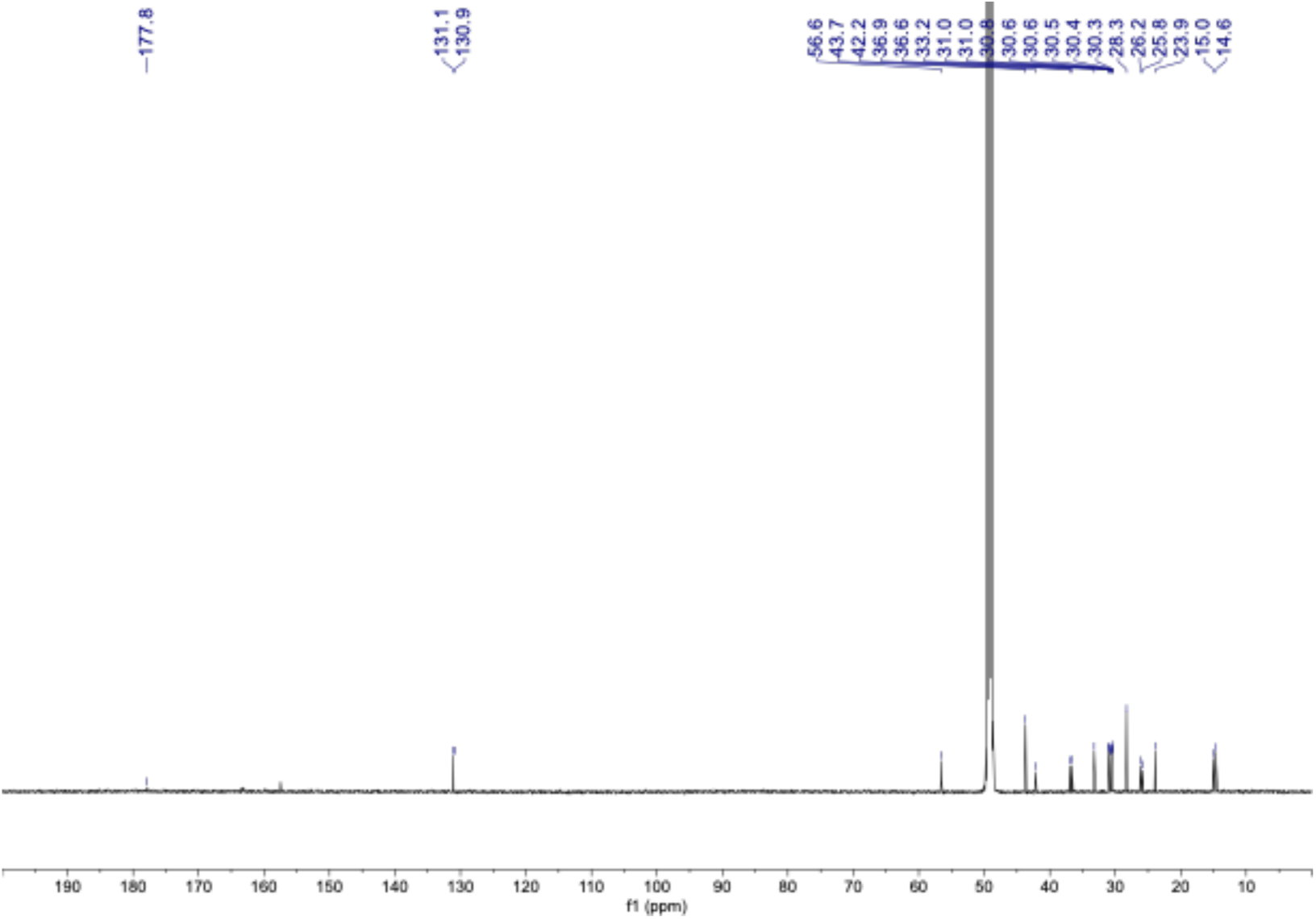
^13^C NMR spectrum of compound **5** in CD_3_OD (150 MHz).

**Supplemental Data 25:**
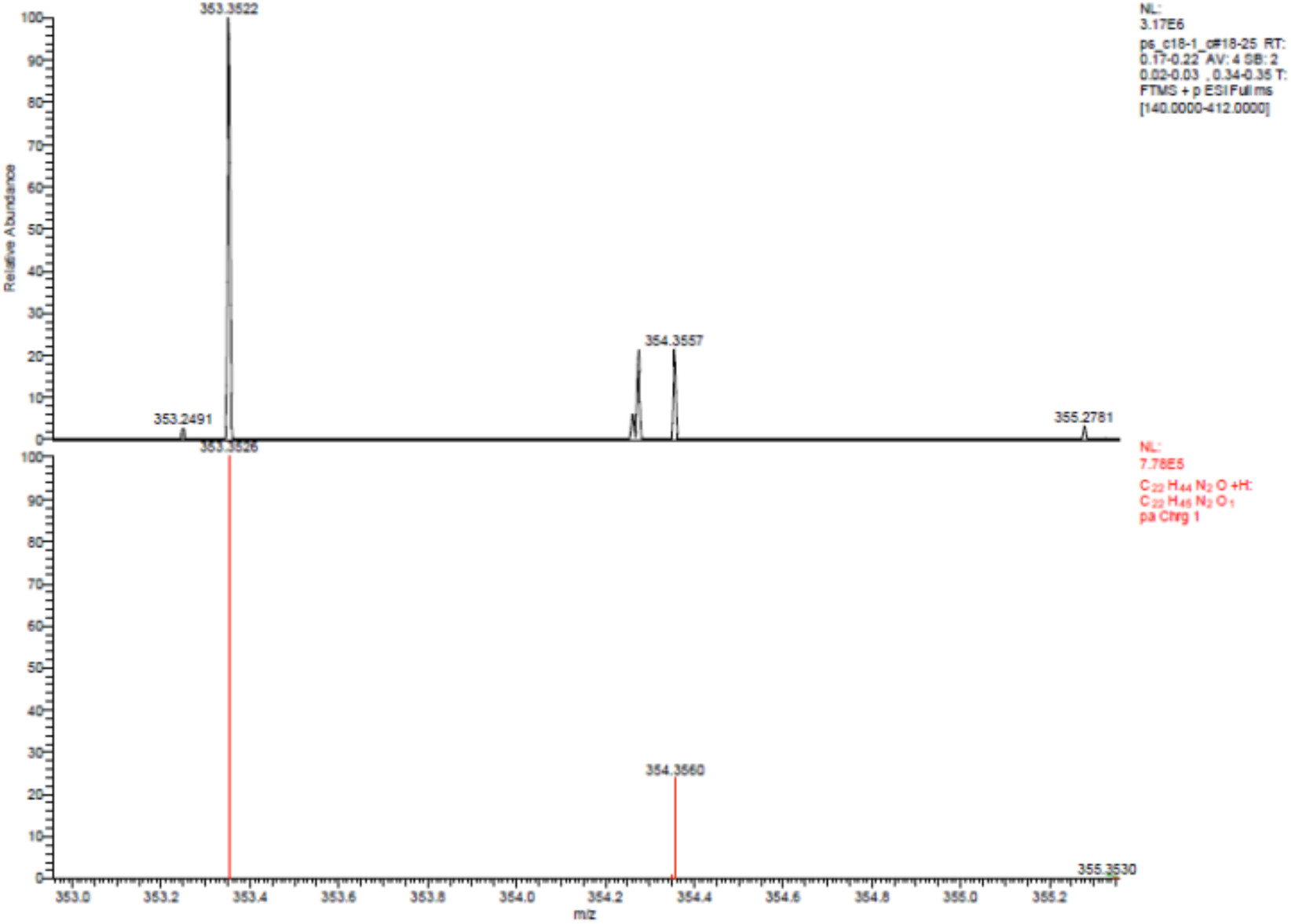
HRESIMS spectrum of compound **5**.

**Supplemental Data 26:**
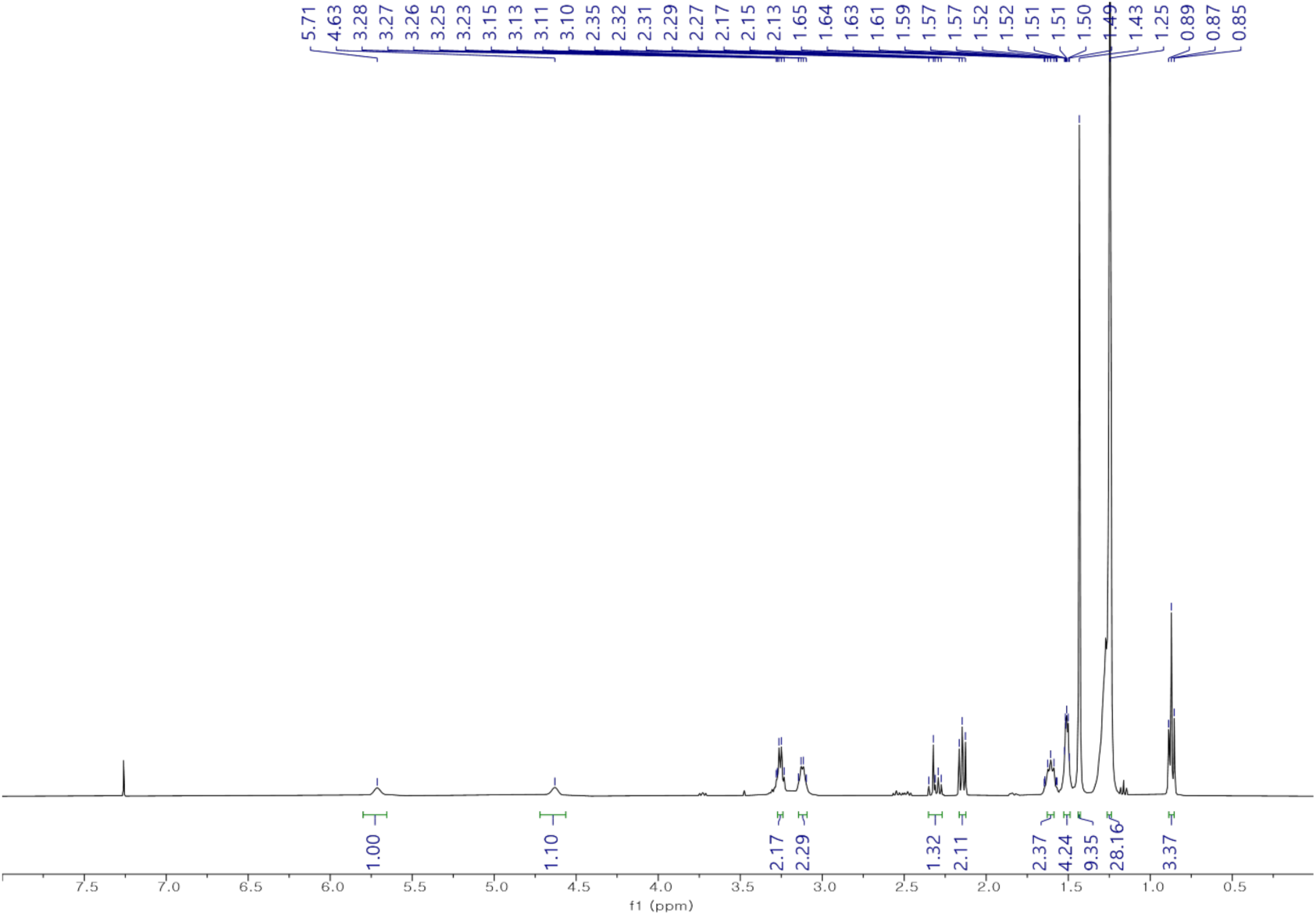
^1^H NMR spectrum of compound **6** in CDCl_3_ (600 MHz).

**Supplemental Data 27:**
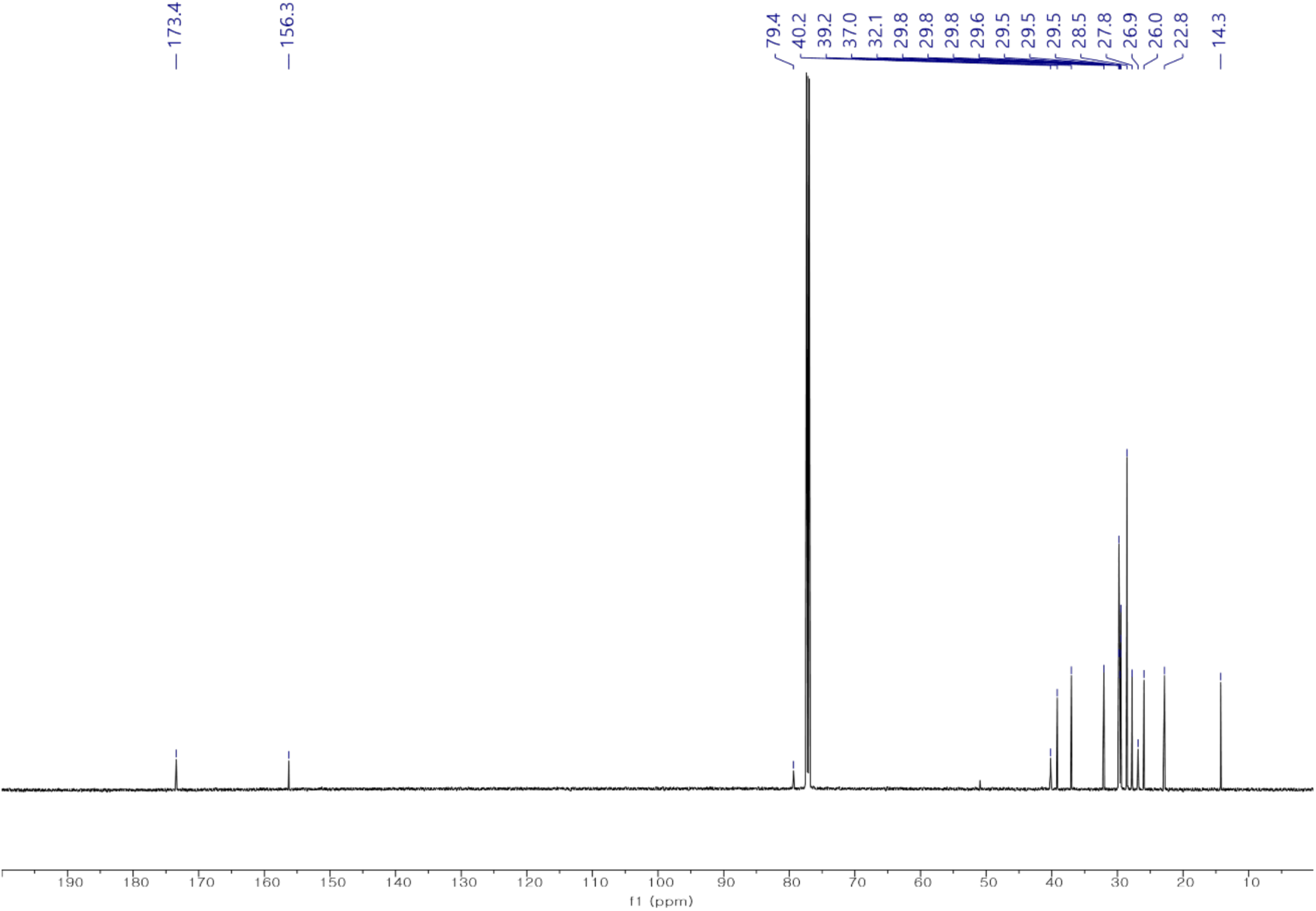
^13^C NMR spectrum of compound **6a** in CDCl_3_ (150 MHz).

**Supplemental Data 28:**
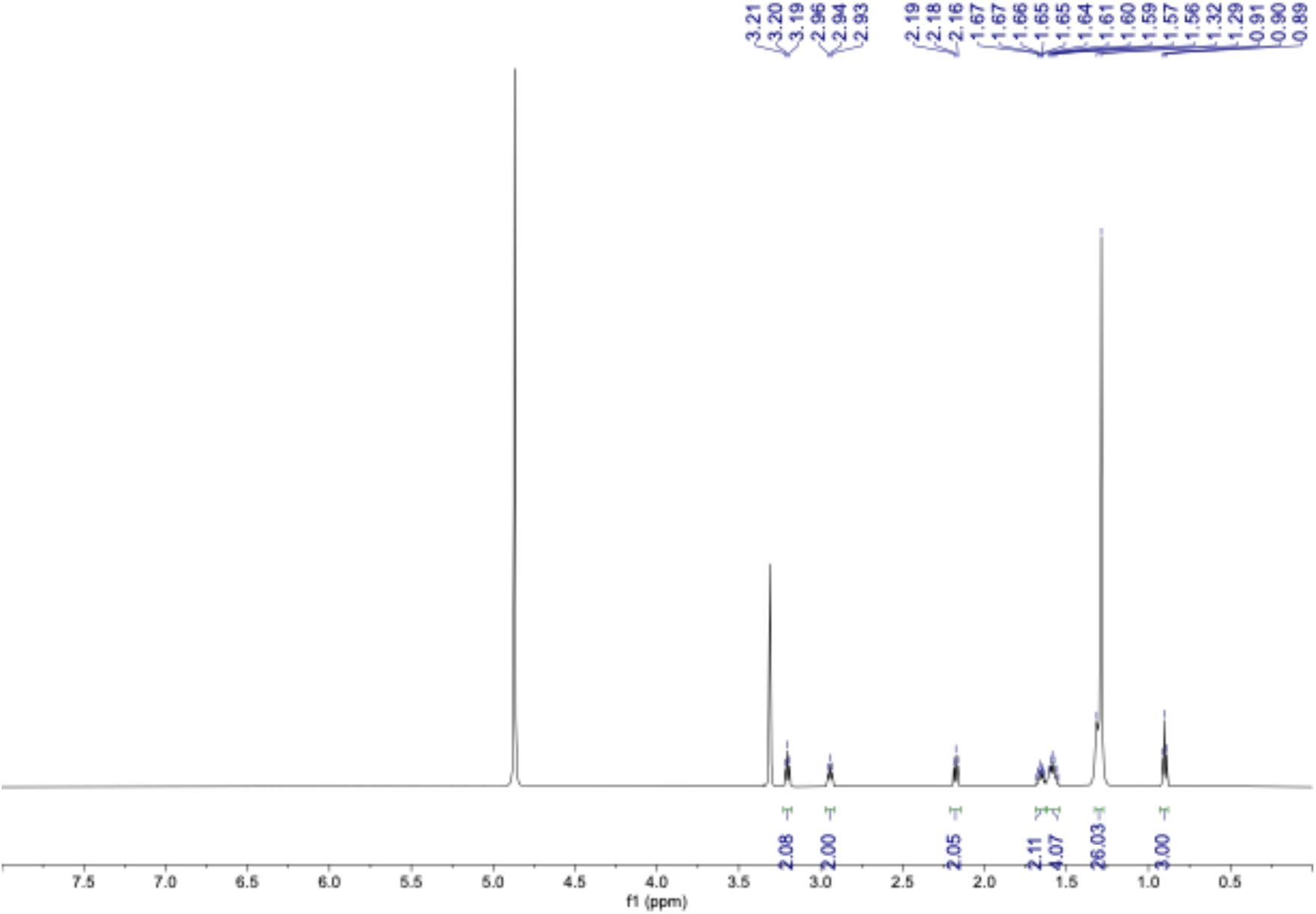
^1^H NMR spectrum of compound **6** in CD_3_OD (600 MHz).

**Supplemental Data 29:**
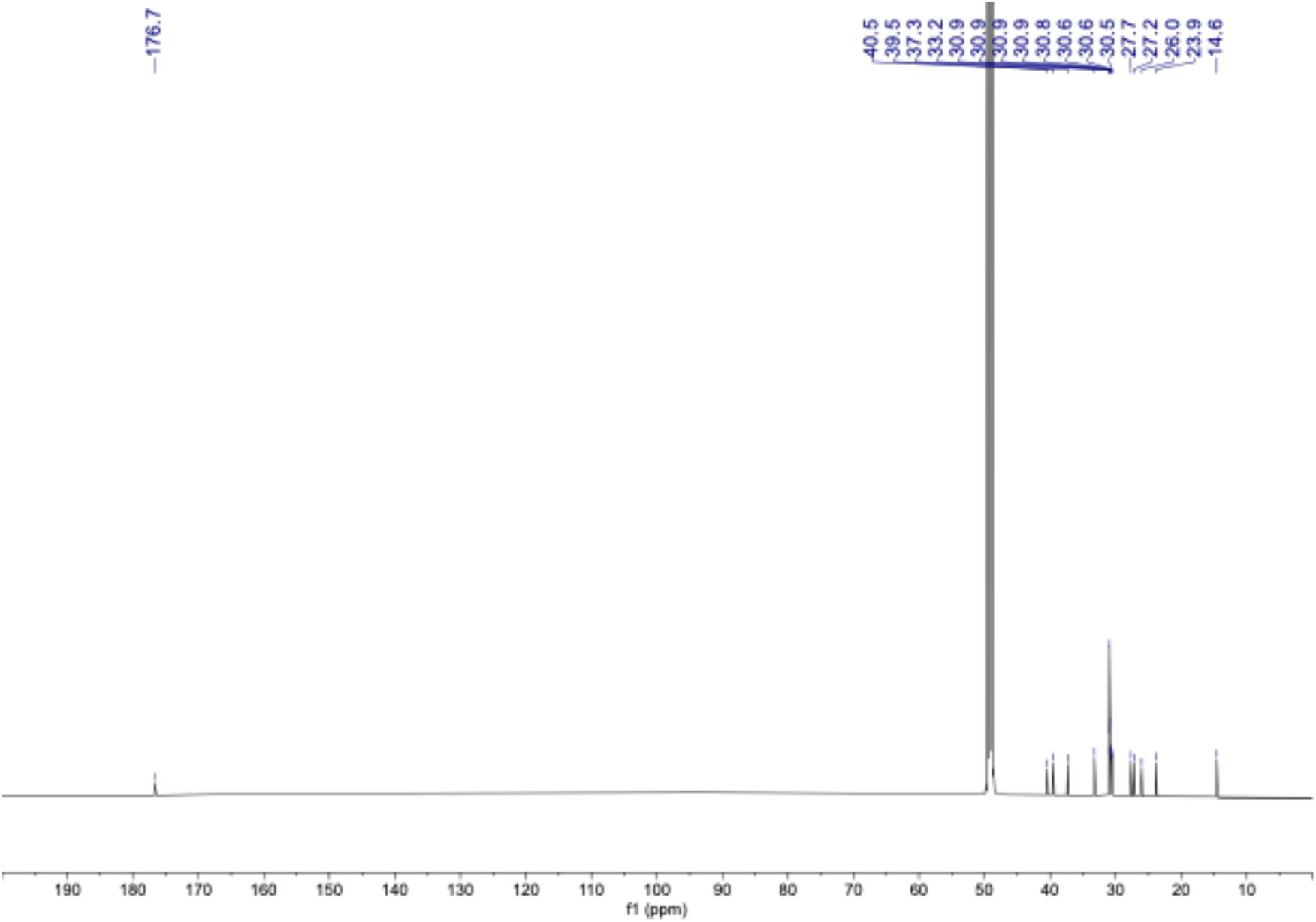
^13^C NMR spectrum of compound **6** in CD_3_OD 150 MHz.

**Supplemental Data 30:**
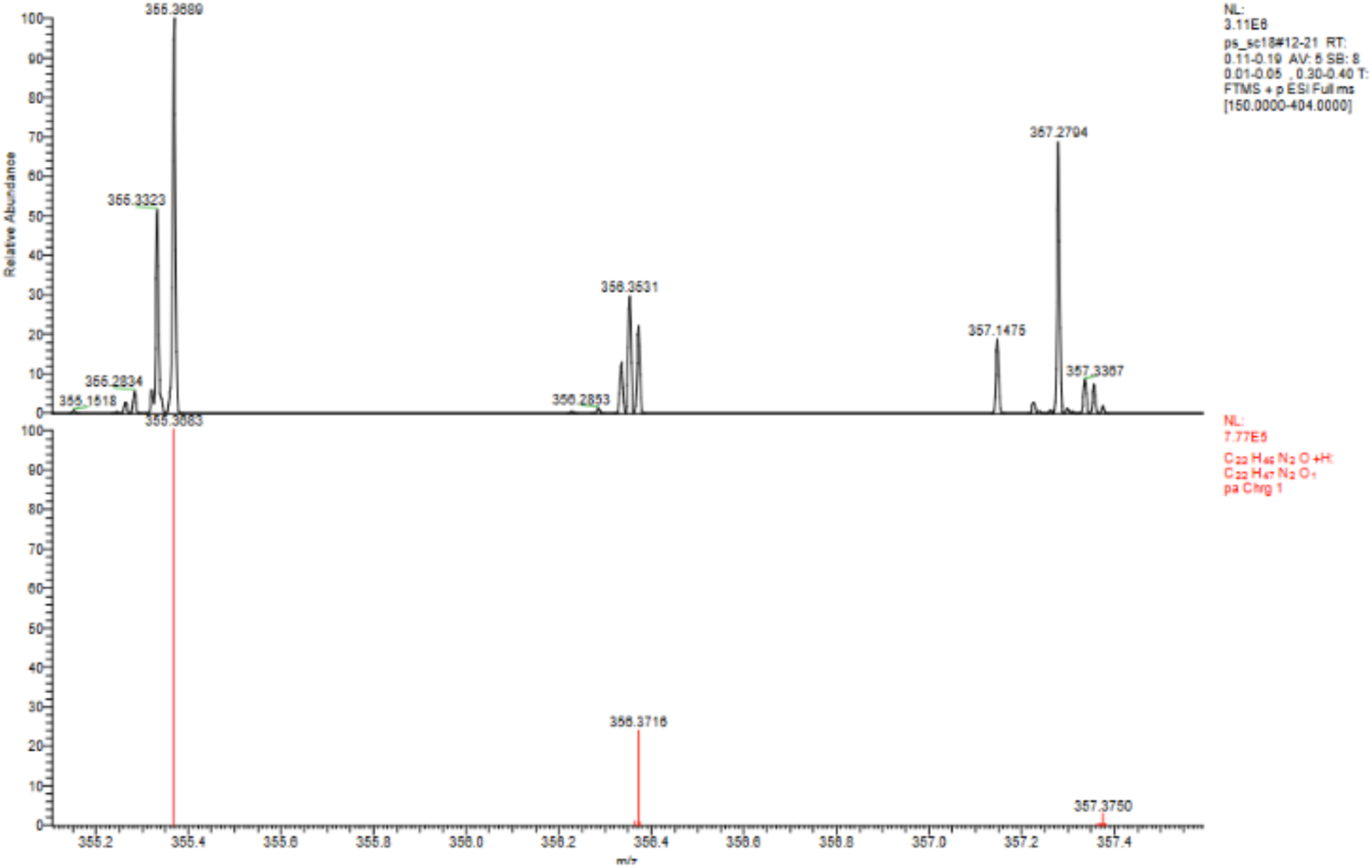
HRESIMS spectrum of compound **6**.

##### ^1^H NMR spectra data of *N*-oleoylputrescine hydrochloride synthetic standard generated in BioDuro LLC

**Supplemental Data 31.**
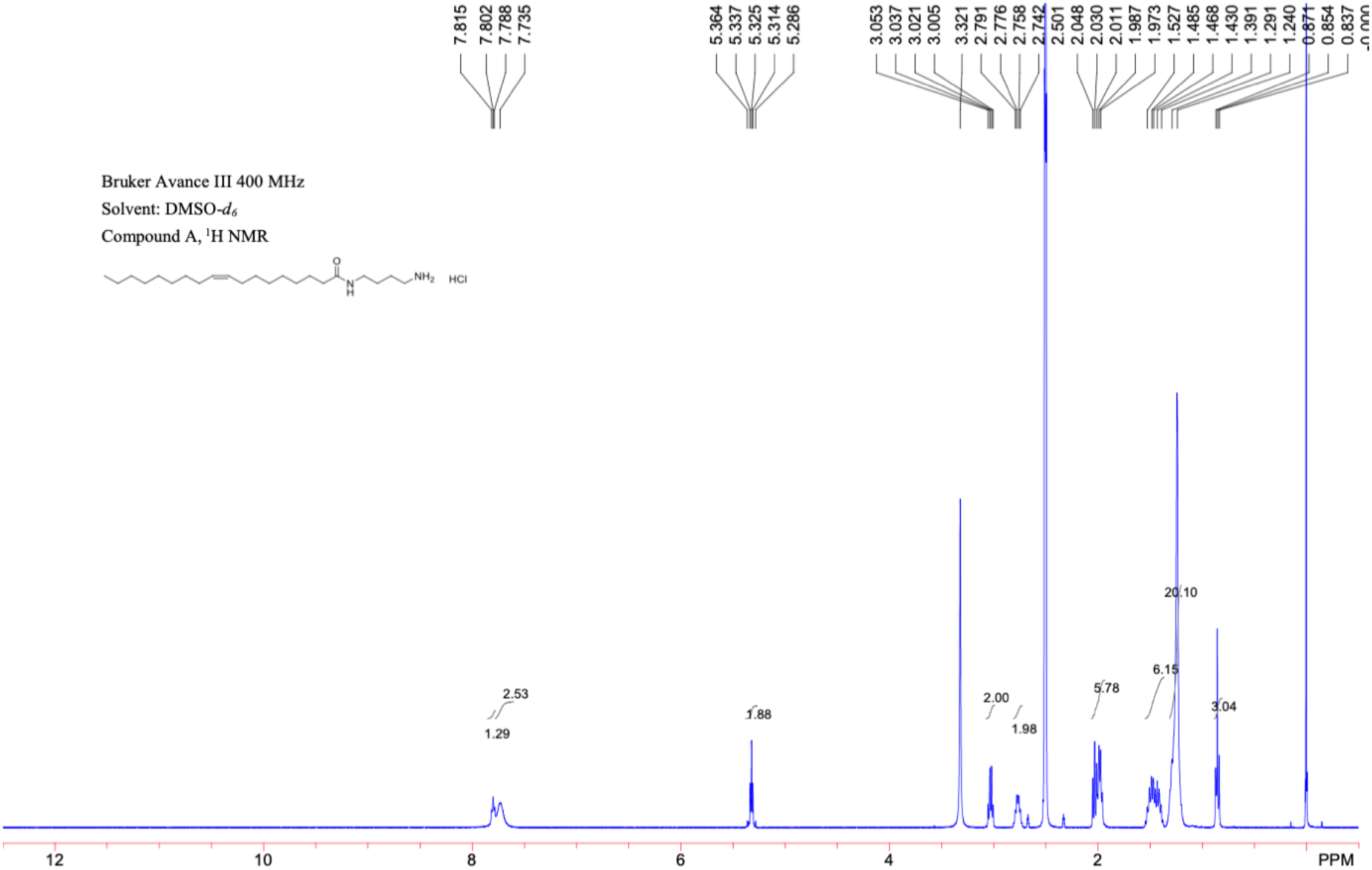
1^H^ NMR spectrum (400 MHz, DMSO) of *N*-oleoylputrescine hydrochloride

**Supplemental Data 32:**
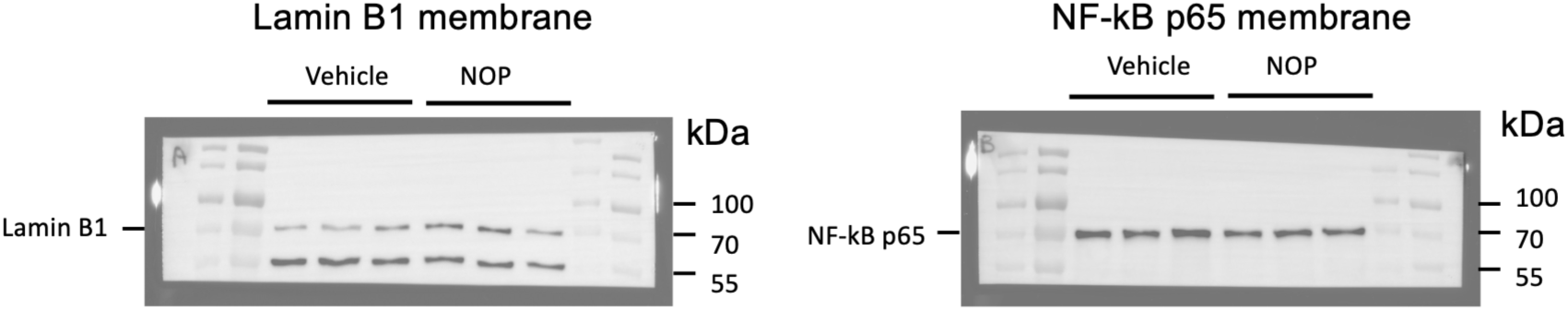
Uncropped Western blot images corresponding to Data 5h

## REFERENCES

1. Krautkramer, K.A., Fan, J., and Bäckhed, F. (2021). Gut microbial metabolites as multi-kingdom intermediates. Nat. Rev. Microbiol. 19, 77–94. 10.1038/s41579-020-0438-4.

2. Lavelle, A., and Sokol, H. (2020). Gut microbiota-derived metabolites as key actors in inflammatory bowel disease. Nat. Rev. Gastroenterol. Hepatol. 17, 223–237. 10.1038/s41575-019-0258-z.

3. Rooks, M.G., and Garrett, W.S. (2016). Gut microbiota, metabolites and host immunity. Nat. Rev. Immunol. 16, 341–352. 10.1038/nri.2016.42.

4. Blander, J.M., Longman, R.S., Iliev, I.D., Sonnenberg, G.F., and Artis, D. (2017). Regulation of inflammation by microbiota interactions with the host. Nat. Immunol. 18, 851–860. 10.1038/ni.3780.

5. Spivak, I., Fluhr, L., and Elinav, E. (2022). Local and systemic effects of microbiome-derived metabolites. EMBO Rep. 23, e55664. 10.15252/embr.202255664.

6. Iliev, I.D., Ananthakrishnan, A.N., and Guo, C.-J. (2025). Microbiota in inflammatory bowel disease: mechanisms of disease and therapeutic opportunities. Nat. Rev. Microbiol. 23, 509–524. 10.1038/s41579-025-01163-0.

7. Bhosle, A., Bae, S., Zhang, Y., Chun, E., Avila-Pacheco, J., Geistlinger, L., Pishchany, G., Glickman, J.N., Michaud, M., Waldron, L., et al. (2024). Integrated annotation prioritizes metabolites with bioactivity in inflammatory bowel disease. Mol. Syst. Biol. 20, 338–361. 10.1038/s44320-024-00027-8.

8. Franzosa, E.A., Sirota-Madi, A., Avila-Pacheco, J., Fornelos, N., Haiser, H.J., Reinker, S., Vatanen, T., Hall, A.B., Mallick, H., McIver, L.J., et al. (2019). Gut microbiome structure and metabolic activity in inflammatory bowel disease. Nat. Microbiol. 4, 293–305. 10.1038/s41564-018-0306-4.

9. Lloyd-Price, J., Arze, C., Ananthakrishnan, A.N., Schirmer, M., Avila-Pacheco, J., Poon, T.W., Andrews, E., Ajami, N.J., Bonham, K.S., Brislawn, C.J., et al. (2019). Multi-omics of the gut microbial ecosystem in inflammatory bowel diseases. Nature 569, 655–662. 10.1038/s41586-019-1237-9.

10. Elmassry, M.M., Sugihara, K., Chankhamjon, P., Kim, Y., Camacho, F.R., Wang, S., Sugimoto, Y., Chatterjee, S., Chen, L.A., Kamada, N., et al. (2025). A meta-analysis of the gut microbiome in inflammatory bowel disease patients identifies disease-associated small molecules. Cell Host Microbe 33, 218–234.e12. 10.1016/j.chom.2025.01.002.

11. Elmarsafawi, A.G., Hesterberg, R.S., Fernandez, M.R., Yang, C., Darville, L.N., Liu, M., Koomen, J.M., Phanstiel, O., Atkins, R., Mullinax, J.E., et al. (2023). Modulating the polyamine/hypusine axis controls generation of CD8+ tissue-resident memory T cells. JCI Insight 8. 10.1172/jci.insight.169308.

12. Fritsch, S.D., Sukhbaatar, N., Gonzales, K., Sahu, A., Tran, L., Vogel, A., Mazic, M., Wilson, J.L., Forisch, S., Mayr, H., et al. (2023). Metabolic support by macrophages sustains colonic epithelial homeostasis. Cell Metab. 35, 1931–1943.e8. 10.1016/j.cmet.2023.09.010.

13. Latour, Y.L., Gobert, A.P., and Wilson, K.T. (2020). The role of polyamines in the regulation of macrophage polarization and function. Amino Acids 52, 151–160. 10.1007/s00726-019-02719-0.

14. Nakamura, A., Kurihara, S., Takahashi, D., Ohashi, W., Nakamura, Y., Kimura, S., Onuki, M., Kume, A., Sasazawa, Y., Furusawa, Y., et al. (2021). Symbiotic polyamine metabolism regulates epithelial proliferation and macrophage differentiation in the colon. Nat. Commun. 12, 2105. 10.1038/s41467-021-22212-1.

15. Puleston, D.J., Baixauli, F., Sanin, D.E., Edwards-Hicks, J., Villa, M., Kabat, A.M., Kamiński, M.M., Stanckzak, M., Weiss, H.J., Grzes, K.M., et al. (2021). Polyamine metabolism is a central determinant of helper T cell lineage fidelity. Cell 184, 4186–4202.e20. 10.1016/j.cell.2021.06.007.

16. Susskind, B.M., and Chandrasekaran, J. (1987). Inhibition of cytolytic T lymphocyte maturation with ornithine, arginine, and putrescine. J. Immunol. 139, 905–912.

17. Wagner, A., Wang, C., Fessler, J., DeTomaso, D., Avila-Pacheco, J., Kaminski, J., Zaghouani, S., Christian, E., Thakore, P., Schellhaass, B., et al. (2021). Metabolic modeling of single Th17 cells reveals regulators of autoimmunity. Cell 184, 4168–4185.e21. 10.1016/j.cell.2021.05.045.

18. Yurdagul, A., Subramanian, M., Wang, X., Crown, S.B., Ilkayeva, O.R., Darville, L., Kolluru, G.K., Rymond, C.C., Gerlach, B.D., Zheng, Z., et al. (2020). Macrophage Metabolism of Apoptotic Cell-Derived Arginine Promotes Continual Efferocytosis and Resolution of Injury. Cell Metab. 31, 518–533.e10. 10.1016/j.cmet.2020.01.001.

19. Chang, F.-Y., Siuti, P., Laurent, S., Williams, T., Glassey, E., Sailer, A.W., Gordon, D.B., Hemmerle, H., and Voigt, C.A. (2021). Gut-inhabiting Clostridia build human GPCR ligands by conjugating neurotransmitters with diet- and human-derived fatty acids. Nat. Microbiol. 6, 792– 805. 10.1038/s41564-021-00887-y.

20. Cho, W., York, A.G., Wang, R., Wyche, T.P., Piizzi, G., Flavell, R.A., and Crawford, J.M. (2022). N-Acyl Amides from Neisseria meningitidis and Their Role in Sphingosine Receptor Signaling. Chembiochem 23, e202200490. 10.1002/cbic.202200490.

21. Tanes, C., Bittinger, K., Gao, Y., Friedman, E.S., Nessel, L., Paladhi, U.R., Chau, L., Panfen, E., Fischbach, M.A., Braun, J., et al. (2021). Role of dietary fiber in the recovery of the human gut microbiome and its metabolome. Cell Host Microbe 29, 394–407.e5. 10.1016/j.chom.2020.12.012.

22. Atiya Ali, M., Poortvliet, E., Strömberg, R., and Yngve, A. (2011). Polyamines in foods: development of a food database. Food Nutr. Res. 55. 10.3402/fnr.v55i0.5572.

23. Muñoz-Esparza, N.C., Latorre-Moratalla, M.L., Comas-Basté, O., Toro-Funes, N., Veciana-Nogués, M.T., and Vidal-Carou, M.C. (2019). Polyamines in Food. Front. Nutr. 6, 108. 10.3389/fnut.2019.00108.

24. Ye, C., Ho, D.J., Neri, M., Yang, C., Kulkarni, T., Randhawa, R., Henault, M., Mostacci, N., Farmer, P., Renner, S., et al. (2018). DRUG-seq for miniaturized high-throughput transcriptome profiling in drug discovery. Nat. Commun. 9, 4307. 10.1038/s41467-018-06500-x.

25. Coombes, J.L., and Powrie, F. (2008). Dendritic cells in intestinal immune regulation. Nat. Rev. Immunol. 8, 435–446. 10.1038/nri2335.

26. Mellman, I., and Steinman, R.M. (2001). Dendritic cells: specialized and regulated antigen processing machines. Cell 106, 255–258. 10.1016/s0092-8674(01)00449-4.

27. Kaiko, G.E., Ryu, S.H., Koues, O.I., Collins, P.L., Solnica-Krezel, L., Pearce, E.J., Pearce, E.L., Oltz, E.M., and Stappenbeck, T.S. (2016). The Colonic Crypt Protects Stem Cells from Microbiota-Derived Metabolites. Cell 165, 1708–1720. 10.1016/j.cell.2016.05.018.

28. Zhang, Y., Tu, S., Shao, X., Meng, J., Zhang, Z., Wei, W., Jing, X., Qin, Z., Wu, J., Xu, W., et al. (2026). Microbiota-derived IPA protects against colitis by regulating intestinal HMGCS2-mediated ketogenesis to facilitate mucosal healing. Nat. Commun. 17. 10.1038/s41467-026-69341-z.

29. Atreya, I., Atreya, R., and Neurath, M.F. (2008). NF-kappaB in inflammatory bowel disease. J. Intern. Med. 263, 591–596. 10.1111/j.1365-2796.2008.01953.x.

30. McDaniel, D.K., Eden, K., Ringel, V.M., and Allen, I.C. (2016). Emerging Roles for Noncanonical NF-κB Signaling in the Modulation of Inflammatory Bowel Disease Pathobiology. Inflamm. Bowel Dis. 22, 2265–2279. 10.1097/MIB.0000000000000858.

31. Mukherjee, T., Kumar, N., Chawla, M., Philpott, D.J., and Basak, S. (2024). The NF-κB signaling system in the immunopathogenesis of inflammatory bowel disease. Sci. Signal. 17, eadh1641. 10.1126/scisignal.adh1641.

32. Kong, L., Pokatayev, V., Lefkovith, A., Carter, G.T., Creasey, E.A., Krishna, C., Subramanian, S., Kochar, B., Ashenberg, O., Lau, H., et al. (2023). The landscape of immune dysregulation in Crohn’s disease revealed through single-cell transcriptomic profiling in the ileum and colon. Immunity 56, 444–458.e5. 10.1016/j.immuni.2023.01.002.

33. Smillie, C.S., Biton, M., Ordovas-Montanes, J., Sullivan, K.M., Burgin, G., Graham, D.B., Herbst, R.H., Rogel, N., Slyper, M., Waldman, J., et al. (2019). Intra- and Inter-cellular Rewiring of the Human Colon during Ulcerative Colitis. Cell 178, 714–730.e22. 10.1016/j.cell.2019.06.029.

34. Thomas, T., Friedrich, M., Rich-Griffin, C., Pohin, M., Agarwal, D., Pakpoor, J., Lee, C., Tandon, R., Rendek, A., Aschenbrenner, D., et al. (2024). A longitudinal single-cell atlas of anti-tumour necrosis factor treatment in inflammatory bowel disease. Nat. Immunol. 25, 2152–2165. 10.1038/s41590-024-01994-8.

35. Friedrich, M., Pohin, M., Jackson, M.A., Korsunsky, I., Bullers, S.J., Rue-Albrecht, K., Christoforidou, Z., Sathananthan, D., Thomas, T., Ravindran, R., et al. (2021). IL-1-driven stromal-neutrophil interactions define a subset of patients with inflammatory bowel disease that does not respond to therapies. Nat. Med. 27, 1970–1981. 10.1038/s41591-021-01520-5.

36. Arias-Salvatierra, D., Silbergeld, E.K., Acosta-Saavedra, L.C., and Calderon-Aranda, E.S. (2011). Role of nitric oxide produced by iNOS through NF-κB pathway in migration of cerebellar granule neurons induced by Lipopolysaccharide. Cell. Signal. 23, 425–435. 10.1016/j.cellsig.2010.10.017.

37. Jia, J., Liu, Y., Zhang, X., Liu, X., and Qi, J. (2013). Regulation of iNOS expression by NF-κB in human lens epithelial cells treated with high levels of glucose. Invest. Ophthalmol. Vis. Sci. 54, 5070–5077. 10.1167/iovs.13-11796.

38. Imam, T., Park, S., Kaplan, M.H., and Olson, M.R. (2018). Effector T Helper Cell Subsets in Inflammatory Bowel Diseases. Front. Immunol. 9, 1212. 10.3389/fimmu.2018.01212.

39. Scott, M.C., Steier, Z., Pierson, M.J., Stolley, J.M., O’Flanagan, S.D., Soerens, A.G., Wijeyesinghe, S.P., Beura, L.K., Dileepan, G., Burbach, B.J., et al. (2025). Deep profiling deconstructs features associated with memory CD8+ T cell tissue residence. Immunity 58, 162–181.e10. 10.1016/j.immuni.2024.11.007.

40. Wang, Z., Wang, Z., Wang, J., Diao, Y., Qian, X., and Zhu, N. (2016). T-bet-Expressing B Cells Are Positively Associated with Crohn’s Disease Activity and Support Th1 Inflammation. DNA Cell Biol. 35, 628–635. 10.1089/dna.2016.3304.

41. Hao, Z., and Rajewsky, K. (2001). Homeostasis of peripheral B cells in the absence of B cell influx from the bone marrow. J. Exp. Med. 194, 1151–1164. 10.1084/jem.194.8.1151.

42. Hitchcock, D.S., Krejci, J.N., Sturgeon, C.E., Dennis, C.A., Jeanfavre, S.T., Avila-Pacheco, J.R., and Clish, C.B. (2025). Eclipse: a Python package for alignment of two or more nontargeted LC-MS metabolomics datasets. Bioinformatics 41. 10.1093/bioinformatics/btaf290.

43. Dührkop, K., Fleischauer, M., Ludwig, M., Aksenov, A.A., Melnik, A. V, Meusel, M., Dorrestein, P.C., Rousu, J., and Böcker, S. (2019). SIRIUS 4: a rapid tool for turning tandem mass spectra into metabolite structure information. Nat. Methods 16, 299–302. 10.1038/s41592-019-0344-8.

44. Cho, W., York, A.G., Wang, R., Wyche, T.P., Piizzi, G., Flavell, R.A., and Crawford, J.M. (2022). N-Acyl Amides from Neisseria meningitidis and Their Role in Sphingosine Receptor Signaling. Chembiochem 23, e202200490. 10.1002/cbic.202200490.

45. Chang, F.-Y., Siuti, P., Laurent, S., Williams, T., Glassey, E., Sailer, A.W., Gordon, D.B., Hemmerle, H., and Voigt, C.A. (2021). Gut-inhabiting Clostridia build human GPCR ligands by conjugating neurotransmitters with diet- and human-derived fatty acids. Nat. Microbiol. 6, 792– 805. 10.1038/s41564-021-00887-y.

46. Thompson, L.R., Sanders, J.G., McDonald, D., Amir, A., Ladau, J., Locey, K.J., Prill, R.J., Tripathi, A., Gibbons, S.M., Ackermann, G., et al. (2017). A communal catalogue reveals Earth’s multiscale microbial diversity. Nature 551, 457–463. 10.1038/nature24621.

47. Bolyen, E., Rideout, J.R., Dillon, M.R., Bokulich, N.A., Abnet, C.C., Al-Ghalith, G.A., Alexander, H., Alm, E.J., Arumugam, M., Asnicar, F., et al. (2019). Reproducible, interactive, scalable and extensible microbiome data science using QIIME 2. Nat. Biotechnol. 37, 852–857. 10.1038/s41587-019-0209-9.

48. Callahan, B.J., McMurdie, P.J., Rosen, M.J., Han, A.W., Johnson, A.J.A., and Holmes, S.P. (2016). DADA2: High-resolution sample inference from Illumina amplicon data. Nat. Methods 13, 581–583. 10.1038/nmeth.3869.

49. Quast, C., Pruesse, E., Yilmaz, P., Gerken, J., Schweer, T., Yarza, P., Peplies, J., and Glöckner, F.O. (2012). The SILVA ribosomal RNA gene database project: improved data processing and web-based tools. Nucleic Acids Res. 41, D590–D596. 10.1093/nar/gks1219.

50. Mallick, H., Rahnavard, A., McIver, L.J., Ma, S., Zhang, Y., Nguyen, L.H., Tickle, T.L., Weingart, G., Ren, B., Schwager, E.H., et al. (2021). Multivariable association discovery in population-scale meta-omics studies. PLoS Comput. Biol. 17, e1009442. 10.1371/journal.pcbi.1009442.

51. Goto, N., Goto, S., Imada, S., Hosseini, S., Deshpande, V., and Yilmaz, Ö.H. (2022). Lymphatics and fibroblasts support intestinal stem cells in homeostasis and injury. Cell Stem Cell 29, 1246–1261.e6. 10.1016/j.stem.2022.06.013.

52. Miyoshi, H., and Stappenbeck, T.S. (2013). In vitro expansion and genetic modification of gastrointestinal stem cells in spheroid culture. Nat. Protoc. 8, 2471–2482. 10.1038/nprot.2013.153.

53. Paik, D., Yao, L., Zhang, Y., Bae, S., D’Agostino, G.D., Zhang, M., Kim, E., Franzosa, E.A., Avila-Pacheco, J., Bisanz, J.E., et al. (2022). Human gut bacteria produce ΤΗ17-modulating bile acid metabolites. Nature 603, 907–912. 10.1038/s41586-022-04480-z.

54. O’Connell, D.J., Kolde, R., Sooknah, M., Graham, D.B., Sundberg, T.B., Latorre, I., Mikkelsen, T.S., and Xavier, R.J. (2016). Simultaneous Pathway Activity Inference and Gene Expression Analysis Using RNA Sequencing. Cell Syst. 2, 323–334. 10.1016/j.cels.2016.04.011.

55. Picelli, S., Faridani, O.R., Björklund, A.K., Winberg, G., Sagasser, S., and Sandberg, R. (2014). Full-length RNA-seq from single cells using Smart-seq2. Nat. Protoc. 9, 171–181. 10.1038/nprot.2014.006.

56. Andersson, D., Kebede, F.T., Escobar, M., Österlund, T., and Ståhlberg, A. (2024). Principles of digital sequencing using unique molecular identifiers. Mol. Aspects Med. 96, 101253.

57. Li, H., and Durbin, R. (2009). Fast and accurate short read alignment with Burrows–Wheeler transform. bioinformatics 25, 1754–1760.

58. Sherman, B.T., Hao, M., Qiu, J., Jiao, X., Baseler, M.W., Lane, H.C., Imamichi, T., and Chang, W. (2022). DAVID: a web server for functional enrichment analysis and functional annotation of gene lists (2021 update). Nucleic Acids Res. 50, W216–W221. 10.1093/nar/gkac194.

59. Liberzon, A., Birger, C., Thorvaldsdóttir, H., Ghandi, M., Mesirov, J.P., and Tamayo, P. (2015). The Molecular Signatures Database (MSigDB) hallmark gene set collection. Cell Syst. 1, 417–425. 10.1016/j.cels.2015.12.004.

60. Korotkevich, G., Sukhov, V., Budin, N., Shpak, B., Artyomov, M.N., and Sergushichev, A. (2016). Fast gene set enrichment analysis. Preprint, 10.1101/060012 https://doi.org/10.1101/060012.

61. Korotkevich, G., Sukhov, V., Budin, N., Shpak, B., Artyomov, M.N., and Sergushichev, A. (2016). Fast gene set enrichment analysis. Preprint, 10.1101/060012 https://doi.org/10.1101/060012.

62. Liberzon, A., Birger, C., Thorvaldsdóttir, H., Ghandi, M., Mesirov, J.P., and Tamayo, P. (2015). The Molecular Signatures Database (MSigDB) hallmark gene set collection. Cell Syst. 1, 417–425. 10.1016/j.cels.2015.12.004.

63. Hao, Y., Hao, S., Andersen-Nissen, E., Mauck, W.M., Zheng, S., Butler, A., Lee, M.J., Wilk, A.J., Darby, C., Zager, M., et al. (2021). Integrated analysis of multimodal single-cell data. Cell 184, 3573–3587.e29. 10.1016/j.cell.2021.04.048.

64. Finak, G., McDavid, A., Yajima, M., Deng, J., Gersuk, V., Shalek, A.K., Slichter, C.K., Miller, H.W., McElrath, M.J., Prlic, M., et al. (2015). MAST: a flexible statistical framework for assessing transcriptional changes and characterizing heterogeneity in single-cell RNA sequencing data. Genome Biol. 16, 278. 10.1186/s13059-015-0844-5.

65. Love, M.I., Huber, W., and Anders, S. (2014). Moderated estimation of fold change and dispersion for RNA-seq data with DESeq2. Genome Biol. 15, 550. 10.1186/s13059-014-0550-8.

66. Borcherding, N., Vishwakarma, A., Voigt, A.P., Bellizzi, A., Kaplan, J., Nepple, K., Salem, A.K., Jenkins, R.W., Zakharia, Y., and Zhang, W. (2021). Mapping the immune environment in clear cell renal carcinoma by single-cell genomics. Commun. Biol. 4, 122. 10.1038/s42003-020-01625-6.

67. Chun, E., Lavoie, S., Fonseca-Pereira, D., Bae, S., Michaud, M., Hoveyda, H.R., Fraser, G.L., Gallini Comeau, C.A., Glickman, J.N., Fuller, M.H., et al. (2019). Metabolite-Sensing Receptor Ffar2 Regulates Colonic Group 3 Innate Lymphoid Cells and Gut Immunity. Immunity 51, 871–884.e6. 10.1016/j.immuni.2019.09.014.

